# Evaluating the definition and distribution of spring ephemeral wildflowers in eastern North America

**DOI:** 10.1101/2023.10.04.560873

**Authors:** Abby J. Yancy, Benjamin R. Lee, Sara E. Kuebbing, Howard S. Neufeld, Michelle Elise Spicer, J. Mason Heberling

## Abstract

The herbaceous layer accounts for the majority of plant biodiversity in eastern North American forests, encompassing substantial variation in life history strategy and function. One group of early season herbaceous understory species, colloquially referred to as “spring ephemeral wildflowers,” are of particular ecological and cultural importance. Despite this, little is known about the prevalence and biogeographic patterns of the spring ephemeral strategy. Here, we used georeferenced and dated observations from the Global Biodiversity Information Facility (GBIF) to define the phenological strategies of 559 herbaceous, vascular, understory plant species in eastern North America from a composite species list encompassing 16 site-level species lists ranging from Georgia to southern Canada. Specifically, we estimated activity periods from regional observations (primarily consisting of citizen scientist iNaturalist observations) and classified species as ephemeral if they completed all aboveground activity (including leafing, flowering, fruiting, and senescence) prior to an estimated date of canopy closure derived from remote-sensed data. We then evaluated the richness of these species at the landscape scale using estimates of biogeographic and environmental drivers aggregated for 100 km x 100 km grid cells. Importantly, our spatially-explicit approach defines each species’ spring ephemerality along a continuous scale (which we call the Ephemerality Index, EI) based on the proportion of its range in which it senesces before canopy closure (with EI = 0 indicating a species that is never ephemeral and EI = 1 indicating a species that is always ephemeral). We found that 18.4% (103 species) of understory wildflowers exhibited spring ephemerality in at least part of their range, with only 3.4% of all species exhibiting ephemeral behavior in all parts of their range. Ephemeral species had higher overall richness and composed a higher proportion of understory biodiversity in low-elevation areas with intermediate spring temperatures and elevated spring precipitation. Spring ephemerals peaked in both absolute species richness and relative proportion at mid latitudes. These biogeographic patterns deserve further study in other regions of the world and to uncover mechanisms behind these patterns. Using our new metric, our results demonstrate that the spring ephemeral strategy is not a discrete category, but rather a continuum that can vary across species’ ranges.

## Introduction

Understory herbaceous species comprise approximately 80% of the total plant biodiversity in eastern North American deciduous temperate forests (Gilliam 2007; Spicer et al. 2020). They provide important ecosystem services and functions including early season floral resources and contributions to nutrient cycling in soils (Muller and Bormann, 1976; Gilliam 2007). A subset of these species, commonly referred to as spring ephemeral wildflowers, are only active in spring, relying solely on elevated light availability while canopy trees are leafless to facilitate their annual energy budgets (Uemura 1994; Neufeld and Young 2014). The duration of favorable growing conditions before tree canopy closure is variable across latitude, where the duration decreases with increasing latitude (Neufeld and Young 2014). Due to this difference, it is speculated that the diversity of spring active plants should be higher in southeastern North American deciduous forests, where the spring ephemeral strategy is thought to be most advantageous (Routhier and Lapointe 2002). Despite growing attention from researchers, the spring ephemeral strategy is not well defined in scientific literature and, as a result, relatively little is known about the distribution of this phenological strategy across North America.

Light availability is a critical limiting resource for understory plants in all forests with closed canopy environments. Understory light levels are dependent not only on canopy openness (e.g., the proportion of light intercepted by canopy tree leaves), but also on the canopy tree phenology (i.e., the timing and duration of when canopy trees have leaves). The amount of light reaching the forest floor decreases with increasing canopy leaf area index and varies with cyclical annual ‘phenoseasons’ in temperate deciduous forests (Hutchinson and Matt 1977), the dominant forest type across eastern North America. This includes the availability of sunflecks, which are more abundant before full canopy leaf expansion, but can provide critical light to summer-green and evergreen understory plant species (Way and Pearcy 2012). In these systems, light is therefore most limiting starting with the onset of canopy leaf out in late spring, lasting through the summer until canopy leaf senescence in autumn as the trees return to dormancy for the winter. As a result, understory plant species have adapted a wide variety of different strategies to survive and persist in such a highly variable environment.

Plants growing in these forest understories must either be able to tolerate or avoid shade in order to assimilate enough carbon from photosynthesis needed for survival, reproduction, and growth of new tissue (Hull 2002; Valladares and Niinemets 2008). Shade tolerators, such as most summer-blooming wildflowers (*sensu* Neufeld and Young 2014), generally employ photosynthetic strategies comprised of low maximum photosynthetic rates, sacrificing peak performance in direct sunlight for improved respirational efficiency, and low light compensation points by maintaining positive carbon assimilation rates even at low light levels (Hull 2002; Valladares and Niinemets 2008). In contrast, shade avoiders typically maintain high maximum photosynthetic rates while altering their growing season activity to maximize their access to high light availability (Lapointe 2001), thereby reducing or eliminating their need for better efficiency in shady conditions. Importantly, temperate understory plant species commonly employ some combination of both strategies by maximizing photosynthetic rates in early spring and then downregulating their photosynthetic machinery once the canopy closes above them (Rothstein and Zak 2001; Bauerle et al. 2012; Peltier and Ibáñez 2015; Heberling et al. 2019).

Many shade avoiders in temperate deciduous forests fall into the category of “spring ephemeral,” meaning that they rely solely on access to spring light availability to assimilate carbon before retreating back to belowground dormancy as the canopy closes (Neufeld and Young 2014). Most of these species create large belowground rooting structures to store carbon (Lubbers and Lechowicz 1989; Lapointe and Lerat 2006; Gandin et al. 2011). These storage structures facilitate summer dormancy and belowground growth during the autumn and winter, until emergence the following spring (Lapointe and Lerat 2006). Access to spring light is important for the carbon budgets of all spring-active plants (Heberling et al. 2019, Lee and Ibáñez 2021a,b), but spring carbon gain represents 100% of spring ephemeral carbon assimilation each year.

The term “spring ephemeral” is used colloquially in botanical literature and there is little consensus about the exact definition of the term or which species constitute “true” spring ephemerals. For example, while many authors utilize the term to describe species which meet the strict definition of ephemerality we employ here (i.e., the tendency of a species to employ the spring ephemeral strategy, defined as spring-active plants that complete all aboveground activity, including leafing, flowering, fruiting, and senescence, prior to the timing of full canopy closure), other authors equate the term to spring blooming species generally, even if the given species retains leaves for weeks or months after canopy closure. Although one possibility for this inconsistency is a lack of strict definitions for this life history strategy, it is also possible that some species might differ in their ephemerality over the extent of their range owing to phenotypic plasticity, ecotypic variation, or complex interactions with local environmental cues. This is supported by evidence showing different phenological sensitivities (i.e., the magnitude of change in phenology over a driver like spring temperature or time) for this group of species across large geographical gradients (Miller et al. 2022, Alecrim et al. 2022), as well as smaller scale, natural variations in canopy closure (Dion et al. 2017). Specifically, Miller et al. (2022) and Alecrim et al. (2022) found that access to light had different relationships with spring temperature depending on what part of the temperate deciduous biome was considered (northern, central, or southern). In contrast, Dion et al. (2017) found that *Allium tricoccum*, a spring-active wildflower, delayed leaf senescence under tree canopies with later leaf out phenology, compared to those under canopies with earlier closure, resulting in higher biological success. These findings suggest that plants have adapted different responsiveness to environmental cues in different parts of their ranges, which could theoretically lead to differences in overall phenological strategy.

Here, we adopt a strict definition of spring ephemerality where species are defined as such if their aboveground activity (including leafing, flowering, fruiting, and senescence) is entirely limited to the period of high light availability that occurs prior to canopy leaf out. Using this definition, we combined remote-sensed imagery and observational and herbarium collections data aggregated from the Global Biodiversity Information Facility (GBIF) to estimate spring ephemerality in forest herbaceous understory plants across eastern North America. Our goals were 1) to develop a new, continuous metric of spring ephemerality allowing for spatial variation in ephemerality across species ranges, and 2) to quantify how spring ephemeral species richness varies across the landscape. Importantly, our study proposes a solution to some of the debate over what defines spring ephemerality, providing important context for future research of this highly diverse group of plants and their ecology.

## Methods

We divided our analysis into two parts, consistent with the goals described above. In Goal I, we combined regional species checklists with observations of plant activity aggregated from GBIF (https://www.gbif.org/; accessed between March 12-13, 2023) and remote-sensed MODIS green-up data to define spring ephemerality by species and region (i.e., grid cell, described below). In Goal II, we used this information to explore the geographic distribution of the spring ephemeral strategy.

### Goal I: Defining spring ephemerality

To meet our first goal, we acquired 16 site-level floristic species checklists collected in forested natural areas across eastern North America and representative of regional species pools (Table S1, Fig. 1A). Species lists were distributed across the eastern North American deciduous forest biome with sites from as far south as Congaree National Park in South Carolina, northwest as far as Huron Mountain Club on the upper peninsula of Michigan, and as far northeast as Laurence Mauricie National Park in Quebec. A full list of site names, abbreviations, and reference citations is contained in Table S1. We consolidated the site-level lists into a combined species list incorporating all species from all sites (n = 1666 species) and used the USDA PLANTS Database (https://plants.usda.gov/home) to identify each species’ taxonomy and growth habit. We retained all herbaceous vascular species (excluding woody plants, grasses, sedges, and ferns). This resulted in 834 species encompassing 81 taxonomic families and 329 genera. We then excluded non-forest species based on published habitat descriptions (Weakley 2022), aggregated cultivars and subspecies to the species level, and updated species names to account for synonymy using the *taxize* package in R (Chamberlain and Szöcs 2013). We removed species that did not have enough observations to conduct a full statistical analysis (i.e., fewer than 10 observations per grid cell, see below), resulting in a final species list of 559 species (see Supplemental Data).

**Fig. 1:**
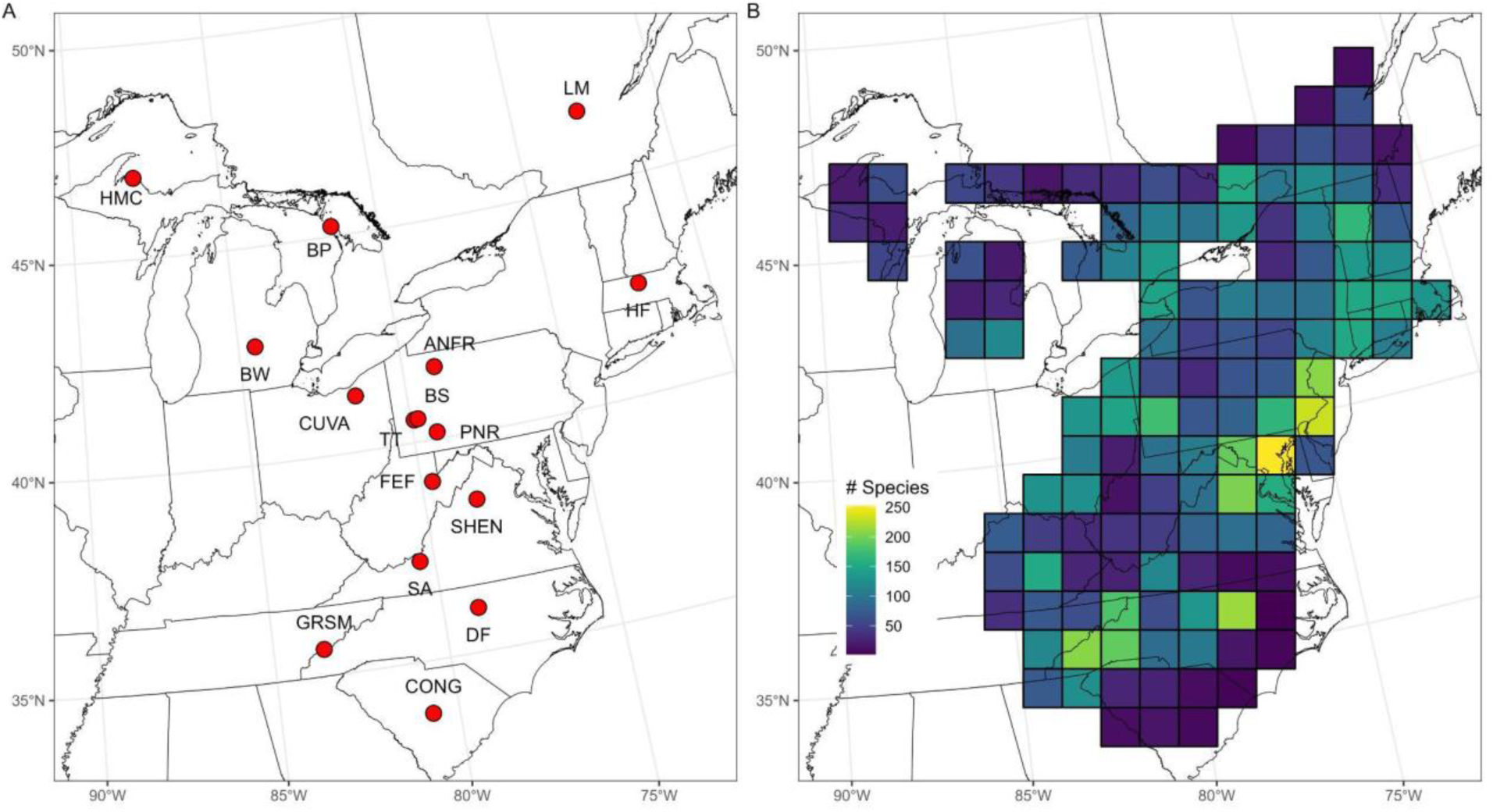
A) Map of the 16 sites for which species lists were used to compile the comprehensive species list used in this study. Site abbreviations and descriptions of site-level species lists can be found in Table S1. B) Map of the 128 equal-area grid cells used to extract GBIF observations of plant activity, remote-sensed green up and environmental data. Cell color indicates the estimated understory plant richness in each cell.

To define spring ephemerality, we needed information about 1) the phenology of the understory plants in our species list and 2) the relative timing of tree canopy closure for each of the 16 sites. For the former, we followed the general approach described in Belitz et al. (2023), where we created an equal area grid consisting of 100 km x 100 km grid cells with which to extract phenological observations and evaluate ecological variation. First, we extracted all research-grade species observations from the Global Biodiversity Information Facility (GBIF, data downloaded March 12-13, 2023) for between 2015-2021 using the *rgbif* package in R (Chamberlain et al. 2023; filtered list of observations available at https://doi.org/10.15468/dd.zyywht). Observations from GBIF were primarily composed of community-science data (Fig. S1; 98.5% of total observations). The majority of these observations were collected by users of iNaturalist (https://www.inaturalist.org/; 86.7% of total observations), a community-science platform where users upload images of plants, animals, and other biota along with metadata about where and when the image was collected. With the help of artificial intelligence software and an extensive user-base, users also identify each organism to species where possible, with “research-grade” observations being those where more than two-thirds of identifiers agree on a taxon. For species that did not return any observations from the GBIF search, we extracted observations directly from iNaturalist using the *rinat* package in R (Barve and Hart 2022). A full list of data sources is provided in the Supplemental Data. For each species, we removed duplicate observations (same species observed in the same location on the same day and by the same user) to avoid pseudoreplication of observations and we removed all observations assigned to the first of a month due to a previously-described artifact where observations that are missing exact day data are occasionally automatically and erroneously assigned to the first of the month (Belitz et al. 2023).

Spring phenology varies temporally and spatially, with evidence of variable sensitivity to spring temperatures across large geographic gradients (e.g., Kharouba and Vellend 2015, Alecrim et al. 2022, Miller et al. 2022, Gallinat et al. 2021). To account for this variability, and to assess whether spring ephemerality within a given species varies across space, we constructed a grid across eastern North America consisting of equal-area 100 km x 100 km grid cells bounded by 25 to 47.88 °N and −97.00 to −52.02 °W, for an initial total of 1,089 cells that encompassed all 16 sites where species lists were compiled. We then collated species-level observations in each cell using location data accompanying each observation and removed cells with no observations (e.g., those in the middle of the Atlantic Ocean).

Previous research has shown that estimates of activity period or growing season length can be strongly biased by the number of observations used to make the estimations (Pearse et al. 2017; Belitz et al. 2020b). Specifically, estimates of active periods are more accurate when they are made on the basis of more observations. However, observation-based biases have been documented in iNaturalist and other crowd-sourced datasets (Belitz et al. 2020b), such as higher observer effort in populated areas. As such, we adopted criteria that are considered best practice for assessing spatial variation in phenological activity (Belitz et al. 2023). Namely, we only considered a species within a given grid cell if there were more than 10 recorded observations of that species in that cell in our dataset, a threshold that was previously shown to allow accurate estimation of active period duration (Belitz et al. 2020b). Further, we used the *phenesse* R package (Belitz et al. 2020a) to estimate the 99th percentile of activity period for each species and grid cell, a measure considered to be equivalent to the end of the observed active period (Belitz et al. 2020b), therefore representing an estimate of early summer senescence in spring ephemeral wildflowers.

For each species present in a given cell, regardless of number of observations, we estimated activity periods from 500 iterations of bootstrapped, randomized distributions of observational data. In short, this approach (detailed fully in Belitz et al. 2020b) allowed us to statistically account for differences in observation effort that might otherwise skew our estimates of active season length. We chose the 99th percentile estimator in this analysis because it was the most conservative of all thresholds that we considered (95th and 99th percentiles of raw data and 95th and 99th percentiles of *phenesse* results; Fig. S2). Furthermore, although threshold choice changed species-level estimates of ephemerality index, it did not significantly affect the overall signal in our final analysis (Fig. S2), indicating that the modeling approach was robust to this choice in statistical approximation .

Next, to ensure that the species in our combined list were representative of the geographic scope of grid cells used in analysis, we narrowed the cells used in our analysis by removing those that were further than 100 km from one of the 16 sites with species lists and that were not between any two sites. We further removed grid cells if over half of their area was covered by water (primarily associated with cells in the Great Lakes and coastal Atlantic Ocean regions), or if estimates of canopy closure were overly biased by agricultural green-up (primarily associated with the U.S. Midwest region and the Ontario peninsula; see next paragraph). Figure S2 provides a graphical description of which of the original 1,089 cells were or were not included as well as reasoning for excluding cells from the broader analysis. Cumulatively, this grid cell thinning resulted in a working dataset comprising 642,526 observations of 559 herbaceous, non-woody forest understory species from 128 100 km x 100 km grid cells ranging from northern Georgia to northern Wisconsin and southern Quebec (Fig. 1B).

Lastly, because our definition of spring ephemerality depends not only on the understory phenology, but also on the phenology of overstory canopy trees, we estimated cell-level canopy close phenology using MCD12Q2 Enhanced Vegetation Index (EVI) green-up data collected by MODIS (Gray et al. 2019). Specifically, we extracted the “maturity” parameter, which corresponds to the day of the year when a pixel (500 m spatial resolution) reaches 90% of its peak green-up value each year. This metric has high fidelity to ground-truthed measures of canopy development and start of spring (Peng et al. 2017), and should be interpreted as a relatively conservative proxy for the beginning of the summer shady period in deciduous forests. We averaged pixel EVI values within each grid cell across the same years used to collate GBIF and iNaturalist observations (2015-2021), excluding pixels associated with impervious surfaces and water cover. We then estimated the day of the year of canopy closure as the median pixel value within each cell. As briefly described above, we removed any cells where canopy closure was estimated to occur after day 181 (∼July 1st), which indicated a strong skew from agricultural green up associated with summer crops like corn and soybean (e.g., Wardlow et al. 2007) in August and September (Fig. S3).

Using the combined grid cell-level understory herbaceous phenology and estimated canopy close information, we designated every species in every cell where it was observed as either “spring ephemeral” or not, with ephemerals defined as species that were only active (i.e., observed or collected in the GBIF database) prior to the estimated day of canopy close. Importantly, this allowed species to be considered ephemeral in some cells while not being considered ephemeral in others (*e.g.*, a species could have the same temporal distribution of observations in all cells where it occurs, but the ephemerality designation could differ because of variation in the cell-level estimates of the day of canopy closure in that year). Species were thus assigned an overall ephemerality index (EI) value, calculated as the proportion of cells where it was designated ephemeral divided by the total number of cells in which it was observed. For example, *Dicentra cucullaria* was defined as ephemeral in 36 out of the 67 cells it was observed in for a EI value of 0.54. Species that were ephemeral in every cell in which they are observed would have an EI of 1, while those that were never considered ephemeral in any cell would have an EI of 0.

Following the assignment of EI values, we looked for species with categorizations that were likely erroneous, such as for those that we knew had an evergreen leaf habit but returned an EI value > 0. For each species with 0 > EI > 1, we looked for primary literature and field guides describing its leaf habit and, if it was described as being evergreen, we manually changed its EI to 0. This resulted in a total of 25 species being converted to an EI of 0 (Table S2). Figure S4 presents a graphical representation of relationships between estimated canopy close and estimated species-level end of season timing, specifically highlighting the 25 evergreen species that were misclassified according to our dataset. Importantly, we did not alter EI values for species described by field guides as deciduous, leaving open the possibility that, although we would not consider a species to be ephemeral based on our personal observations/experience, it is possible that ephemerality varies across space and that the EI metric is picking up on true biogeographical variation in this trait. A comprehensive list of species and their EI values can be found in the Supplemental Data.

Importantly, in order to conduct this analysis we made several statistical and theoretical assumptions that likely strongly shaped our results. These assumptions primarily concern how we quantify plant active periods (both for understory wildflowers and as it pertains to canopy close phenology) and how we filtered observations, species, and grid cells. For the sake of full transparency, Table S3 contains a list of assumptions (including those associated with our statistical analysis in Goal II) and the justifications for making them.

### Goal II: Environmental drivers of spring ephemerality

Our second goal was to explore environmental correlations with the grid cell-level richness of spring ephemeral species and the proportion of total species in each cell that are considered spring ephemerals. We therefore took the cell-level species lists with ephemerality definitions from Goal I and tallied the number of species that were defined as ephemeral in each cell (note that this is not the the EI value, rather it is the cell-level ones and zeros that are then averaged across all cells to calculate the EI value). We also calculated the proportion of species in each cell that were considered ephemeral out of the total richness of understory herbaceous species observed in that cell.

We used hierarchical Bayesian models to quantify correlations between these responding variables and various environmental and biogeographical drivers. First, we modeled the richness of ephemeral species in cell *i* using a Poisson likelihood distribution:

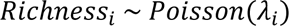

with mean and variance λ. This parameter was then log-transformed and modeled with common intercept *⍺* and linear relationships *β* associated with *n* fixed effects drivers:

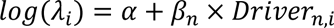

The final model included five fixed effects, all of which were averaged at the 100 km x 100 km cell level: squared April temperature (°C), May precipitation (mm), elevation (meters above sea level), human population density (people km^-2^), and total herbaceous species richness (including all species, not just spring ephemerals, with a maximum possible value of 582, the number of species evaluated in this study). Other drivers were considered in preliminary data exploration (including other monthly ranges of average temperature and precipitation), but we chose the five listed based on preliminary data exploration and *a priori* assumptions. The full rationale for choosing the suite of drivers is described in the Supplemental Methods.

We modeled the proportion of ephemeral species relative to total understory species richness in cell *i* using a normal likelihood distribution:

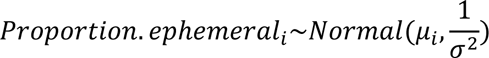

with mean *μ* and variance 1/σ^2^. The mean was modeled similarly to the first model, with common intercept *⍺* and linear relationships *β* associated with *n* fixed effects drivers:

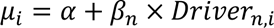

The proportion calculation already implicitly accounts for total species richness, so only the other four drivers were included in this model (squared April temperature, May precipitation, elevation, and human population density).

April temperature and May precipitation data were averaged from the same years in which herbaceous observation data were collected (2015-2021), accessed from WorldClim 2.1 (Fick & Hijmans 2017). Elevation was similarly averaged at the grid cell level using WorldClim data. We included human population density (based on 2020 census estimates and accessed via the NASA Socioeconomic Data and Applications Center; https://sedac.ciesin.columbia.edu/data/set/gpw-v4-population-density-rev11/data-download) as a proxy for bias in observer effort (*sensu* Li et al. 2019). In short, cells with higher human density are likely to have higher number of observations than cells with lower human density, which could bias iNaturalist sampling efforts.

Our estimates of activity period duration already account for some bias that might otherwise be associated with observer effort. That is, the bootstrapped estimation approach we used is meant to prevent species being incorrectly defined as ephemeral based on low sampling effort within a given grid cell. Still, greater observer effort could result in higher likelihood of observing more total species or rarer species, and so we account for it in our models using human population density as a fixed effect driver. All drivers were mean-centered prior to analysis with standard deviation of 0.5, following Gelman (2008) so that the relative influence of each driver could be directly compared. Copies of the raw and transformed data are freely available at [LINK TO BE PROVIDED AT PUBLICATION]. All parameters were assigned minimally informative prior distributions: *⍺*, *βn ∼ Normal*(0, 1000), *σ^2^* ∼ *Uniform*(0, 100). Models were run in *JAGS* (Plummer 2003) using the *R2jags* package (Su & Yajima 2021) in R v4.3.0. We used three MCMC chains per model and posterior distributions were assessed from 70,000 iterations following a 30,000 iteration warmup period. We report posterior estimated means with 95% Bayesian Credible Intervals (BCI), and describe results as statistically significant if the BCIs do not overlap zero.

## Results

### Species-level ephemerality

The 128 100 km x 100 km grid cells in this analysis contained an average of 82 forest herb species (± 57 s.d.; Fig. 1B), with a total number of 559 unique understory herbaceous species that we evaluated for spring ephemerality. Out of these species, 103 (18.4%) had an Ephemerality Index of >0 (Fig. 2), meaning that they were categorized as ephemeral in at least one of the grid cells in which they occurred. Nineteen of these species (3.4% of total) had an EI = 1 meaning they were ephemeral in every cell in which they occurred (e.g., *Scilla siberica* and *Euphorbia spathulata*, Fig. 3H&I), while 94 (16.8% of total) had 0 < EI < 1. The remaining 446 species (79.8%) had an EI = 0, meaning that they were never categorized as ephemeral in any cell that they occurred (e.g., *Antennaria virginica* and *Arisaema triphyllum*; Fig. 3A&B). Of these, 25 (6.2% of total) were species that were originally classified as having EI > 0 despite being defined as evergreen or semi-evergreen in the primary literature (average EI for these species before adjustment was 0.28 ± 0.26, meaning they had relatively low EI values to begin with). Note that these misclassifications were most likely the result of too few observations of these species in certain cells. For example, iNaturalist observations are known to be biased to heavier spring observation effort (partially as a result of the City Nature Challenge which typically occurs in May; Di Cecco et al. 2021), leading to the possibility that evergreen species may not be as commonly measured outside of the early growing season. Other research on herbarium collections found that botanists are biased toward collecting plants that are actively flowering (e.g., Panchen et al. 2019). We posit that a similar bias could be present in iNaturalist data where evergreen angiosperms are primarily observed during their flowering period despite being present in the forest year-round. Only six species (*Cardamine angustata, Krigia dandelion*, *Muscari botryoides*, *Narcissus pseudonarcissus*, *Scilla siberica* (Fig. 3I), and *Viola bicolor*) had an EI = 1 and were observed in more than two different cells, three of which (*Muscari botryoides*, *Narcissus pseudonarcissus*, and *Scilla siberica*) are introduced geophytes that are commonly cultivated.

**Fig. 2:**
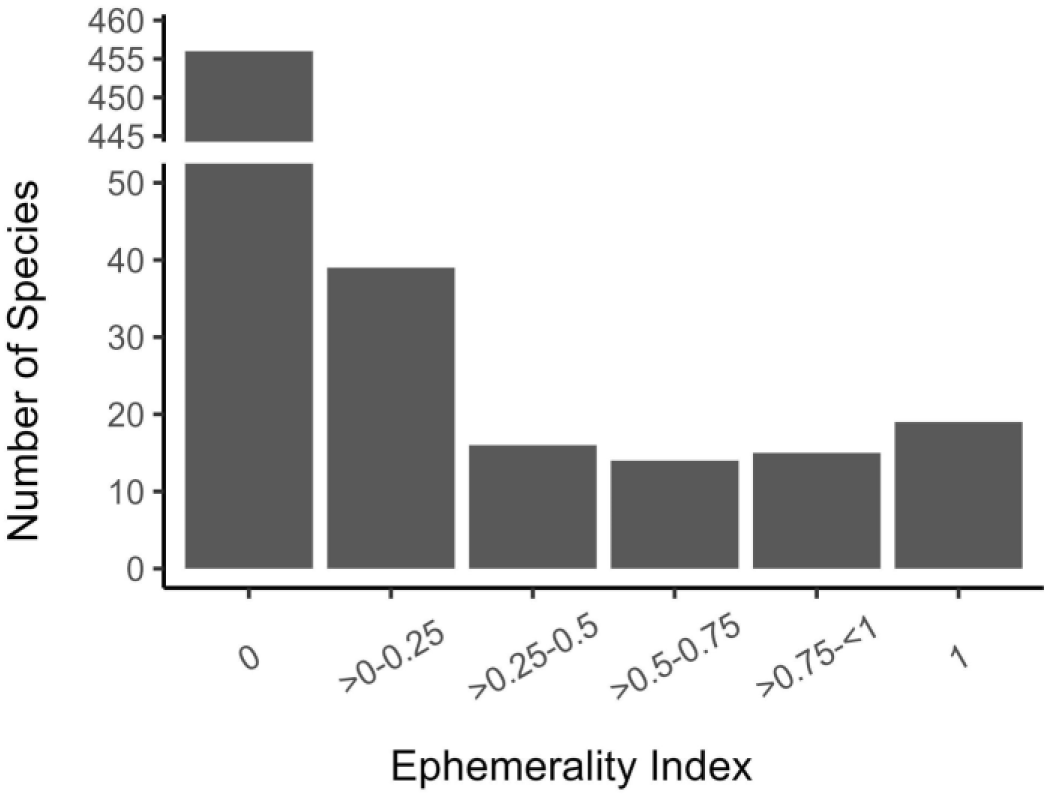
Distribution of ephemerality index (EI) values calculated among the 559 herbaceous understory plants in this study. EI values range from 0 (never ephemeral in any grid cell) to 1 (always ephemeral in every grid cell in which it occurs). There is a break in the y axis to better show the variation in the columns with EI > 0.

**Fig. 3:**
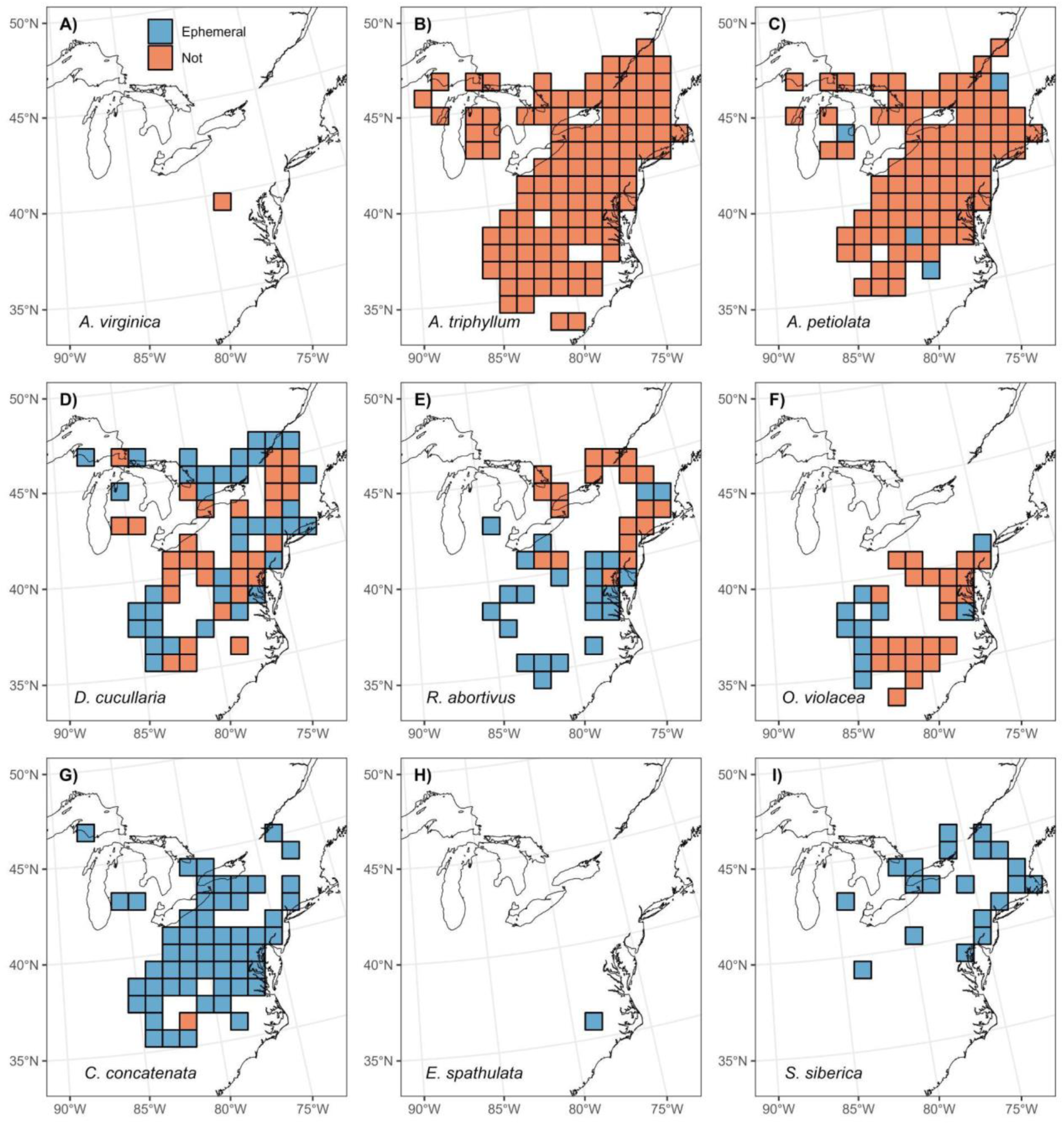
Maps showing examples of ephemerality classification for nine species of understory wildflower: (A) *Antennaria virginica*, (B) *Arisaema triphyllum*, (C) *Alliaria petiolata*, (D) *Dicentra cucullaria*, (E) *Ranunculus abortivus*, (F) *Oxalis violacea*, (G) *Cardamine concatenata*, (H) *Euphorbia spathulata*, and (I) *Scilla siberica*. In each panel, 100 km x 100 km grid cells are shaded blue if the species was defined as ephemeral or shaded red if it was not. Absent grid cells indicate that data for that species were not available in that location.

Out of the 94 species with EI values between 0 and 1, 30 of them had EI >= 0.5, thereby suggesting a high prevalence of spring ephemerality. For example, *Cardamine concatenata* was defined as ephemeral in 55 of the 56 cells it was observed in (EI = 0.98; Fig. 3G). Furthermore, despite there being 94 species defined as ephemeral in only part of their range (see Supplemental Materials), there was substantial variation in the geographic distribution of cells where species were defined as ephemeral. Some species, such as *Dicentra cucullaria* (EI = 0.54), had no immediately discernible pattern to the distribution of ephemerality across their observed range of cells, with ephemeral and non-ephemeral cells occurring in all parts of their ranges (Fig. 3D). Others showed distributions that appeared more directional. For example, *Ranunculus abortivus* (EI = 0.55, Fig. 3E) was primarily classified as ephemeral in the southern portion of its range, but not the north. Another example, *Oxalis violacea* (EI = 0.30, Fig. 3F), was primarily classified as ephemeral in the western portion of its range and as non-ephemeral in the east. Importantly, 75 out of the total 559 species (13.4%) only occurred in a single cell within our dataset, with 64 having EI = 0 (e.g., *Antennaria virginica*, Fig. 3A) and 11 having EI = 1 (e.g., *Euphorbia spathulata*, Fig. 3H). These species are likely rare or have highly-constrained ranges. Ephemerality index values for these species may thus be somewhat misleading given that they represent a binary value (ephemeral or not), rather than the continuous metric present in species with wide distributions.

### Spatial patterns of spring ephemerality

The richness of spring ephemeral wildflowers across the landscape was strongly associated with latitude, with peak ephemeral richness (Fig. 4A, B), peak total species richness (Fig. 4C,D), and peak proportion of ephemeral species (Fig. 4E,F) occurring at middle latitudes (approximately 40° N). Further, we found several statistically significant associations between ephemerality (richness and proportion) and the four drivers that we assessed in the hierarchical models (April temperature, May precipitation, elevation, and human population density). Ephemeral richness and the proportion of ephemeral species both peaked at intermediate April temperatures as signified with a statistically significant negative quadratic relationship (Fig. 5A,B). Both metrics were greatest at low elevations and at high May precipitation (both drivers had statistically significant relationships in both models). Human population density, a proxy for sampling effort (see Methods), was negatively associated with both richness and proportion of ephemeral species (although it was only statistically significant in the richness model, with 95% Bayesian credible intervals overlapping zero for the proportion ephemeral model). Lastly, the number of species was positively and significantly associated with ephemeral richness, indicating that grid cells with more species are more likely to have a greater number of ephemeral species.

**Fig. 4:**
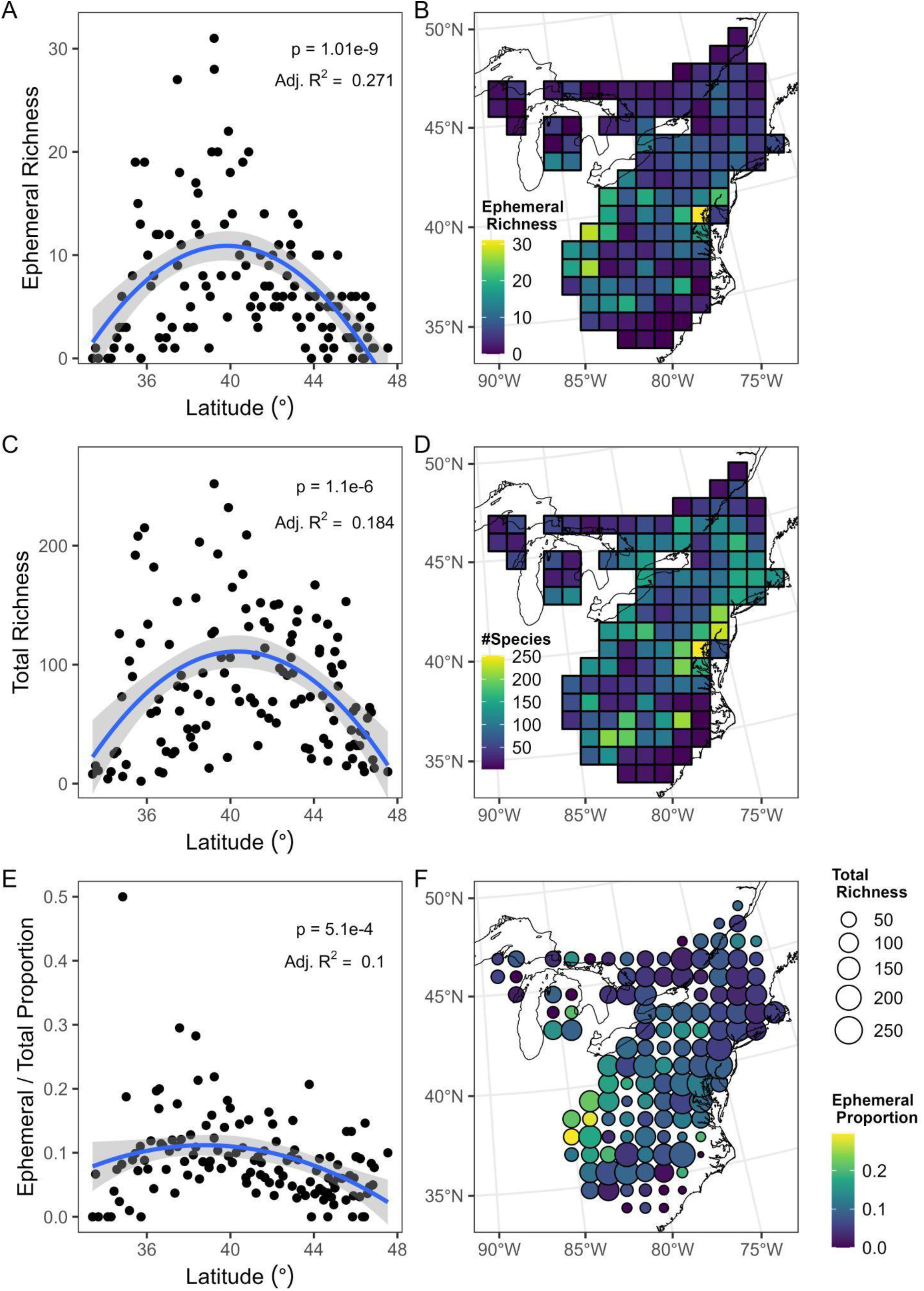
Left panels show quadratic relationships between latitude and (A) spring ephemeral richness, (C) total understory species richness, and (E) proportion of total species richness made up of spring ephemeral species. Panels (B) and (D) show maps of grid cells with fill color indicating (B) ephemeral richness and (D) total richness. Colored circles in panel (F) are centered on the corresponding grid cells with circle size indicating total species richness and color indicating proportion of total species that were defined as spring ephemeral. The cell in panel E with proportion of ephemerality = 0.5 (n = 4 species) was omitted from panel F to better represent variation across the rest of the grid. The unaltered version of panel F is provided as Fig. S5.

**Fig. 5:**
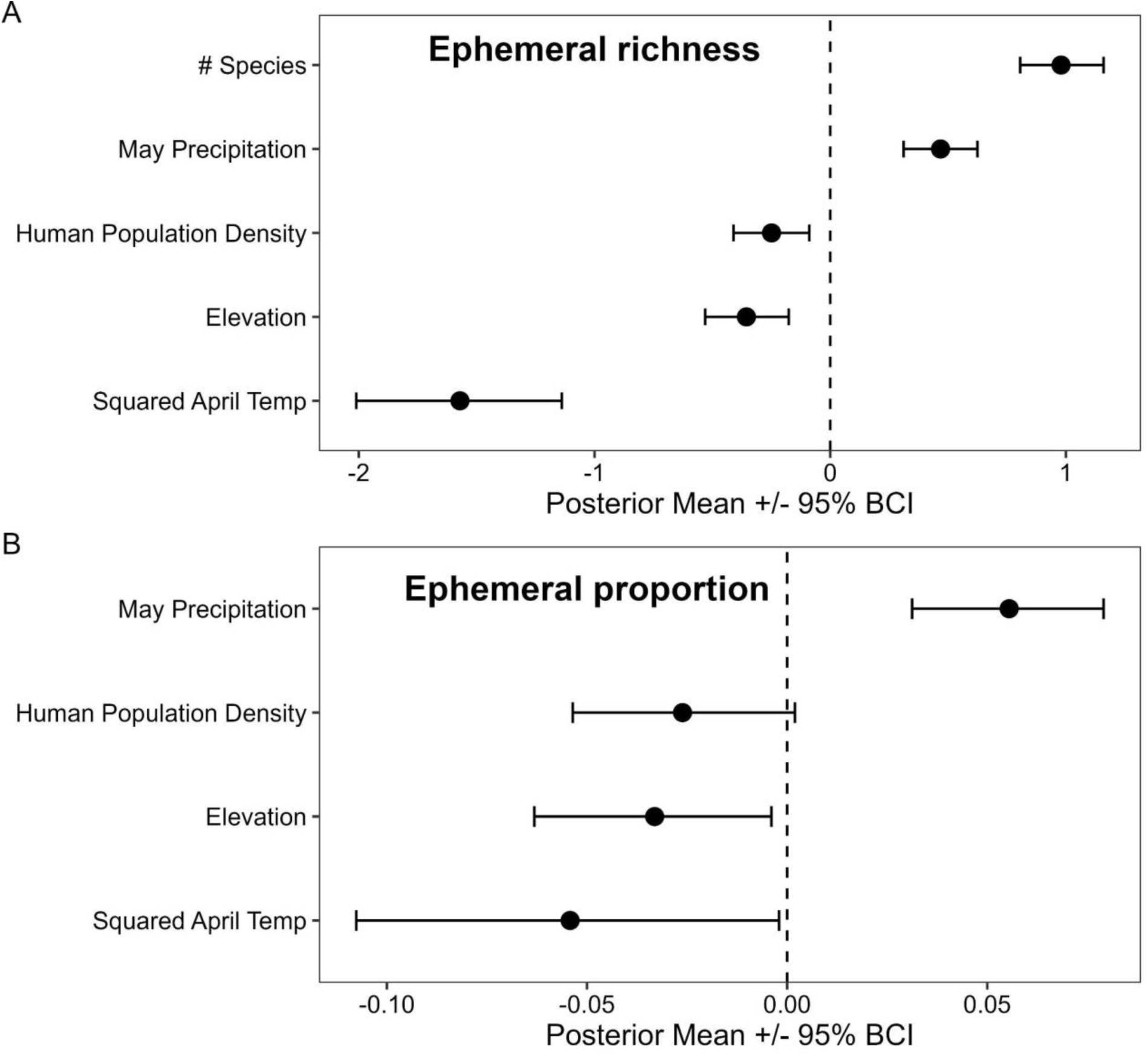
Posterior estimated means (points) and 95% Bayesian Credible Intervals (BCIs, whiskers) for driver fixed effects in the (A) ephemeral richness and (B) ephemeral proportion of total species models. Fixed effects are considered statistically significant if the BCIs do not overlap the dashed zero line.

April temperature was the strongest overall driver of spring ephemeral richness (i.e., the absolute value of the effect size mean was the greatest of all drivers), with temperature effect size ranging from 1.6 times greater (total species number) to 6.3 times greater (human population density) than the other drivers (Fig. 5A). May precipitation and squared April temperature were approximately equally as strong in the ephemeral proportion model (Fig. 5B), albeit with more homogenous effect sizes across all drivers (1.6 to 2.1 times greater effect sizes compared to elevation and human population density, respectively).

## Discussion

### Spring ephemerality is a common strategy but varies across space

A large proportion of North American forest understory vascular herbaceous species (18.4%) meet our strict definition of ‘spring ephemeral’ (completing their fruiting, flowering, and senescence before the canopy closes above them) in at least part of their range. This estimate is four times higher than estimated by Spicer et al. (2019), which defined “true ephemerals” as plants that bloomed between March and May and senesced before July 1 based upon descriptions in online flora databases and, importantly, not accounting for differences across species’ ranges. Spring ephemeral wildflower species were found across the entire extent of eastern deciduous forests covered in this study, with spring ephemeral life history strategies identified as far south as Georgia and as far north as Quebec and Ontario.

Importantly, only 19 of the 559 understory herbaceous species we assessed (3.4%) were ephemeral in every cell they were observed in (i.e., EI = 1). Further, only six of these species (*Cardamine angustata, Krigia dandelion*, *Muscari botryoides*, *Narcissus pseudonarcissus*, *Scilla siberica*, and *Viola bicolor*) were observed in more than two cells, suggesting that “true ephemerality”, where a species is defined across its entire range, is rare in widely-dispersed species. However, there are some caveats to this conclusion. First, the scale at which access to light limits performance is at the individual level, meaning that our 100 km x 100 km estimates of canopy-understory overlap may be too coarse to accurately estimate species-level ephemerality characteristics. Moreover, this could be especially true if access to spring light affects annual survival probability (as has been shown for some understory tree species, Lee and Ibanez 2021a) – ephemeral wildflower species may be more likely to be found in microenvironments that have particularly delayed canopy closure phenology at a scale that is much finer than we capture in this analysis (Dion et al. 2017).

Another important caveat in the interpretation of our results is that both understory and overstory phenology is shifting in response to climate. For example, several recent studies provided evidence that canopy tree phenology is shifting earlier in the year at a faster rate than understory wildflower phenology is shifting (Heberling et al. 2019, Lee et al. 2022, Miller et al. 2022; but see Alecrim et al. 2023), a pattern which has continued for at least the past 120 years and which is projected to continue into the future. Because we estimated wildflower active period length from observations made only over a recent seven-year-long period (2015-2021), we find it likely that our results reflect higher overlap between wildflower and tree phenology than historically occurred, and therefore that our results are likely biased in favor of conservative ephemerality estimates. Furthermore, because access to spring light is expected to be reduced even more in the future (Lee et al. 2022), it is possible that some ephemeral species will no longer be defined as such under warmer future spring conditions. This seven-year-long window also relied on non-standardized surveying from volunteers on the platform iNaturalist, where the sampling effort is not uniform across species’ ranges leading to discrepancies among observations spatially which limits the representation across ranges (Di Cecco et al. 2021).

Despite these caveats, several wide-ranging species had high EI values very close to one (e.g., *Cardamine concatenata*, Fig. 3G), suggesting that a lack of wide-ranging true ephemerals may be partially precluded by exceptional individuals or populations that have anomalously late phenology that is not reflective of a species as a whole. On one hand, this highlights a potential limitation in our analytical approach, suggesting that our results are influenceable by the quality and scope of community-science observations. This can be problematic given that the data we used rely on non-expert plant identification collected with non-uniform sampling effort. On the other hand, however, we argue that it also represents a significant strength of our approach because it reinforces the notion that the spring ephemerality strategy is not a simple binary, but rather a gradient in plant resource acquisition that is messy because nature is messy. Species like *C. concatenata* are still defined as highly ephemeral from their high EI values, and it provides researchers and practitioners with a fuller context with which to differentiate among species phenological strategies. Furthermore, the potential bias applies to species with EI values very close to zero as well, such as *Alliaria petiolata* (Fig. 3C). This implies that potential miscategorizations evenly affect species from across the gradient of ephemerality, which support that our overall conclusions about where ephemerality occurs across the landscape are robust.

Still, our results indicate that the vast majority of spring ephemeral wildflowers only exhibit ephemeral behavior in a portion of their range, suggesting that inconsistencies in past species-level designations could be the result of range-level differences in expressed phenology. For example, *Trillium grandiflorum* is a charismatic spring-active wildflower with a wide range distributed across eastern North America and that is commonly referred to as a spring ephemeral in the scientific literature (e.g., Lubbers & Lechowichz 1989, Irwin 2001; but see Lapointe 2001 for an example of where it is cited as a spring-active wildflower). In contrast, we found *T. grandiflorum* to have an EI value of 0.35, suggesting that this species is likely not a true ephemeral, albeit with ephemeral designations occurring across the entirety of the species’ observed range (see Supplemental Material II). This incongruence could arise from the fact that flowering occurs and is generally completed by the initiation of canopy closure in this species. Individuals maintain their leaves and develop fruit until well into the summer shady period, which indicates a spring-active, but not ephemeral, wildflower species. We suspect that most who classify *T. grandiflorum* as a spring ephemeral are likely doing so on the basis of their flowering phenology alone, rather than their activity period as a whole. To resolve this potential confusion, we recommend that such discrepancies should be researched on a species-by-species basis in the future, with multi-year observations on the same individuals and populations when possible.

Range-level variation in ephemerality is further important because it suggests that species are likely to be variably vulnerable to climate change in different parts of their range. Previous work suggests that North American spring ephemerals are likely to lose access to spring light under warming climates as tree leaf out timing increases at a faster rate than wildflower emergence in the spring (Heberling et al. 2019, Miller et al. 2022, Lee et al. 2022; but see Alecrim et al. 2022). Thus, in areas where wildflowers have particularly large spring light windows (and therefore areas where they are more likely to be defined as ephemeral in this study), they may be less vulnerable to reductions in spring light compared to those in areas where their springtime activity already overlaps with closed-canopy conditions. However, it is difficult to derive strong conclusions from observational studies, such as this one, and field-based experiments should evaluate the strength of this speculation.

There was also substantial variation in the distribution of cells in which partially ephemeral species (i.e., those with 0 < EI <1) were found to be ephemeral. For example, species like *Dicentra cucullaria* (Fig. 3D) appeared to have a relatively random distribution of cells where it was and was not considered ephemeral. In contrast, other species like *Ranunculus abortivus* and *Oxalis violacea* (Fig. 3E&F) had distributions of ephemerality that appeared to correlate with spatial or environmental gradients. In the case of *R. abortivus*, cells where it was classified as ephemeral were clustered primarily in the south while cells where it was not classified as ephemeral were clustered primarily in the north. For all partially ephemeral species, further research will be needed to ascribe mechanisms to these patterns of spatial variability, likely through experimental transplantation and common garden experiments. Maps of the ephemerality distributions of all 103 species with EI > 0 are included in the second Supplement.

### Environmental drivers of spring ephemerality

In general, results from our models indicate that ephemeral species are most common in central latitudes, with peak ephemeral richness at around 40 °N (Fig. 5A). This pattern was primarily driven by variation in average April temperatures, which were tightly correlated with latitude (Supplemental methods, Fig. S6) and showed a similar quadratic relationship with a peak at middle temperatures (Fig. 6). This strong association with temperature likely reflects a tradeoff between frost and shade avoidance strategies, but further research is needed on the mechanisms explaining this biogeographic pattern we found. Spring ephemeral activity is also bounded by the risk of damage from frost events (Augspurger and Salk 2017). If these species emerge from dormancy too early, late-season frosts can damage foliar and floral tissue (Augspurger and Salk 2017, Gezon et al. 2016), thereby limiting carbon gain and reproductive success. In worst-case scenarios, frost damage can cause whole-plant mortality. This balance between frost avoidance and shade avoidance, a strategy often referred to as ‘phenological escape’ (Jacques et al. 2015; Heberling et al. 2019a, Lee and Ibáñez 2021a,b), directly determines a spring ephemeral’s capacity for annual carbon assimilation (Heberling et al. 2019a). This relationship then cascades to affect seed set success, growth, and survival of these species (Kudo et al. 2008), meaning that access to early seasonal light is a strong selective pressure that determines how and where these species can persist across the landscape.

Temperate forests at both the northern and southern extremes of eastern North America may potentially have a higher composition of evergreen tree canopy species, as the forest type transitions to northern hardwoods-pine/hemlock and oak-pine or subtropical evergreen, respectively (Dyer 2006), or overall extremes in canopy phenology. There is thus less room for opportunistic light exposure for understory wildflowers. Furthermore, in northern regions it is also likely that the window of favorable growth conditions before canopy closure is not beneficial for species utilizing the spring ephemeral strategy because of frost limitations. The duration between the date of last frost and canopy closure decreases with latitude, resulting in a duration of light availability too short to support the spring ephemeral life history strategy (Neufeld and Young 2014). In southern latitudes, the window for optimal growth likely extends much longer, where species can take advantage of windows of high light in both the early spring and in late fall, after canopy leaf fall, making the evergreen strategy more advantageous in southern deciduous forests (Neufeld and Young 2014). For these reasons, we speculate that forest overstory composition and phenology plays a strong limiting role in the distribution of ephemerality across the landscape.

Importantly, however, our analysis is agnostic to the forest type in which each of the 642,526 observations were made, meaning we are not able to investigate how forest canopy deciduousness affects species level ephemerality designations or the distribution of spring ephemerals across the landscape. It is possible that other drivers are responsible for the reduction in ephemeral wildflowers at northern and southern extremes. Still, previous research in eastern North America found that dominant forest canopy type influenced understory species richness, with highest richness observed in northern hardwood forests and relatively lower richness in evergreen and mixed-deciduous forest types (Ellum et al. 2010), echoing the results we found with respect to understory species richness in this study (Fig. 4D). Further research will be required to test these hypotheses explicitly, though.

Spring ephemeral richness and proportion were also significantly positively associated with May precipitation (Fig. 5). Precipitation is more commonly found in northern temperate forests to correlate with senescence phenology at the end of the growing season rather than with start-of-season phenology (e.g., Xie et al. 2015), so we infer that the correlations with precipitation found in our analysis indicate drivers of ephemeral wildflower senescence. Physiologically, one possible mechanism is water availability limiting photosynthetic rates. Spring ephemeral wildflowers often have elevated maximum photosynthetic rates as a strategy to make optimum use of high light availability in a short amount of time (Taylor and Pearcy 1976; Kudo et al. 2008; Heberling et al. 2019), a strategy that negatively affects their shade tolerance and likely contributes to the ephemerality of their activity in general (Neufeld and Young 2014). Insufficient water availability can limit photosynthesis through reduced stomatal conductance (Flexas et al. 2006), and researchers have previously demonstrated that reduced water availability negatively affects spring ephemeral wildflower leaf traits (including photosynthesis; Dion et al. 2017, Sawada et al. 1997, McKenna and Houle 2000). Further, others have found that reduced water availability scales up to affect population-level demography and community-level productivity of spring ephemeral wildflowers (Vasseur and Gagnon 1994, Axmanova et al. 2011). Climate change-driven shifts in water availability are also predicted to indirectly affect plant performance by altering species interactions and community dynamics (e.g., Hajek and Knapp 2022).

The last environmental driver that significantly correlated with spring ephemeral richness and proportion was elevation, which was negatively associated with richness and proportion of spring ephemeral wildflower species (Fig. 6). This result was unexpected because we anticipated that the broad scale that our data was collated at (100 km x 100 km grid cells), would obscure the finer-scale effects that elevation has on individual and population-level plant performance. We believe that the negative association with elevation is likely caused by increased frost risk and a reduction in suitable habitat at high elevations – habitats of many spring-active species’ are described to be in low elevation forest types (e.g. mixed mesophytic, floodplain, and cove forests; Weakley 2022). Temperature is generally negatively correlated with elevation, and previous studies have demonstrated strong phenological responsiveness to elevational gradients in spring temperature (Vitasse et al. 2018, Čufar et al. 2012). Other drivers such as nutrient availability (Mayor et al. 2017), growing season length (Berdanier and Klein 2011), and soil fungal community (Liu et al. 2023) are also strongly related to elevational gradients, with the general pattern of reduced habitat quality at higher elevations. Our study encapsulated a broad elevational gradient ranging from coastal lowlands to Appalachian mountaintops, so it is not surprising that our model was sensitive to this wide amount of environmental heterogeneity.

### Conclusions

Spring ephemeral wildflowers are ubiquitous in eastern North American temperate forests, but to date this charismatic group has lacked strict definition and biogeographic description. Here, we show that spring ephemeral wildflowers are a diverse group of species that comprise a considerable amount of total understory biodiversity. Further, we found that species’ phenological strategies are not fixed across their range, with many species that are often colloquially referred to as ephemeral lacking that trait in parts of their range. Lastly, we found strong evidence that continental-scale environmental gradients are correlated with the distribution of spring ephemeral wildflower species across the landscape. Our novel method to quantify phenological strategy can be applied to other regional floras of the world to test whether these patterns are shared across other temperate forests of the world. Future research is needed to understand the mechanisms which explain the biogeographic pattern we report. Taken together, these lines of evidence suggest to us that wildflowers exhibiting spring ephemerality are particularly vulnerable to asynchronization resulting from ongoing climate change (Heberling et al. 2019). These species have been noted with limited abilities to adjust physiologically to varying light conditions (Taylor and Pearcy 1976). As the early spring light window changes across North America (Lee et al. 2022), we expect spring ephemerals to suffer disproportionately compared to other understory wildflowers.

## Supporting information

Supplemental Data Descriptions

Supplemental Materials - Ephemerality Maps

## Acknowledgements

The authors of this paper thank Rob Guralnick and Daijiang Li for their advice in our analytical approach and for providing feedback on early drafts of this article. We also acknowledge the numerous botanists, ecologists, and volunteers that worked to construct the species lists used in this study and, especially, the 71 organizations and over 57,000 observers that contributed observational data to the GBIF collection. This project was supported through the US National Science Foundation (NSF DEB 1936971 to JMH (including REPS supplement funding AJY), NSF DEB 2223675 to SK, and NSF DBI 2108128 to BRL.

