## Supplemental Figures, Tables, and Methods for "Evaluating the definition and distribution of spring ephemeral wildflowers in eastern North America"

#### Supplemental Tables and Figures:

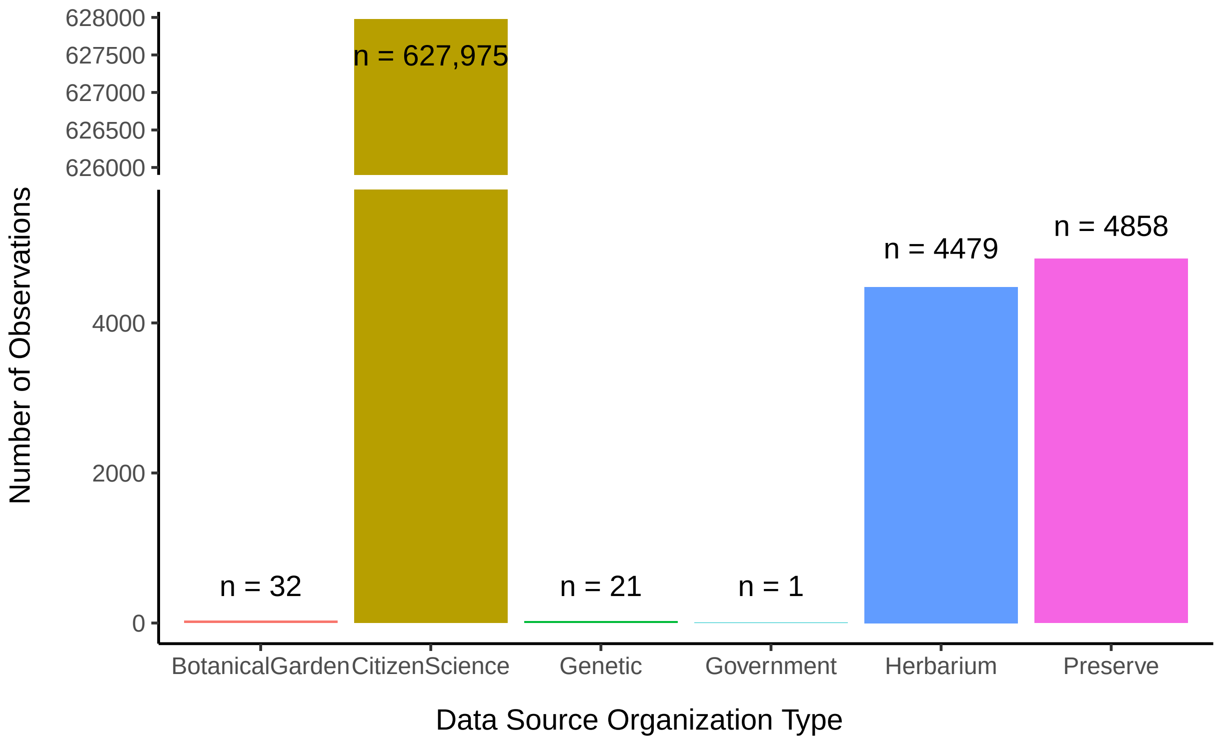


**Fig. S1**: Number of observations from different types of data sources used in this study. A full list of the 72 different data sources is provided in the Supplemental Data.

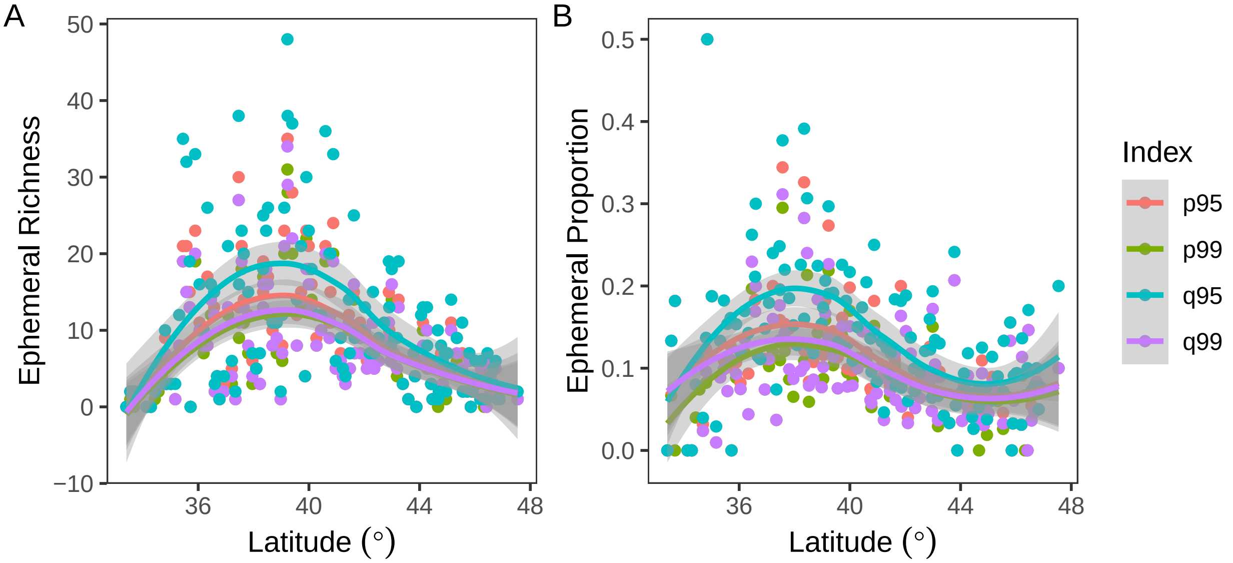


**Fig. S2**: Modeled fits of how (A) ephemeral species richness and (B) proportion of ephemeral species relative to total understory herbaceous species richness relate to latitude. Points represent cell-level values of the responding variables with different colors indicating different estimation indices. Indices beginning with “p” are based on ephemerality definitions using the `phenesse` package whereas those beginning with “q” are based on quantile estimates of raw data. The end of the index names reflect either the 95th or 99th percentile cutoff point. Relationships were fit using the default Loess fit in the `stat_smooth` command of the `ggplot2` package and gray shading represents 95% confidence intervals.

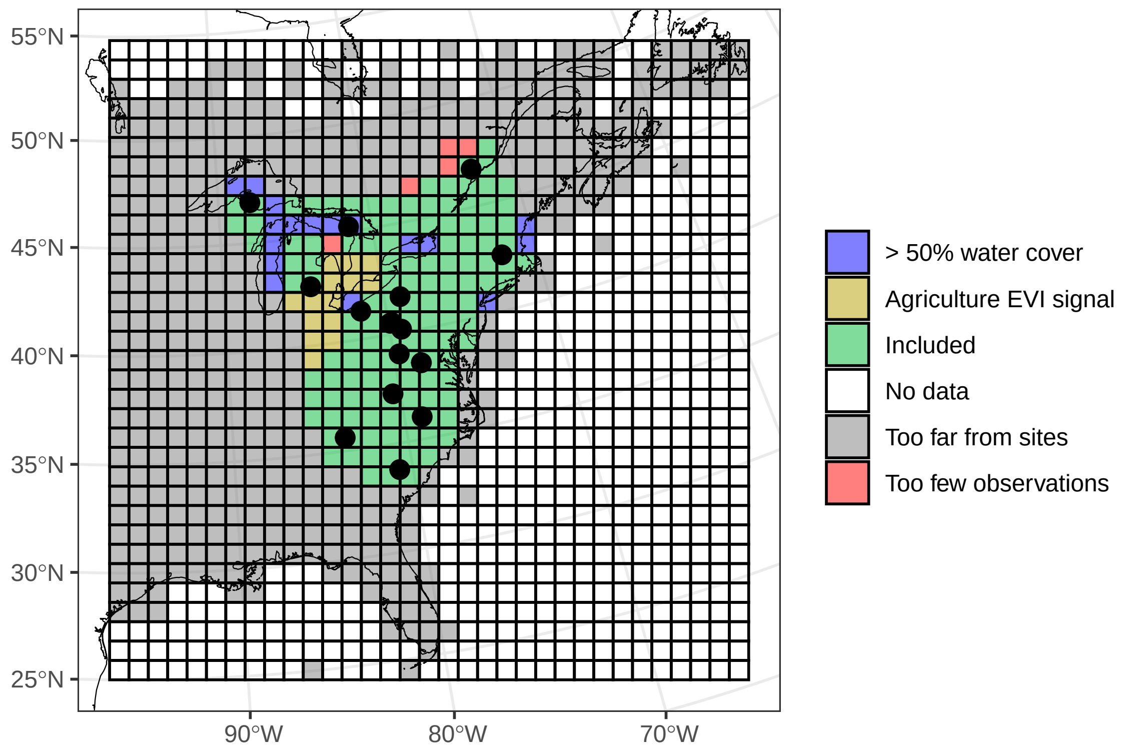


**Fig. S3**: Map of the original 1,089 100 km x 100 km grid cells considered in data set generation. Colors indicate whether cells were included (green, n = 128) or excluded due to one of four criteria (in order of filtering steps): lack of iNaturalist data for species in the combined species list (white, n = 482), being too far away from sites where species lists were assembled (grey, n = 443), too few observations of species in filtered species list (red, n = 5), grid consisted of over 50% water cover by area (blue, n = 17), or estimated day of canopy close was biased by summer-green crop cover (gold, n = 14). Black points show the locations of the 16 sites used to create the combined species list.

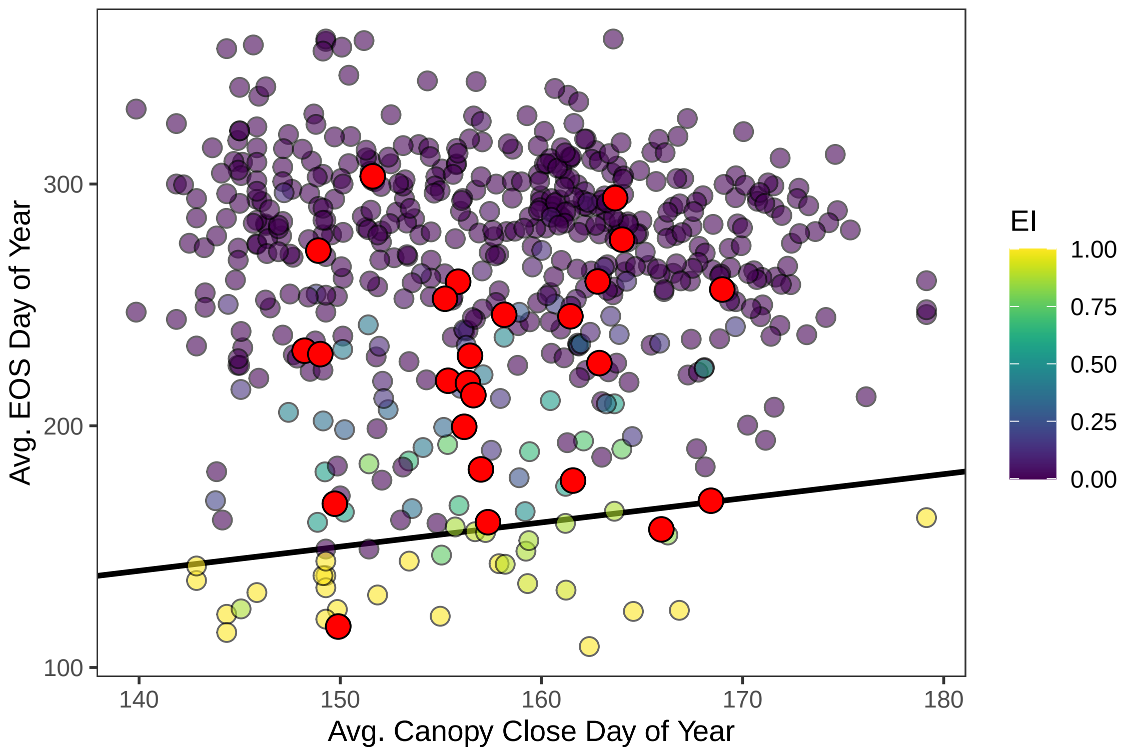


**Fig. S4**: Species level relationships between average canopy close date (day of year) and average species end of season (EOS, day of year). Averages were calculated only using cells where a species was present. Solid black line is the 1:1 line, so points below the line tend to senesce prior to canopy close and those above it tend to maintain activity into the growing season. Point colors indicate species-level ephemerality index (EI) values ranging from never ephemeral (EI = 0, purple points) to always ephemeral (EI = 1, yellow points). The 25 evergreen species for which EI values were manually changed to zero (despite being estimated as being ephemeral in at least part of their range) are indicated with red points.

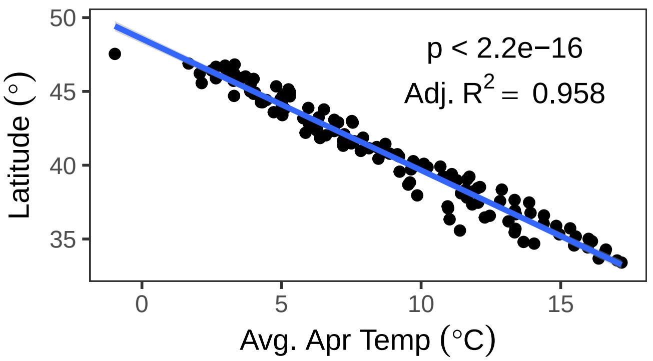


**Fig. S5** : Relationship between average April temperature and latitude at the grid cell level.

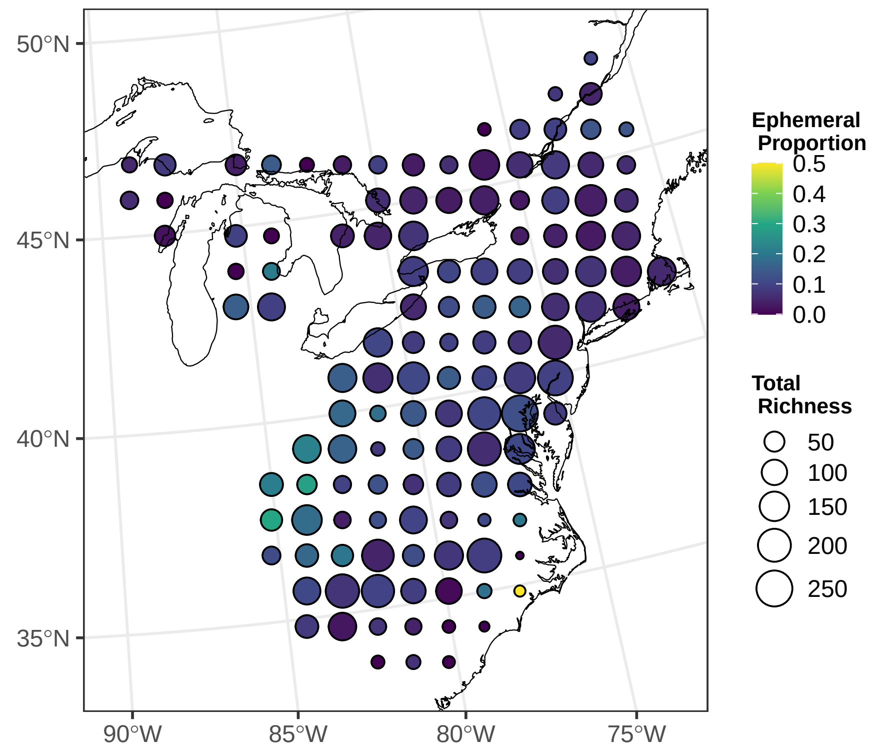


**Fig. S6**: Full version of Figure 4F in the main text, which omits the cell centered at 34.84695 deg. N, -77.71915 deg. W. This cell had an abnormally large proportion of ephemeral species (0.5), but largely only because there were only four species that were assessed in that location.

**Table S1**: List, source, and location of site species lists used to amass the initial understory plant species list used in this study.

| Site Name | Source | Coordinates (°N, °W) |
|---|---|---|
| Huron Mountain Club (HMC) | Spicer et al. (2020) | 46.89, -87.87 |
| La Mauricie (LM) | Spicer et al. (2020) | 46.83, -73.00 |
| Bruce Peninsula (BP) | Spicer et al. (2020) | 45.22, -81.50 |
| Allegheny National Forest (ANF) | Spicer et al. (2020) | 41.49, -70.10 |
| Powdermill Nature Reserve (PNR) | Spicer et al. (2020) | 40.16, -79.27 |
| Fernow Experimental Forest (FEF) | Spicer et al. (2020) | 39.05, -79.67 |
| Southern Appalachia (SA) | Spicer et al. (2020) | 37.29, -80.43 |
| Duke Forest (DF) | Spicer et al. (2020) | 35.87, -80.00 |
| Harvard Forest (HF) | Jenkins and Motzkin (2009) | 42.53, -72.19 |
| Trillium Trail (TT) | Hale et al. (2011) | 42.53, -72.19 |
| Congaree National Park (CONG) | https://irma.nps.gov/NPSpecies/ | 33.79, -80.77 |
| Great Smoky Mountains National Park (GRSM) | https://irma.nps.gov/NPSpecies/ | 35.69, -83.54 |
| Cuyahoga Valley National Park (CUVA) | https://irma.nps.gov/NPSpecies/ | 41.28, -81.57 |
| Shenandoah National Park (SHEN) | https://irma.nps.gov/NPSpecies/ | 38.47, -78.45 |
| Baker Woodlot (BW) | Kolp et al. (2020) | 42.71, -84.47 |
| Barking Slopes (BS) | JM Heberling, unpublished data | 40.53, -79.79 |

See *Supplementary References Cited* for full citations of listed publications

**Table S2**: List of evergreen species that were misclassified as spring ephemeral in at least one 100 km x 100 km grid cell. These species were manually assigned EV values of zero in the statistical analysis.

| Species | Initial EVI value |
|---|---|
| *Allium canadense* | 0.143 |
| *Allium tricoccum* | 0.017 |
| *Aquilegia canadensis* | 0.049 |
| *Chrysogonum virginianum* | 0.091 |
| *Erigeron pulchellus* | 0.231 |
| *Geranium maculatum* | 0.213 |
| *Geum vernum* | 0.333 |
| *Hesperis matronalis* | 0.015 |
| *Houstonia caerulea* | 0.246 |
| *Houstonia pusilla* | 0.957 |
| *Mitella diphylla* | 0.244 |
| *Packera aurea* | 0.146 |
| *Packera glabella* | 0.111 |
| *Panax trifolius* | 0.489 |
| *Phlox divaricata* | 0.698 |
| *Phlox stolonifera* | 0.5 |
| *Salvia lyrata* | 0.024 |
| *Silene caroliniana* | 0.7 |
| *Silene virginica* | 0.033 |
| *Tiarella cordifolia* | 0.5 |
| *Vinca major* | 0.05 |
| *Viola labradorica* | 0.625 |
| *Viola pedata* | 0.231 |
| *Viola primulifolia* | 0.333 |
| *Waldsteinia fragaroides* | 0.091 |

**Table S3**: List of theoretical and statistical assumptions included in this analysis along with justifications for incorporation of those assumptions.

| Type | Assumption | Justification/Citations |
|---|---|---|
| Theoretical | Definition of spring ephemeral species as having completed the entirety of their growing season before the canopy closes in early summer | Our definition of ephemerality is based on a phenological definition of growing season length (sensu Körner et al. 2023) and informed by previous papers that have categorized plants into phenological strategies and "syndromes" (Uemura 1994, Neufeld and Young 2014) |
| Theoretical | 569 species list is representative of understory wildflowers in eastern North American deciduous forests | The combined species list originally comprised 1666 species, out of which we considered 834 (50.0%) that were categorized as herbaceous angiosperms (i.e., excluding grasses, ferns and understory woody plants). Spicer et al. (2020) estimated that ~80% of plant diversity in temperate deciduous forests is from forest floor species, it makes sense that our value is less than 80% since we excluded some groups that are present in their estimate. Although we were unable to include all herbaceous understory plants present in the combined species list (primarily due to lack of observational data for many species), and although the combined species list is likely to have missed some species (especially those that are rare), we find it reasonable to assume that our systematic sampling approach makes the final selection of 569 species relatively representative of species present in this forest type. |
| Statistical | End of season phenology is accurately estimated from the bootstrapped, repeated sampling approach described in the *phenesse* package | This estimation approach has been shown to perform the best when compared to similar statistical methods (Belitz et al. 2020) and avoids making the assumption that the temporal range of observations is the "true" limit to a species' activity period. |
| Statistical | iNaturalist data is inherently biased by observer effort, which can then be accounted for using boot-strapping approaches and by accounting for human population density in modeling design | Related to the point above, the Belitz et al. (2020) estimation approach is specifically designed to account for differences in sampling effort (approximated in this study with number of observers/observations). We also account for human population density in our models which has previously been shown to affect estimation of phenological trends due to associations with sampling effort (Li et al. 2019). |
| Statistical | Canopy close phenology (and thus start of the shady period) is accurately estimated from MODIS EVI data | Peng et al. (2017) previously found that this metric is highly correlated with direct measurements of understory light availability, supporting our use of it in this study. |

| Type | Assumption | Justification/Citations |
|---|---|---|
| Statistical | Cells with EVI maturity dates of DOY >= 181 indicate bias from agricultural green up of summer-green crop species and should be excluded from analysis | The MODIS EVI variable quantifies green up but does not discriminate by land cover type. Summer-green crops often signal late in summer/early in fall (Wardlow et al. 2007), whereas deciduous forests peak in late spring. We thus ruled out any cells where canopy maturity was estimated to occur in the second half of the year. |

### Supplemental Methods:

#### Justification of drivers included in final model design

As described in the main text, we evaluated the role that several environmental drivers play in shaping the richness (and proportion relative to total number of all understory herbaceous species) of spring ephemeral wildflowers across eastern North America. Our final model design included the following drivers: Elevation, May precipitation, human population density, and squared April temperature. Additionally, the richness model included a driver of total understory species richness. This driver, along with human population density, were meant to account for potential bias where higher observer effort could lead to inflated estimates of how many spring ephemeral wildflower species occur in a given location.

However, these were not the only drivers which we considered. Here, we describe results from our preliminary analysis of possible drivers and justify our selection of drivers used in the final model structure.

#### Data

The `cells` data frame is freely available for download (see data supplement). The data includes many columns, but the columns containing the data and drivers of interest are as follows:

**Table S4**: List and description of variables explored in preliminary data analysis.

| Abbreviation | Definition | Unit |
|---|---|---|
| cell.no | Dummy variable for cell ID | |
| NSp | Number of total species observed in each cell | Species |
| ydeg | latitude of cell center | Degrees |
| NEphem.p99 | Richness of spring ephemeral species using 99% cutoff in phenesse | # Species |
| PropEphem.p99 | NEphem.p99/NSp | |
| Tavg.## | Average monthly temp. from ## month or range of months (1970-2000) | deg. C |
| Popdens | Human population density (as of 2020) | People per sq km |
| Tmin.## | Avg. minimum monthly temp. from ## month or range of months (1970-2000) | deg. C |
| Prec.## | Mean precipitation in ## month (1970-2000) | mm |
| Elev | Mean elevation | meters above sea level |

Next, we describe the preliminary analysis driver by driver:

#### Latitude

First, we looked for geographic variation in ephemeral richness in the form of latitudinal gradients. There are many biogeographical distributional patterns that have previously been related to latitude, not the least of which is that overall species diversity tends to be higher near the tropics compared to near the poles.

We looked at correlations between latitude of the center of each cell and our responding variables. In this preliminary analysis we used the default loess smoothing function in `ggplot2` using `stat_smooth`, just to get an idea of the shape of relationship. Each point in the *Fig. S7* represents one grid cell.

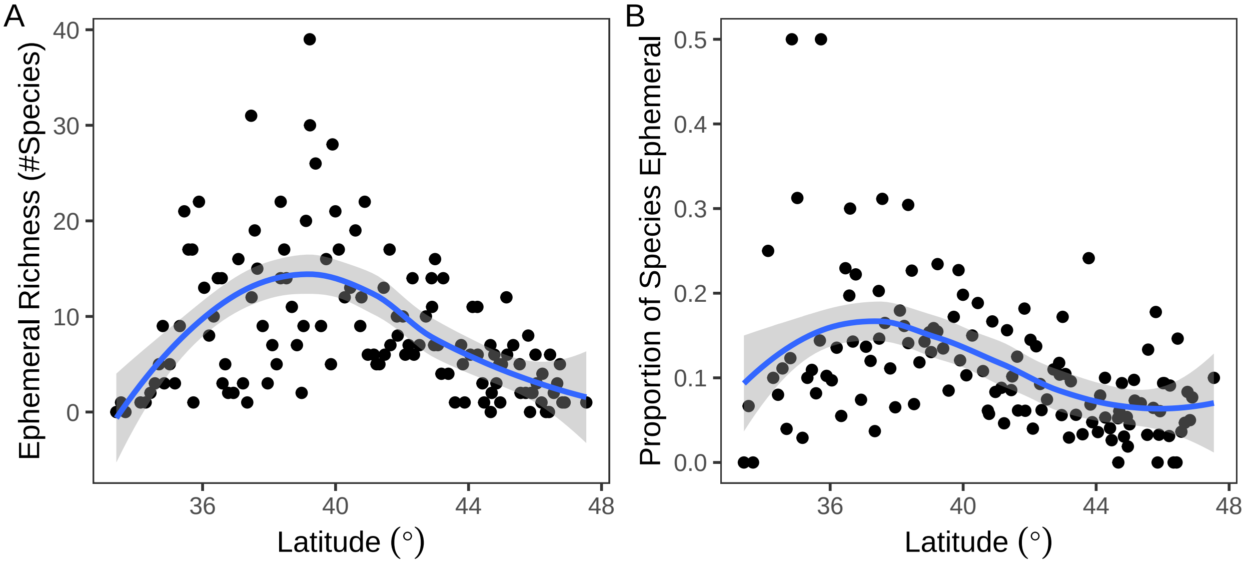


**Fig. S7** : Relationships between latitude and (A) ephemeral species richness, and (B) proportion of total understory herbaceous species that are considered ephemeral in each grid cell.

These graphs suggest quadratic relationships between latitude and our responding variables. Indeed, we present fitted quadratic relationships in **Fig. 4 A, C, & E** in the main text. However, as discussed more in the following section, we did not include latitude in the final model structure because it was highly redundant to relationships with average spring temperature.

#### Average spring and winter temperatures

The relationships with latitude are capturing a lot of variation in the data, but we also wanted to explore relationships with more mechanistic drivers. The first environmental driver we tested was temperature. Spring phenology tends to be strongly correlated with average spring temperature in eastern deciduous forests. Spring ephemeral wildflower performance has been shown to be strongly related to phenological sensitivity (Heberling et al. 2019), so it follows that their distribution would also be sensitive to this variable. Other studies have showed that the phenology of some plants (especially woody species) are also sensitive to winter chilling (i.e., "vernalization") effects (Lee and Ibáñez 2021, Ettinger et al. 2020).

For this analysis, we used 30 year average temperature values downloaded from WorldClim 2.1 (Fick & Hijmans 2017) for monthly intervals. The rationale for using this baseline data as opposed to averaged temp data from the period we downloaded the iNaturalist observations (2015-2021) is that it should be more representative of the period where the initial species lists were constructed (see *Table S1* and *Fig 1A*) and should also be more representative of spatial differences between grid cells because it captures a wider amount of interannual variation.

We initially evaluated correlations between our responding variables and nine different metrics of spring/winter temperature (*Fig. S8*): average monthly temperatures in December through June and multi-month averages in winter (December-February) and spring (March-April):

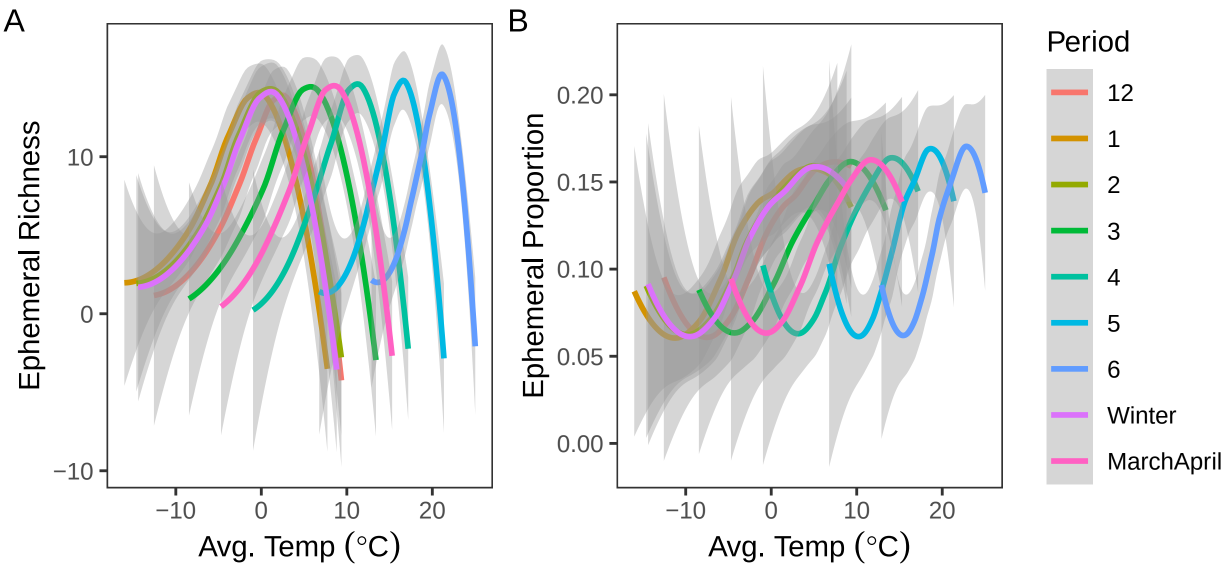


**Fig. S8** : Relationships between average monthly or multi-month temperature and (A) ephemeral species richness, and (B) proportion of total understory herbaceous species that are considered ephemeral in each grid cell.

Similarly to the latitude graphs, all relationships appear to be somewhat quadratic, regardless of which month or multi-month bin used. Interestingly, the relationships in winter months appear to be roughly the same shape as in spring months. This turned out to be primarily due to strong correlations between spring and winter temperatures (*Fig. S9*; i.e., cells with warmer average springs also had warmer average winters).

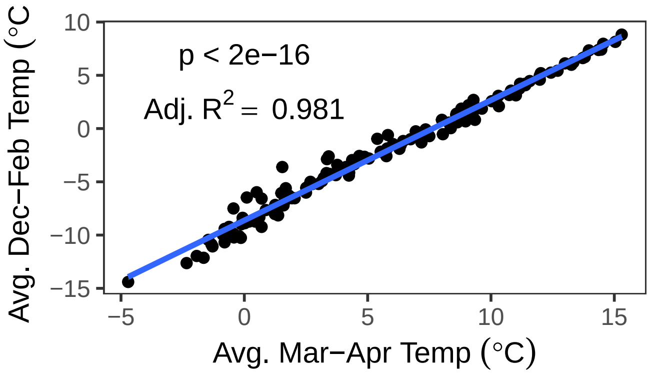


**Fig. S9** : Relationship between average spring and winter temperatures in the cell-level data set.

Still, we decided to specifically evaluate the correlation strength between the different temperature bins and our response variables using quadratic relationships (*Fig. S10*):

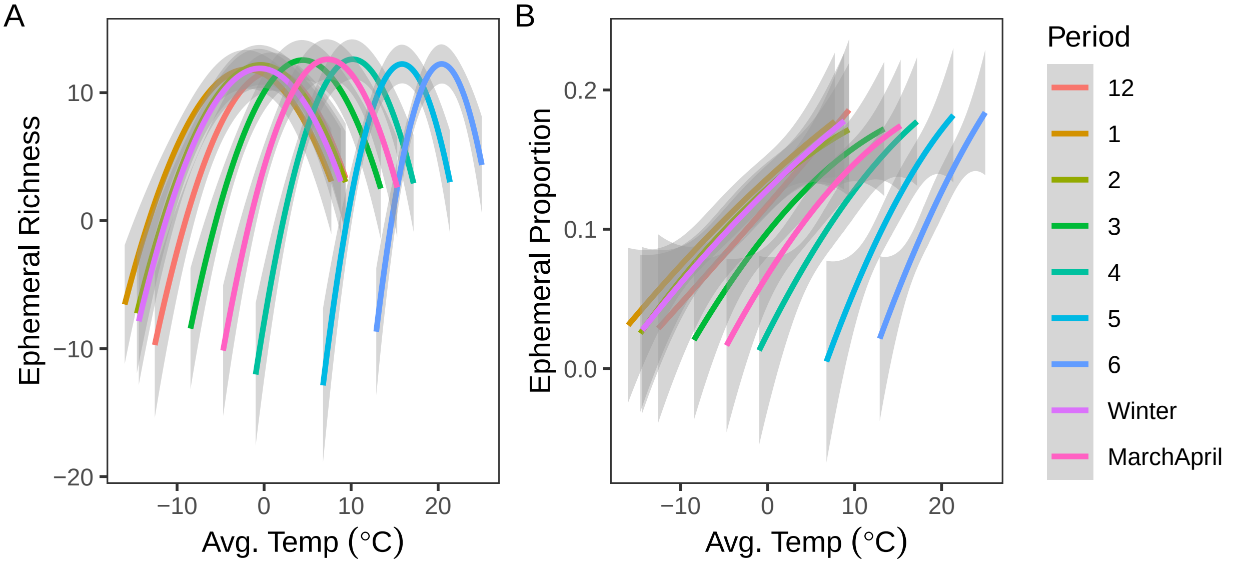


**Fig. S10** : Relationship between average spring and winter temperatures in the cell-level data set.

| Period | Richness | Prop.Ephem |
|---|---|---|
| December | 0.2396 | 0.1669 |
| January | 0.2466 | 0.1636 |
| February | 0.2607 | 0.1681 |
| March | 0.2866 | 0.1818 |
| April | 0.2949 | 0.189 |
| May | 0.2789 | 0.1902 |
| June | 0.2716 | 0.1966 |
| Dec.Feb | 0.2507 | 0.1661 |
| Mar.Apr | 0.2931 | 0.1853 |

**Table S5**: Adjusted $R^2$ values for the relationships between our responding variables and each period of average temperature explored.

Here, we see that average April temperature provides the best correlative strength with both of our response variables. Spring and winter temperatures were shown above to be redundant, so we decided to use only April temperature in this analysis. Further, since spring temperature often correlates strongly with latitude, we evaluated the strength of that relationship in our data set (*Fig. S5* above). As expected, latitude and spring temperatures are strongly (negatively) correlated. This confirmed our decision to not include latitude in the final model structure as a means of avoiding autocorrelational bias among drivers.

#### Minimum monthly/multi-monthly temperatures

We also extracted 30 year averages for minimum monthly temperatures. In some phenology models, average temperatures are used for spring forcing effects, but minimum monthly temperatures are more representative for winter chilling effects. We therefore decided to see if minimum winter temperatures were redundant or not to the average temperatures explored above.

We checked to see if monthly minimum temperatures were correlated with monthly average temperatures (*Fig. S11*). If they are, then there's no real point in going further since it will just be redundant to what we already have with the monthly averages:

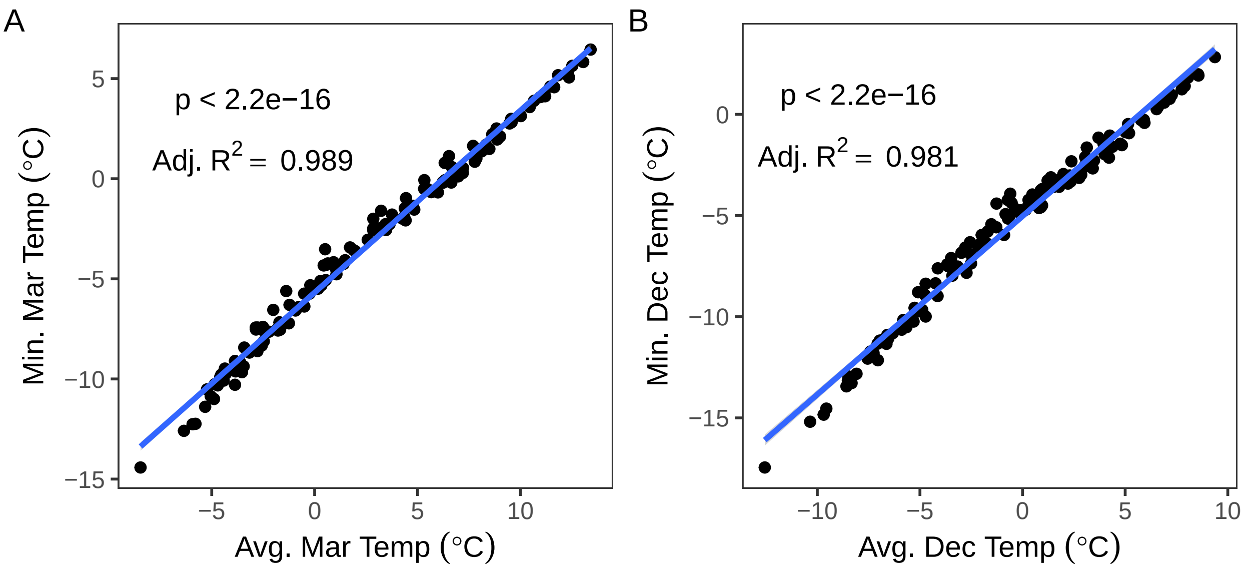


**Fig. S11**: Relationships between average and minimum monthly grid cell-level temperatures in (A) March and (B) December.

The results shown in *Fig. S11* suggest that there is high correlation between average and minimum temperatures in winter months. This means that minimum temperatures will be largely redundant to the average temperatures explored previously, and so we decided not to include minimum temperatures in the final model structure.

#### Mean monthly precipitation

Precipitation is something that isn't often tied to phenology in eastern deciduous forests, but it can sometimes be important in other systems, so we felt it was worth evaluating. If nothing else, precipitation can sometimes be decoupled from average monthly temperatures, meaning that it should at least have the potential to independently explain variation in the responses.

Relationships between precipitation and ephemeral richness and proportion seemed approximately linear. We evaluated relationship strength between our responding variables and seven monthly precipitation bins (*Fig. S12*, *Table S6*); December through June, months which we thought could be the most important with respect to affecting spring wildflower phenology):

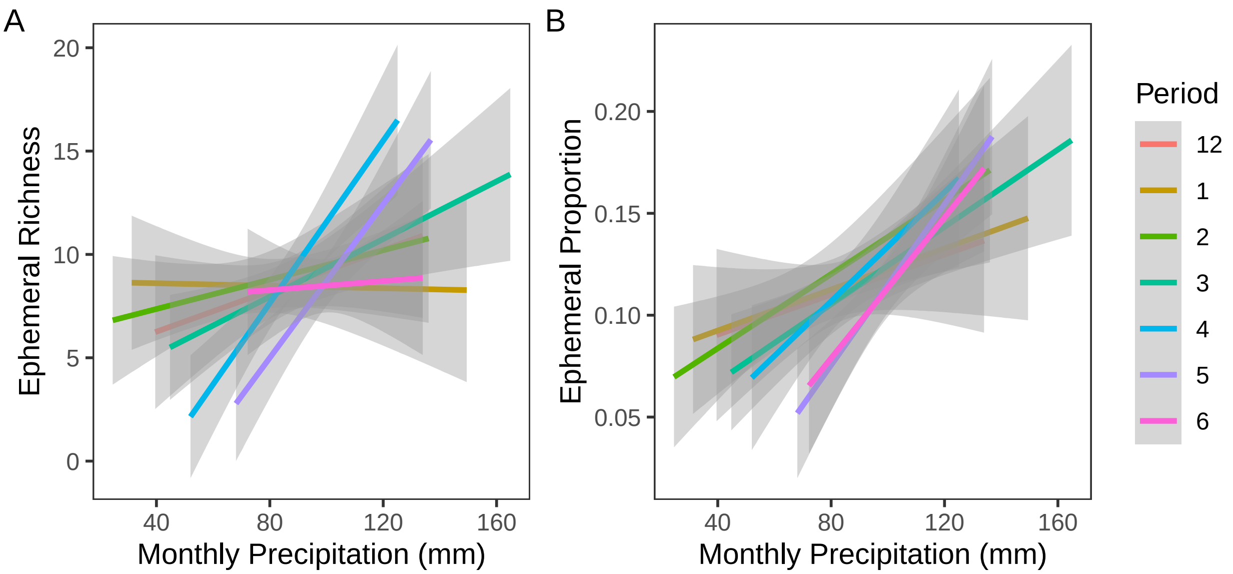


**Fig. S12**: Relationships between monthly precipitation and (A) spring ephmeral wildflower richness and (B) proportion relative to all understory herbaceous plant species.

| Period | Richness | Prop.Ephem |
|---|---|---|
| December | 0.004865 | 0.001884 |
| January | -0.00786 | 0.008535 |
| February | 0.002881 | 0.04721 |
| March | 0.04628 | 0.07001 |
| April | 0.1374 | 0.04457 |
| May | 0.1317 | 0.1147 |
| June | -0.007577 | 0.06262 |

**Table S6**: Adjusted $R^2$ values for the relationships between our responding variables and each period of monthly precipitation.

Here, most monthly periods showed very weak correlative strength with both of our responding variables. However, May precipitation (and to a lesser extent, April precipitation), showed relatively higher explanatory power. We therefore decided to add May precipitation to the final model structure.

#### Elevation

Elevation correlates with some of the other environmental variables (e.g., temperature decreases with elevation), but it also affects plants on a much narrower margin than what we are using in this type of model. Elevation can vary a lot over 10,000 square kilometers (the area of each grid cell in this study), so using a single, mean value for each cell is necessarily going to miss a lot of the topographic variation that could be driving the patterns we're interested in. Still, this variable could be useful in picking out differences associated with larger gradients (e.g., top of the Appalachians vs. Carolina lowlands).

First, we checked to see if elevation correlates strongly with either of the other two metrics so far added to the final model structure (*Fig. S13*):

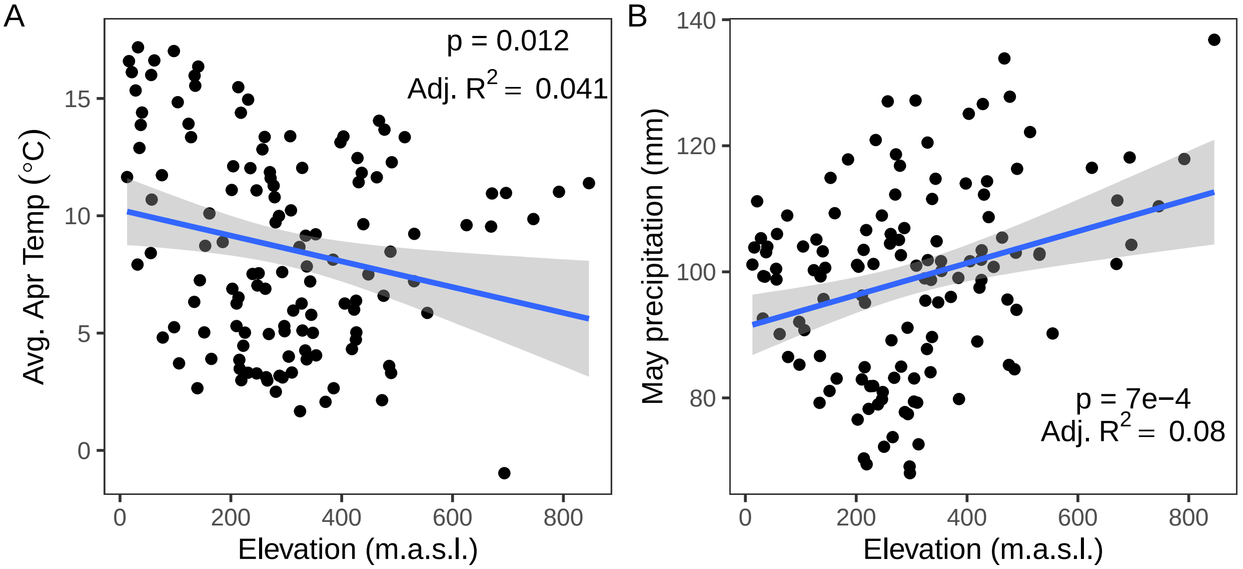


**Fig. S13**: Relationships between elevation and (A) average April temperature and (B) May precipitation.

While both variables were slightly correlated with elevation, neither relationship was very strong, suggesting that elevation could reasonably be included in the model without interfering too much with the other drivers.

Not unexpectedly, checking the (linear) correlation strength between elevation and our responding variables (*Fig. S14*):

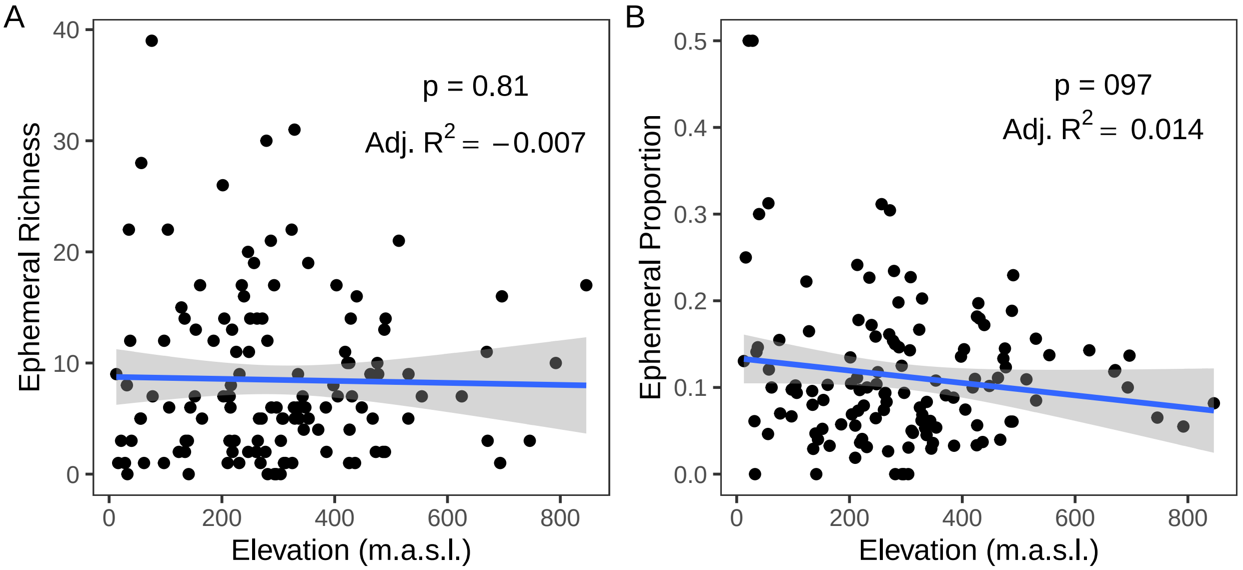


**Fig. S14**: Relationships between elevation and (A) Ephemeral Richness and (B) Proportion of species in each grid cell classified as ephemeral.

Elevation doesn't appear to have an important effect on either metric. Both relationships appear to be very flat, which fits our expectation of this study being conducted on a scale that is not well-suited for using elevation as an explanatory variable. Still, it is possible that, because it is independent-ish of our two other drivers, it could still have an important effect that only shows up in the residuals, so we decided to include it in the model.

#### Human population density

We explored correlations between our responding variables and human population density not because we expect there to be a direct causal link between them, but instead as a measure of sampling effort bias. Analyses involving iNaturalist and other community scientist-driven data sources can be biased by differences in observer effort. For example, observers are more active in summer months than they are in winter or early spring (Di Cecco et al. 2021). Observations also tend to be clustered around cities and national parks, with higher sampling effort around areas with large human populations (Mesaglio and Callaghan 2021).

We accounted for some observer effort already in our use of the `phenesse` package to bootstrap-estimate herbaceous plant activity periods (see Methods), but we also decided to evaluate the effects of human population density to see if it seemed important to include as a type of control driver in our analysis (*Fig. S15*, *sensu* Li et al. 2019):

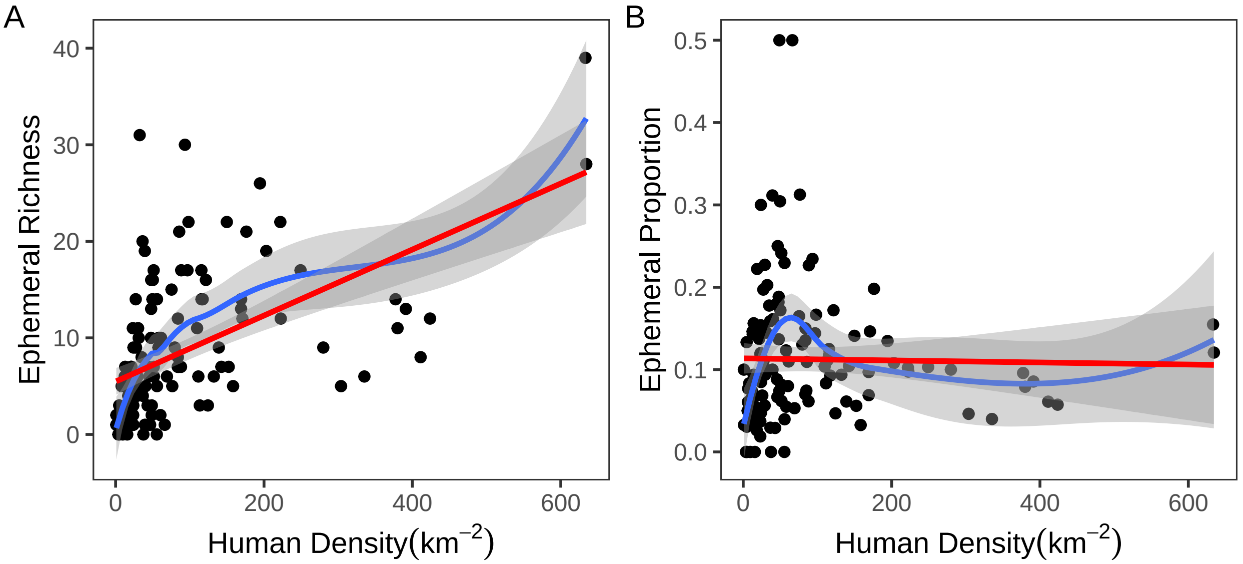


**Fig. S15**: Relationships between human population density and (A) Ephemeral Richness and (B) Proportion of species in each grid cell classified as ephemeral. Blue lines show fits using a Loess function whereas red lines indicate fits from a linear model.

We found the expected positive relationship between human population density and number of ephemeral species in *Fig. S15A*. The much flatter trend between population density and ephemeral proportion is also as expected – increased observer effort should lead to higher overall species richness, not just ephemeral species richness. As such, we decided to include human population density as an explanatory driver in both final model versions.
