## Supplemental Data Descriptions for "Evaluating the definition and distribution of spring ephemeral wildflowers in eastern North America"

Here we provide the descriptions of the supplemental datasets provided at **([TO BE ADDED UPON PUBLICATION]**):

| Data file: | Description: |
|---|---|
| Final GBIF_INAT Observation List.csv | This file contains all 642,526 observations used in this analysis, species identifiers, GBIF and iNaturalist identifiers, coordinates, and grid cell assignments. |
| GBIF data_filtered.csv | A subset of the final observation list, but only including observations downloaded from GBIF. All identifiers correspond to GBIF identifiers |
| iNat missing species data_filtered.csv | A subset of the final observation list, but only including observations downloaded from iNaturalist. All identifiers correspond to iNaturalist identifiers |
| Observation Sources.csv | A list of the 72 different data sources used in this analysis and the number of observations that came from each source |
| cells7_temp.prec.tmin.elev.hpop.cent.csv | Data aggregated at the cell level, including driver information used in the hierarchical Bayesian models. |
| Species level ephemerality indices.csv | Species-level Ephemerality Index values. Values reported in the main manuscript are in the final column: “EG.EI.p99”. |
| Sp x cell Ephemerality values.csv | Species-level ephemerality designations (1 or 0) for each cell in which it was observed. Values reported in the main manuscript are in column “EG.EI.p99”. |
