## Supplemental Materials - Ephemerality Maps for "Evaluating the definition and distribution of spring ephemeral wildflowers in eastern North America"

This supplement contains maps of the grid-cell level ephemerality designations for each of the 103 herbaceous understory wildflower species that were defined as ephemeral in at least 1 grid cell.

#### *Alliaria petiolata*

Ephemerality index: 0.044

n = 91

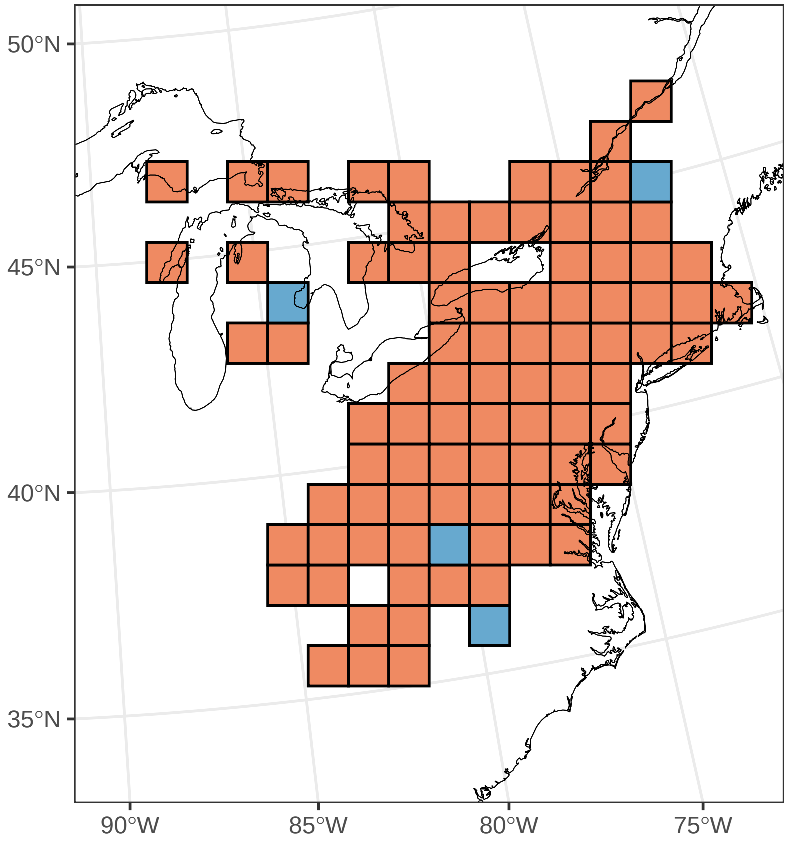

#### *Amsonia tabernaemontana*

Ephemerality index: 0.375

n = 8

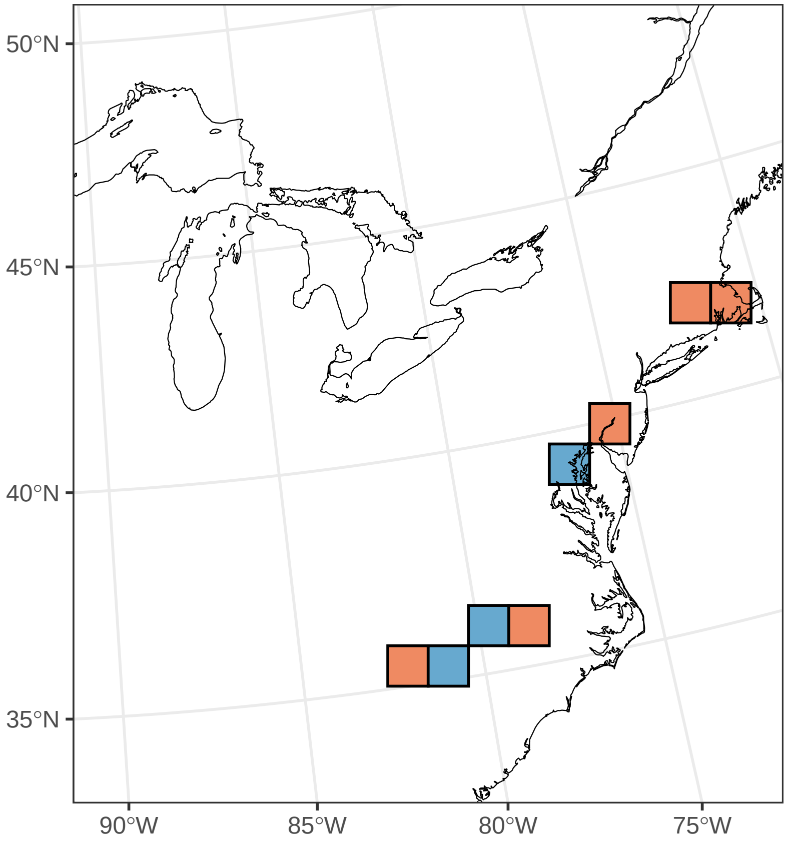

#### *Anemone lancifolia*

Ephemerality index: 1

n = 1

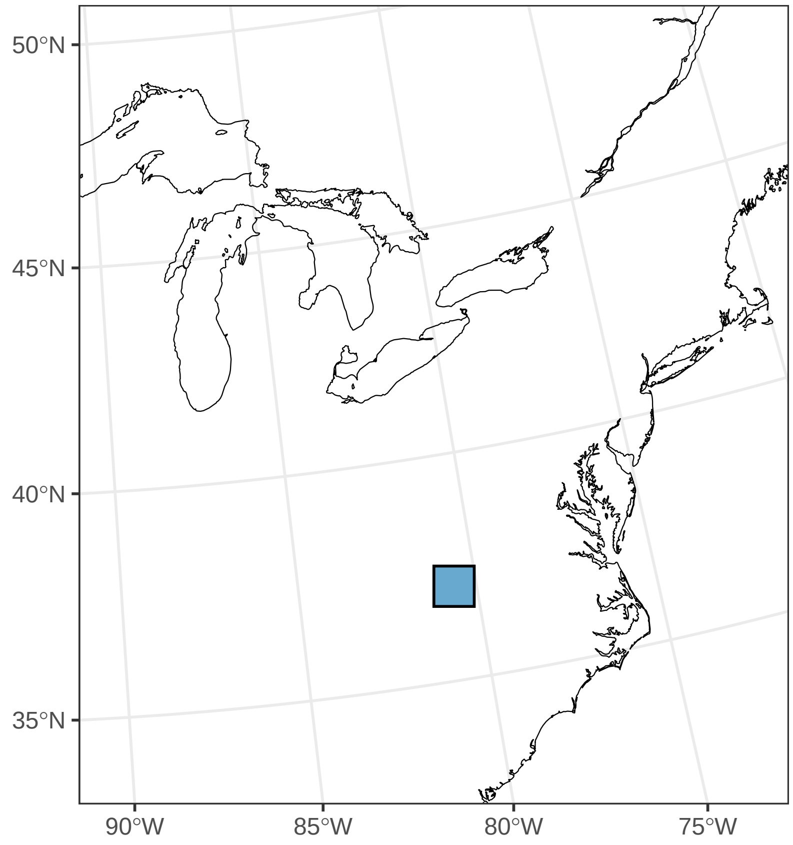

#### *Anemone quinquefolia*

Ephemerality index: 0.566

n = 53

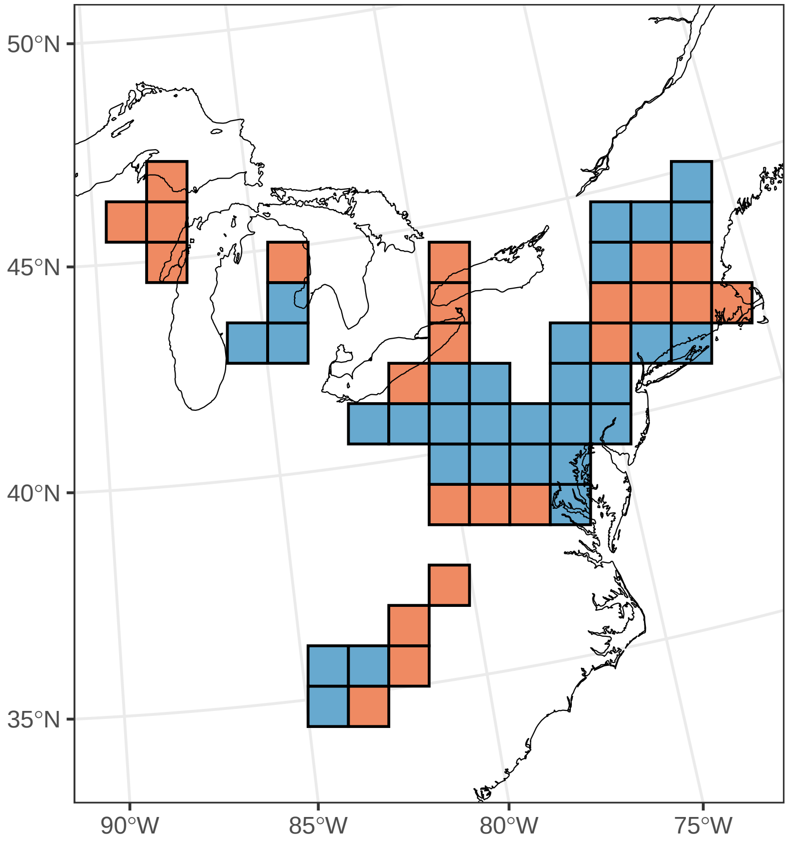

#### ***Antennaria solitaria***

Ephemerality index: 1

n = 2

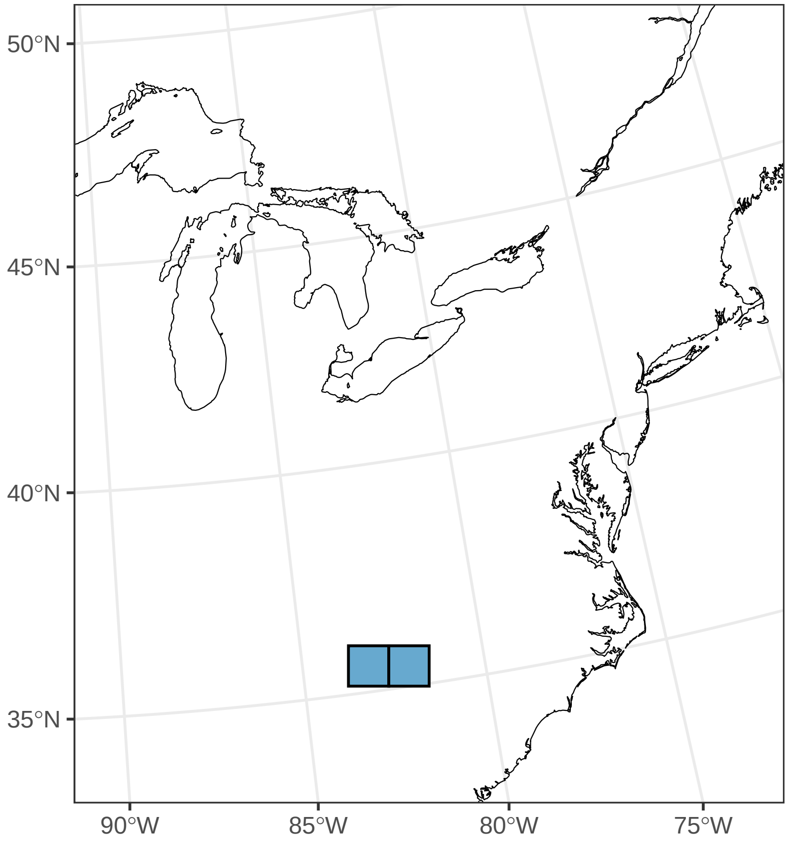

#### *Asclepias quadrifolia*

Ephemerality index: 0.067

n = 15

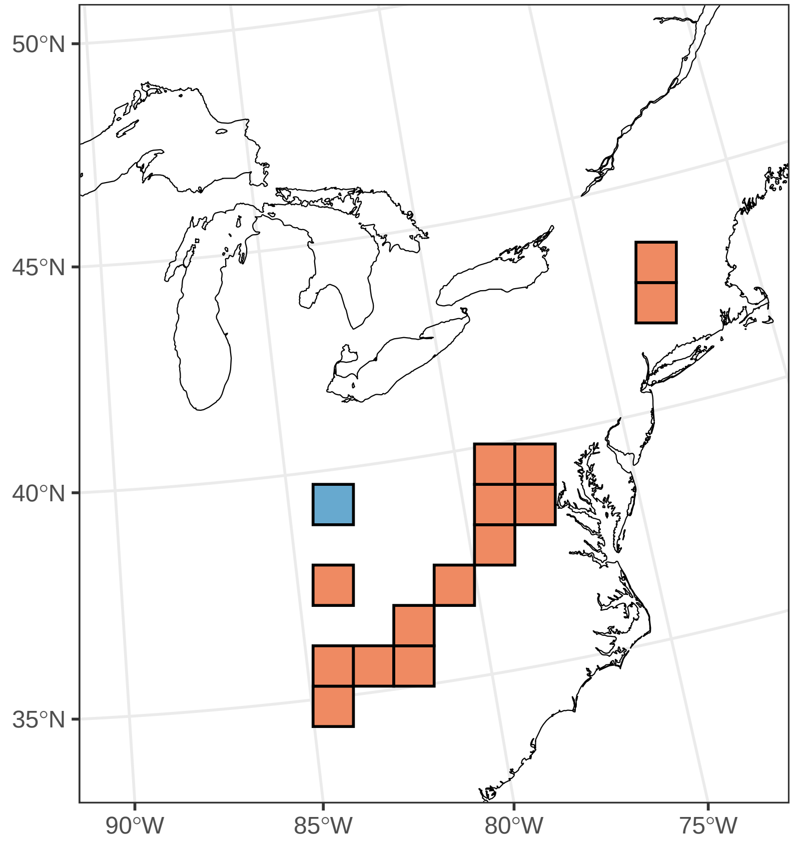

#### ***Camassia scilloides***

Ephemerality index: 0.5

n = 6

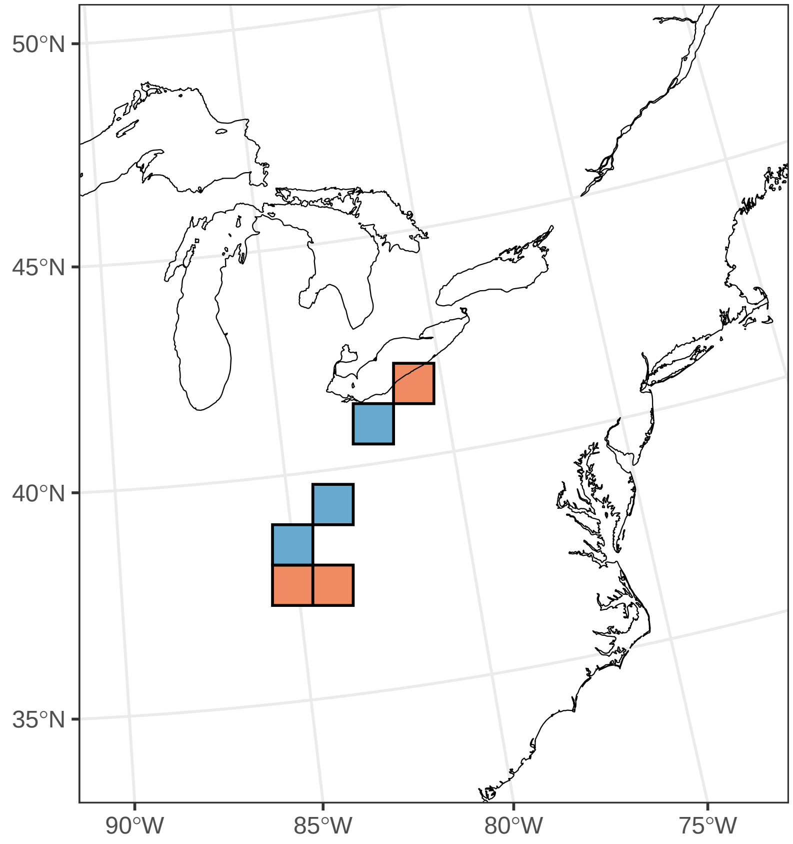

#### ***Cardamine angustata***

Ephemerality index: 1

n = 5

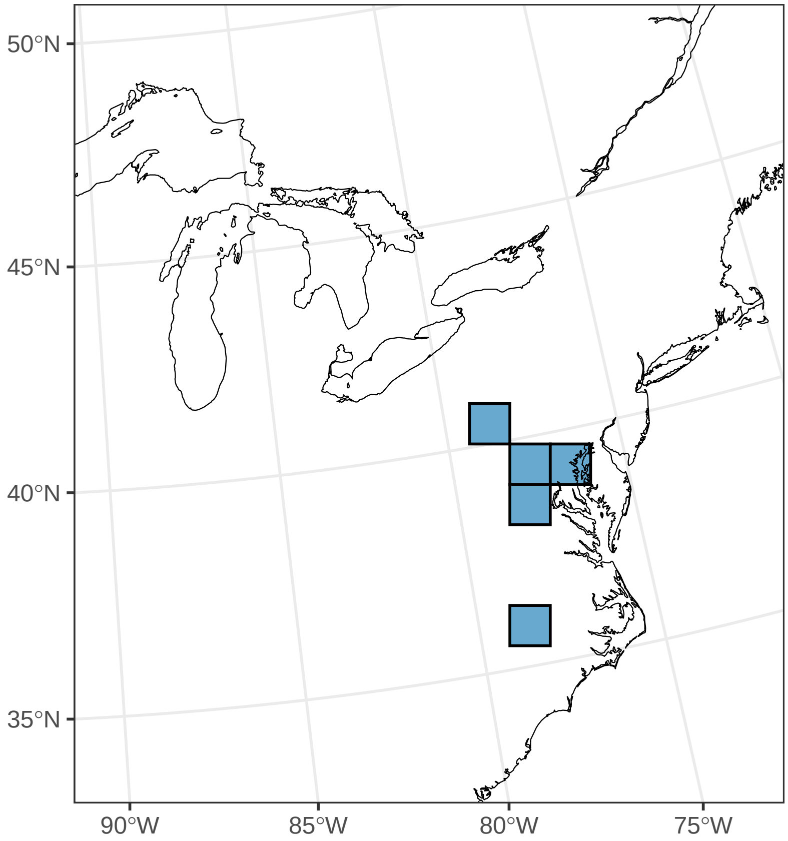

#### ***Cardamine bulbosa***

Ephemerality index: 0.875

n = 8

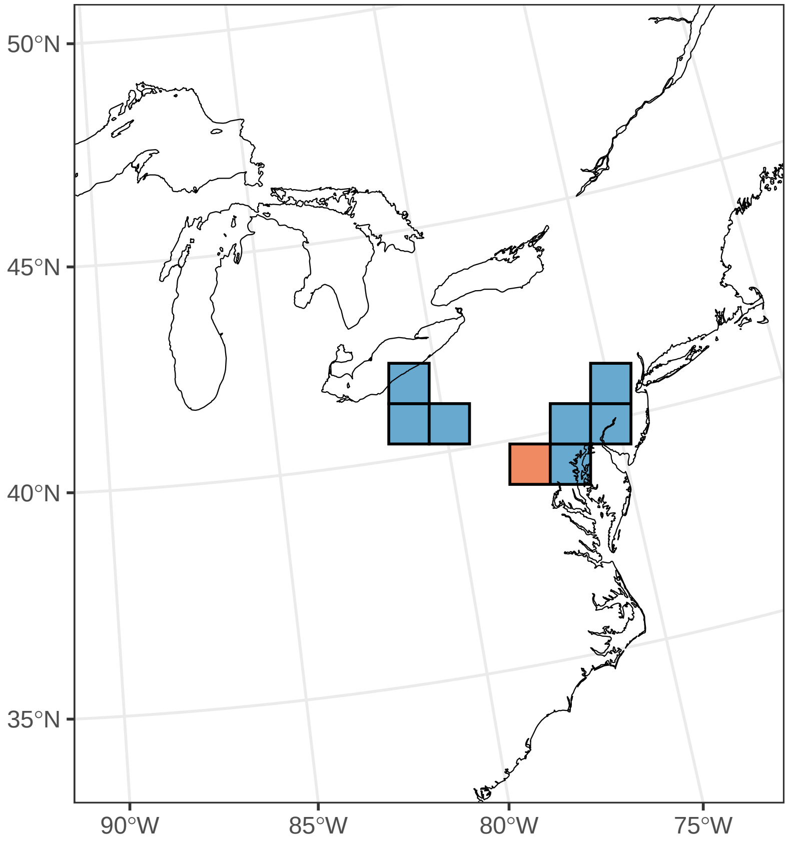

#### *Cardamine concatenata*

Ephemerality index: 0.982

n = 56

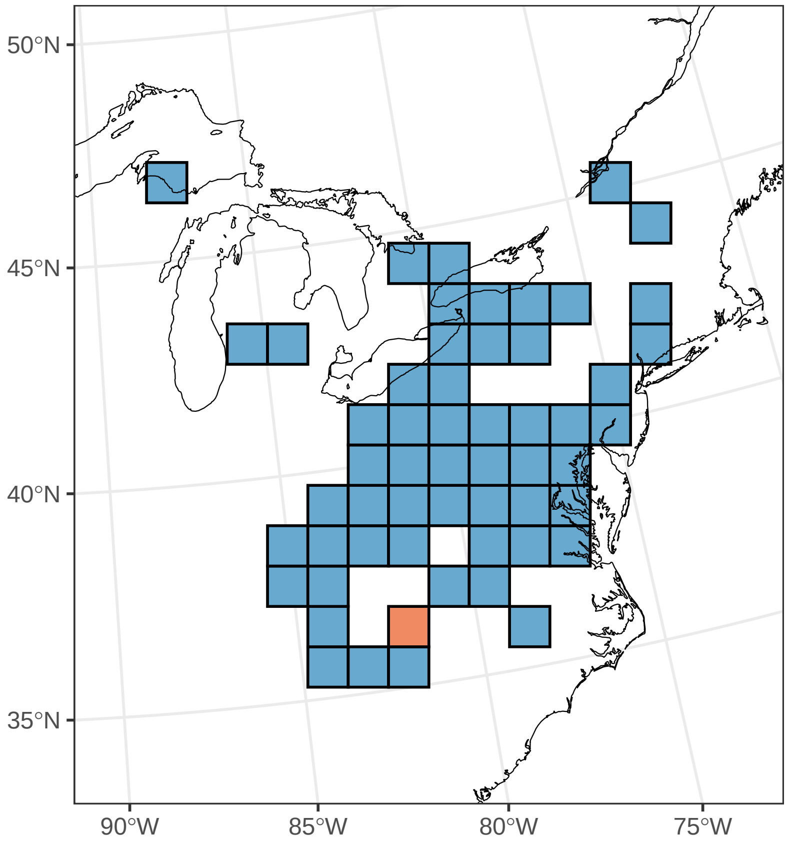

#### *Cardamine diphylla*

Ephemerality index: 0.566

n = 53

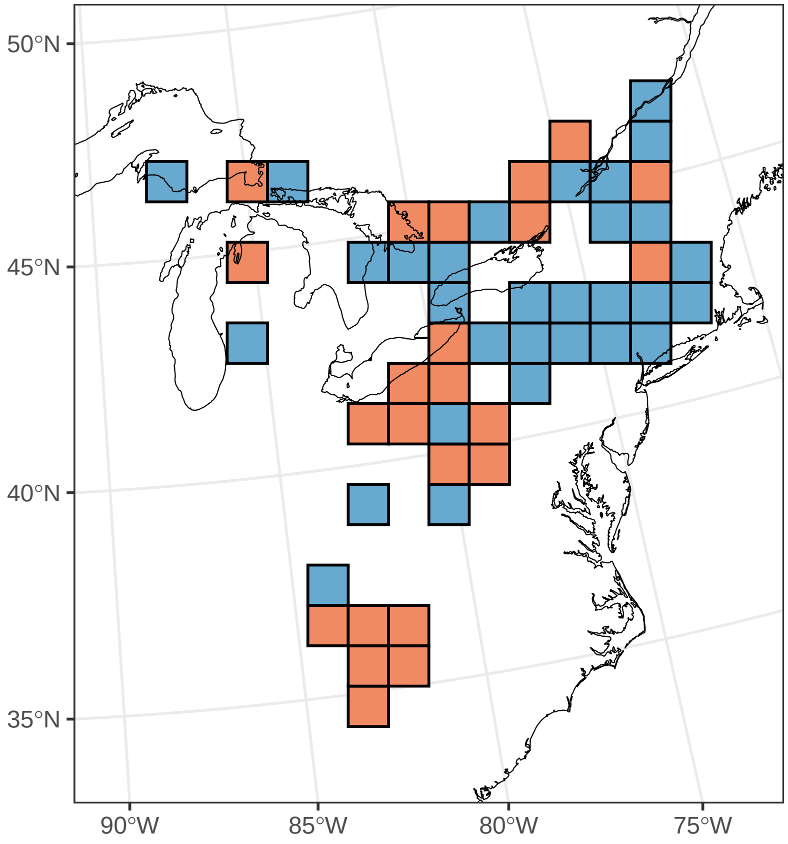

#### ***Cardamine douglassii***

Ephemerality index: 0.938

n = 16

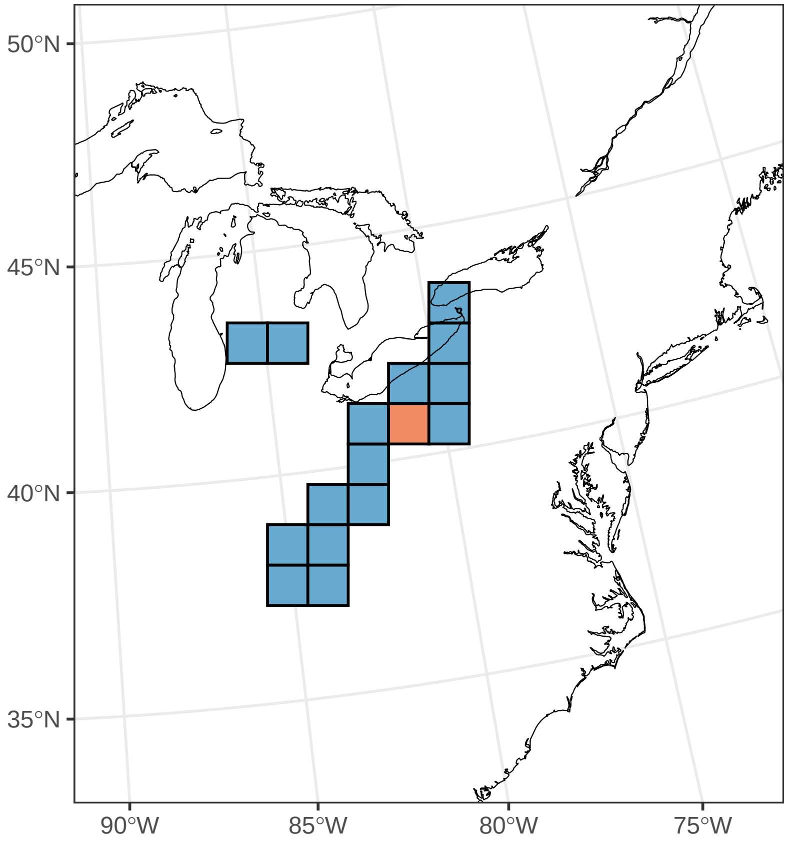

#### *Cardamine pensylvanica*

Ephemerality index: 0.333

n = 6

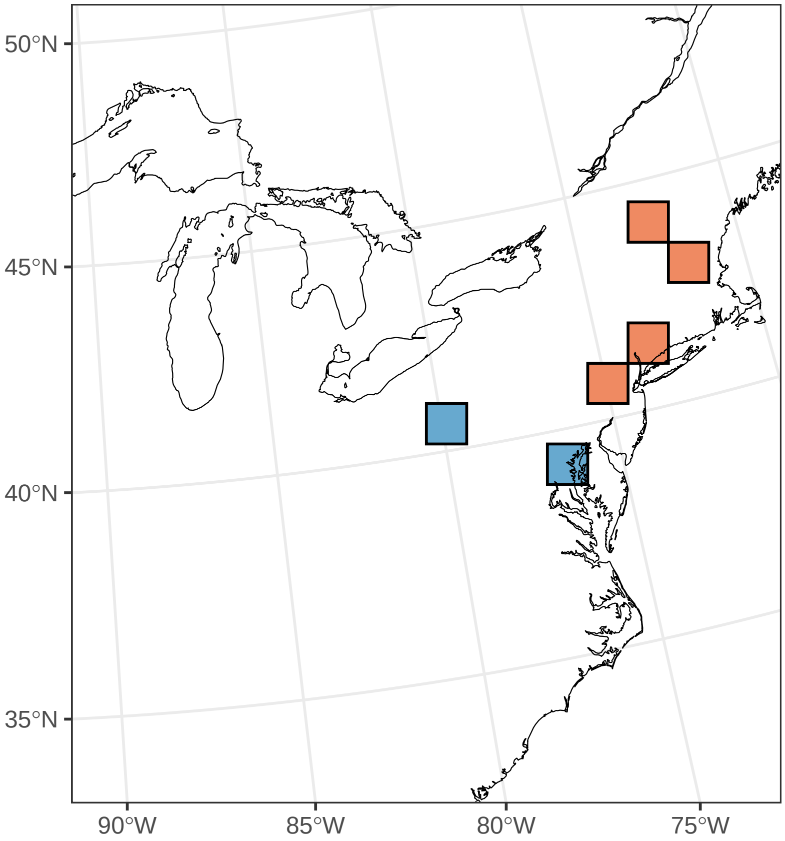

#### ***Caulophyllum giganteum***

Ephemerality index: 0.833

n = 30

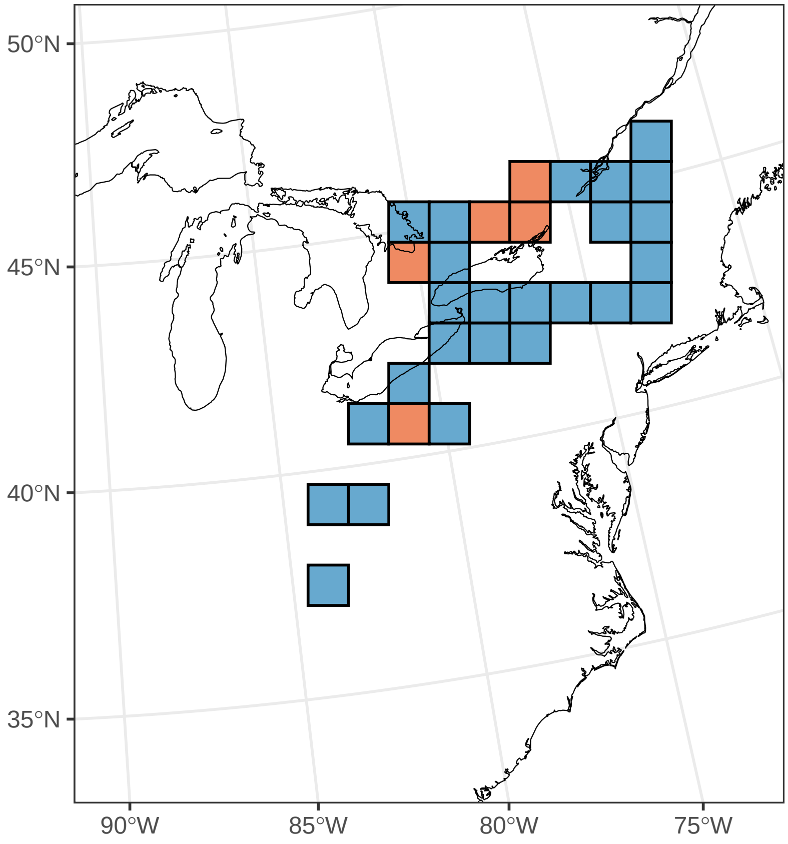

#### ***Claytonia caroliniana***

Ephemerality index: 0.764

n = 55

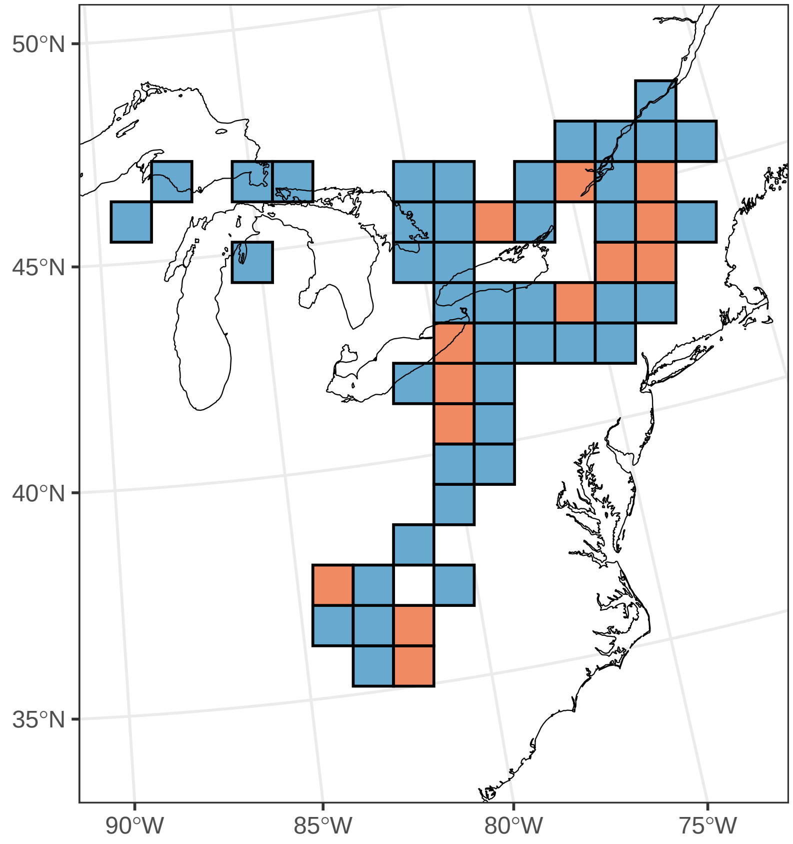

#### ***Collinsia verna***

Ephemerality index: 0.75

n = 8

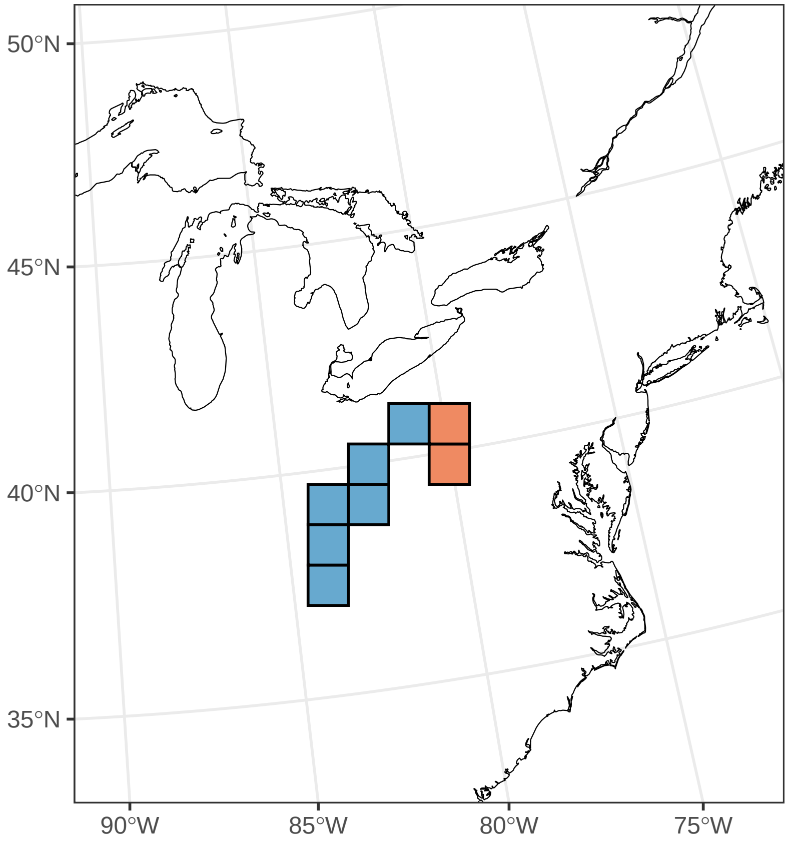

#### ***Convallaria majalis***

Ephemerality index: 0.032

n = 62

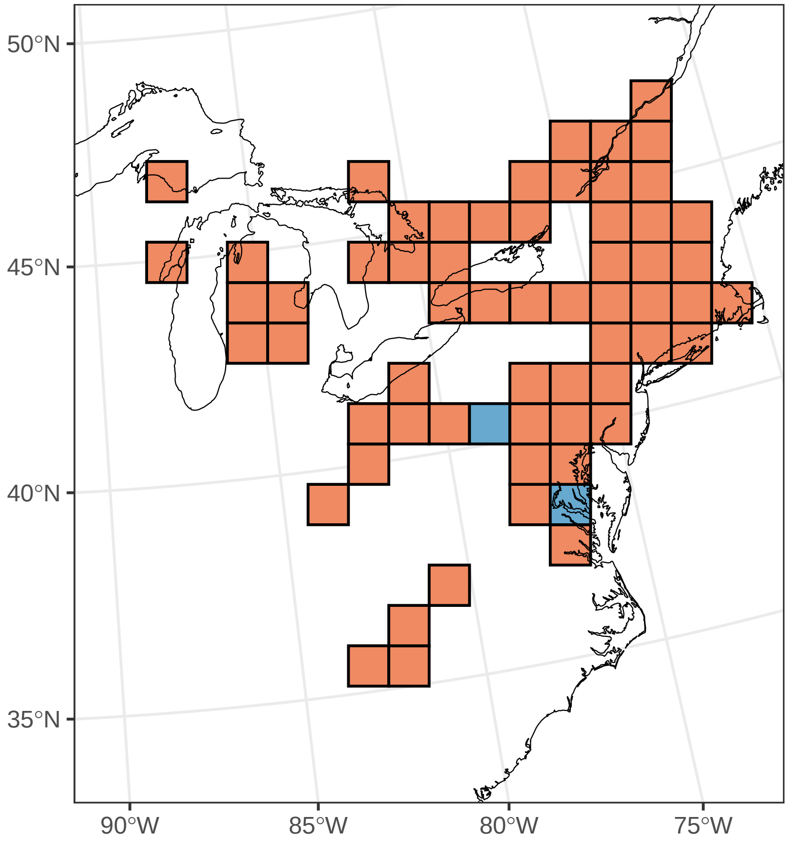

#### ***Corydalis flavula***

Ephemerality index: 0.667

n = 21

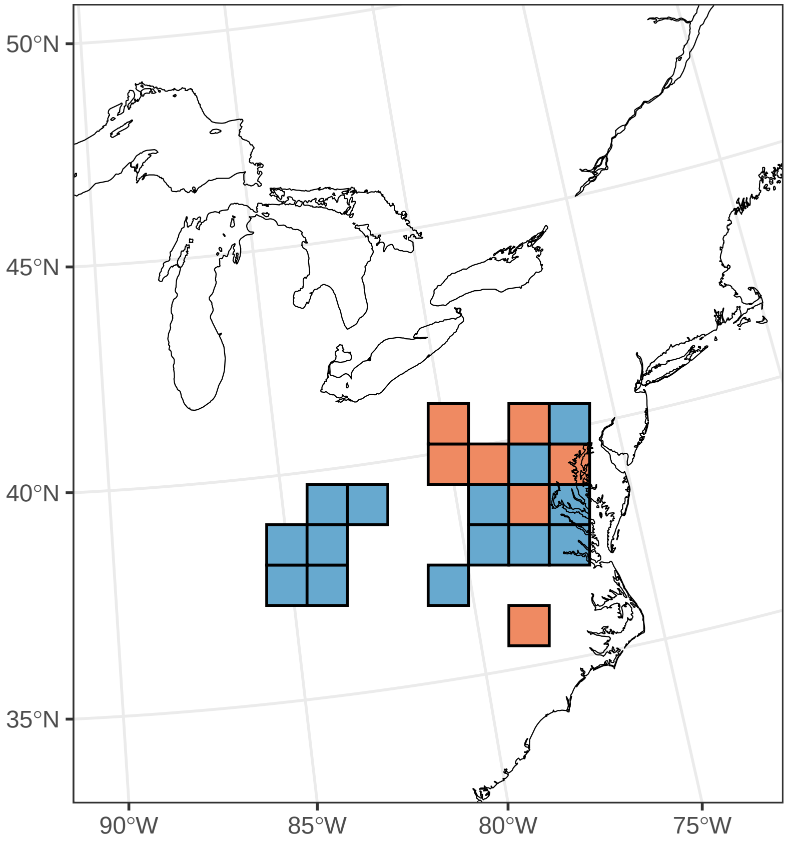

#### ***Cypripedium acaule***

Ephemerality index: 0.021

n = 97

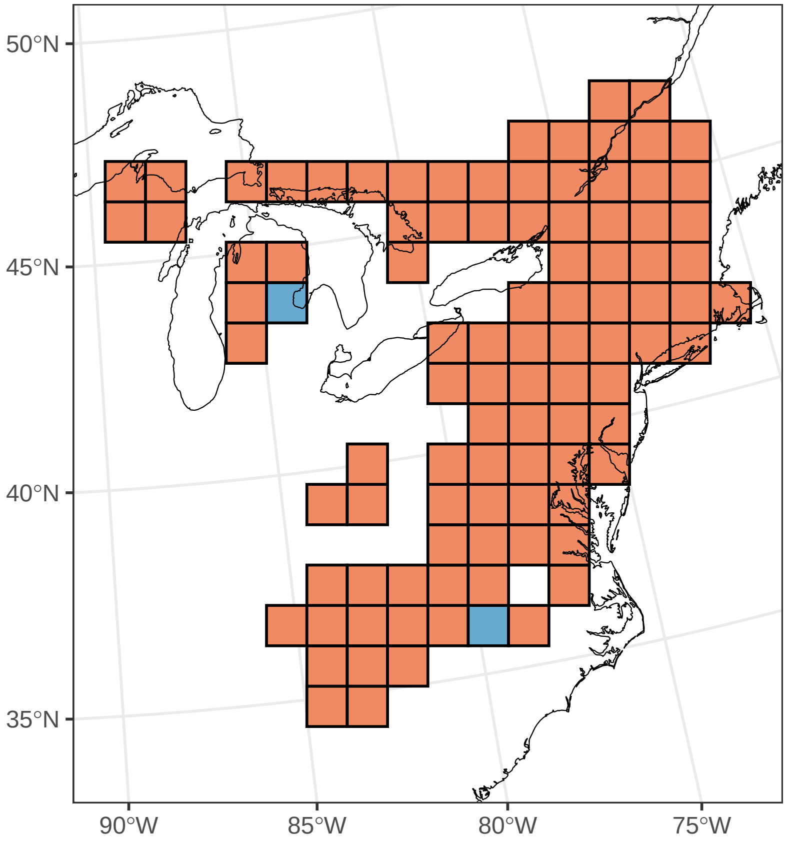

##### ***Cypripedium parviflorum***

Ephemerality index: 0.125

n = 32

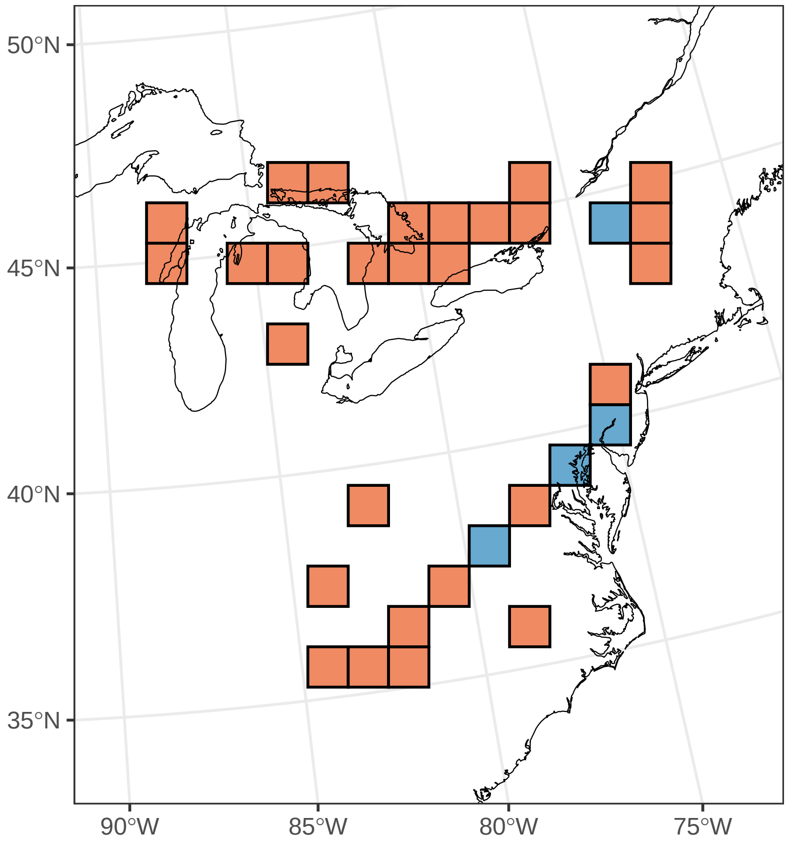

#### ***Dactylorhiza viridis***

Ephemerality index: 1

n = 1

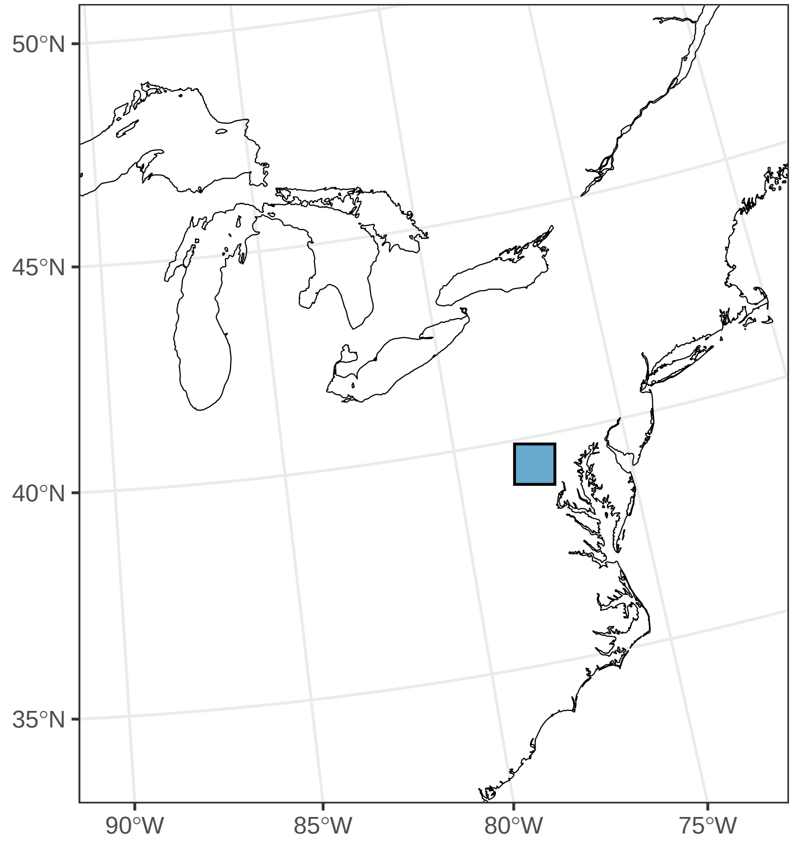

#### *Delphinium tricorne*

Ephemerality index: 0.8

n = 20

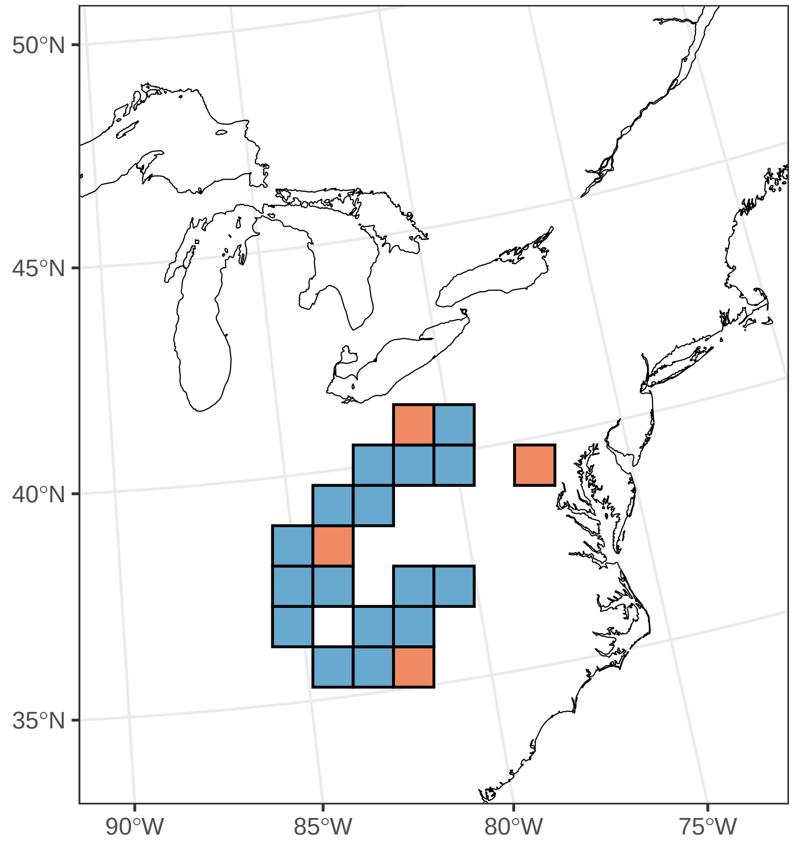

#### *Dicentra canadensis*

Ephemerality index: 0.722

n = 36

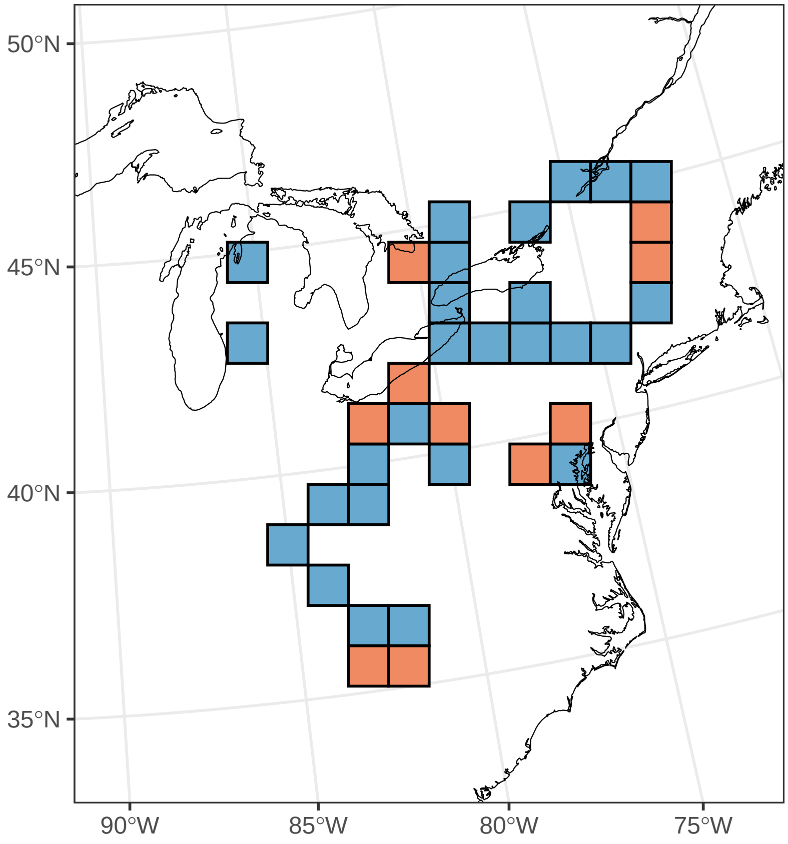

#### ***Dicentra cucullaria***

Ephemerality index: 0.537

n = 67

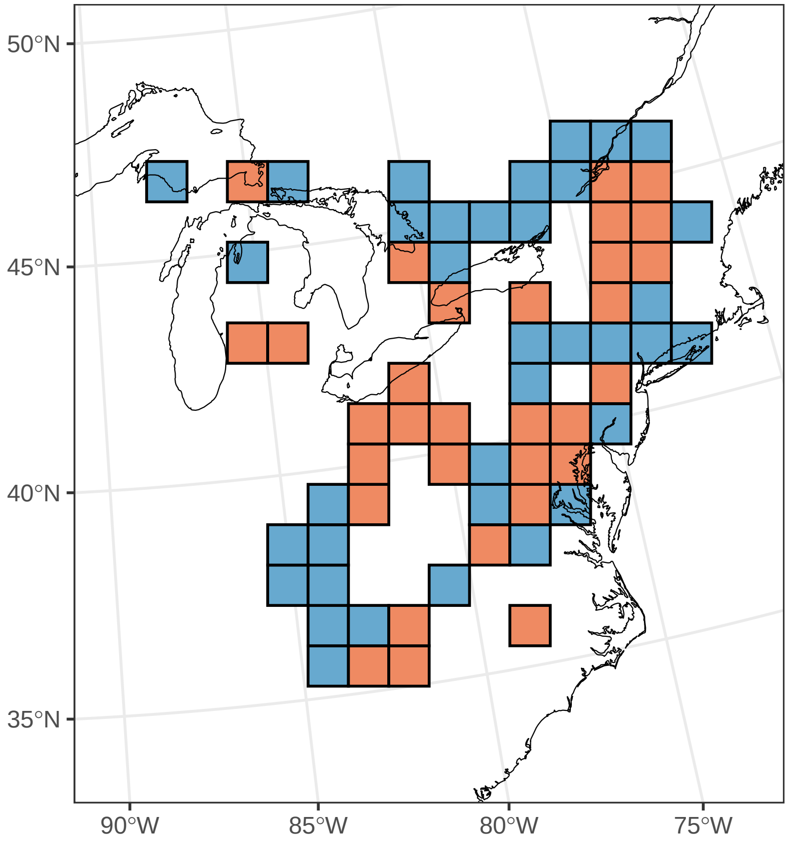

#### ***Erigenia bulbosa***

Ephemerality index: 0.462

n = 13

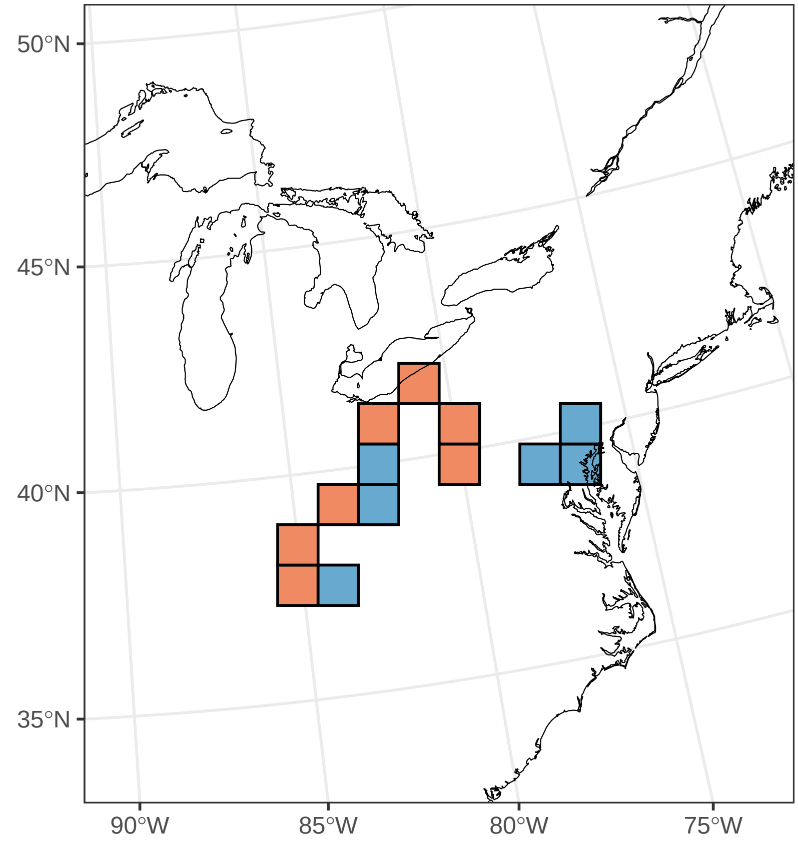

#### ***Erythronium albidum***

Ephemerality index: 0.929

n = 14

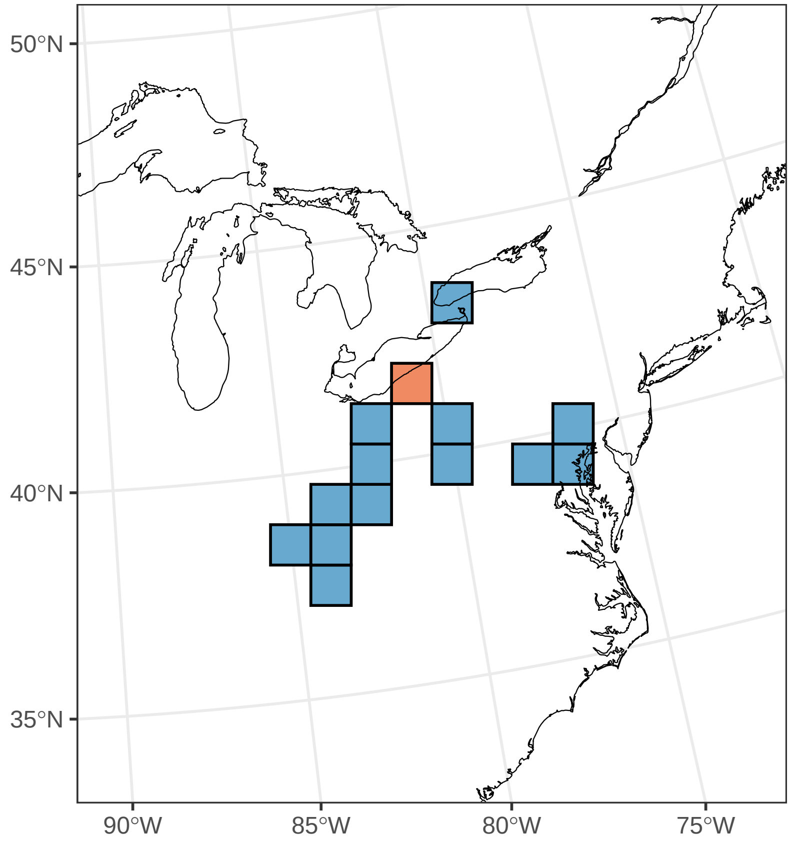

#### ***Erythronium americanum***

Ephemerality index: 0.883

n = 94

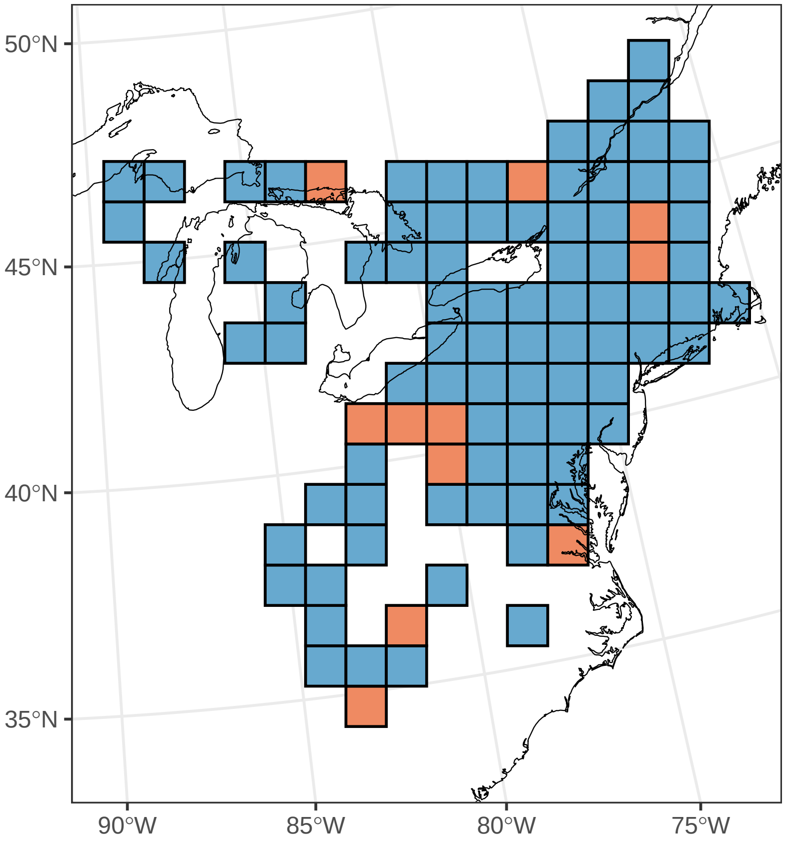

#### ***Erythronium umbilicatum***

Ephemerality index: 0.875

n = 8

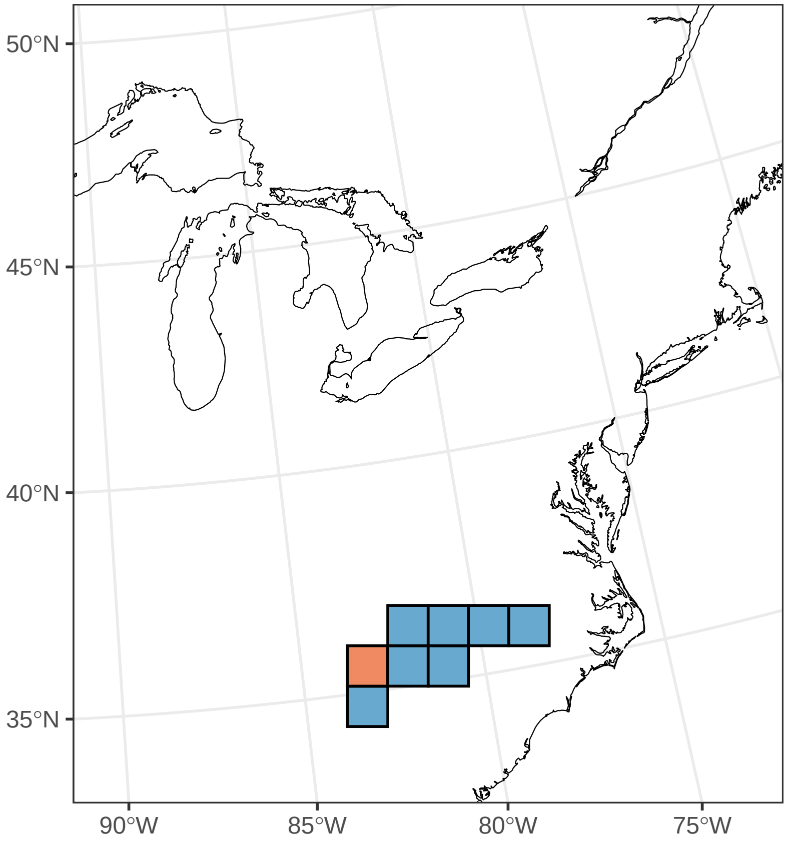

##### ***Euphorbia spathulata***

Ephemerality index: 1

n = 1

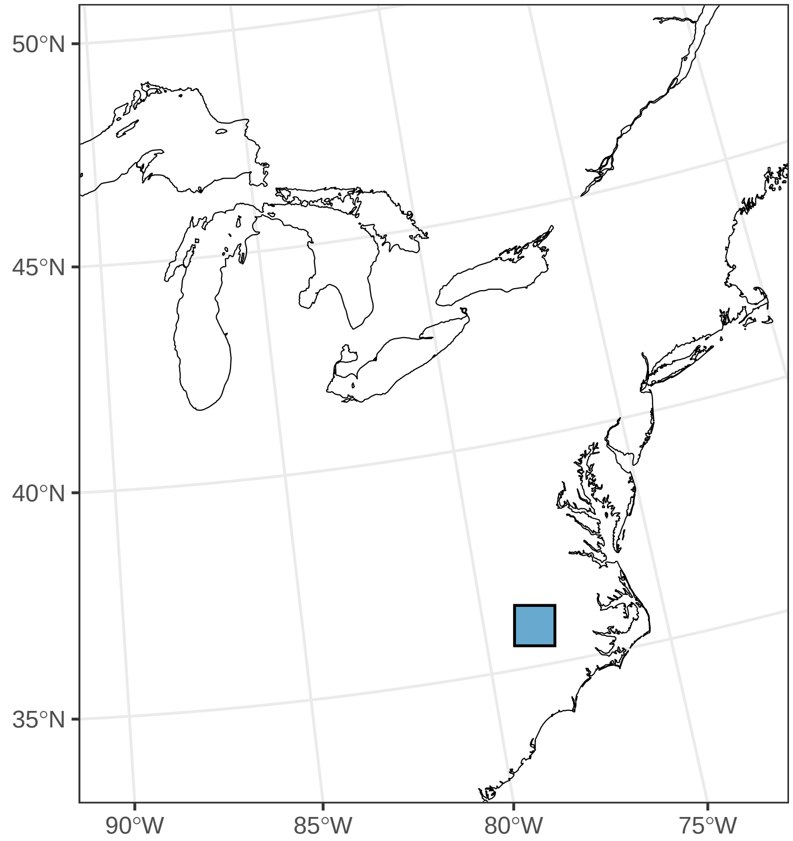

#### ***Floerkea proserpinacoides***

Ephemerality index: 0.875

n = 8

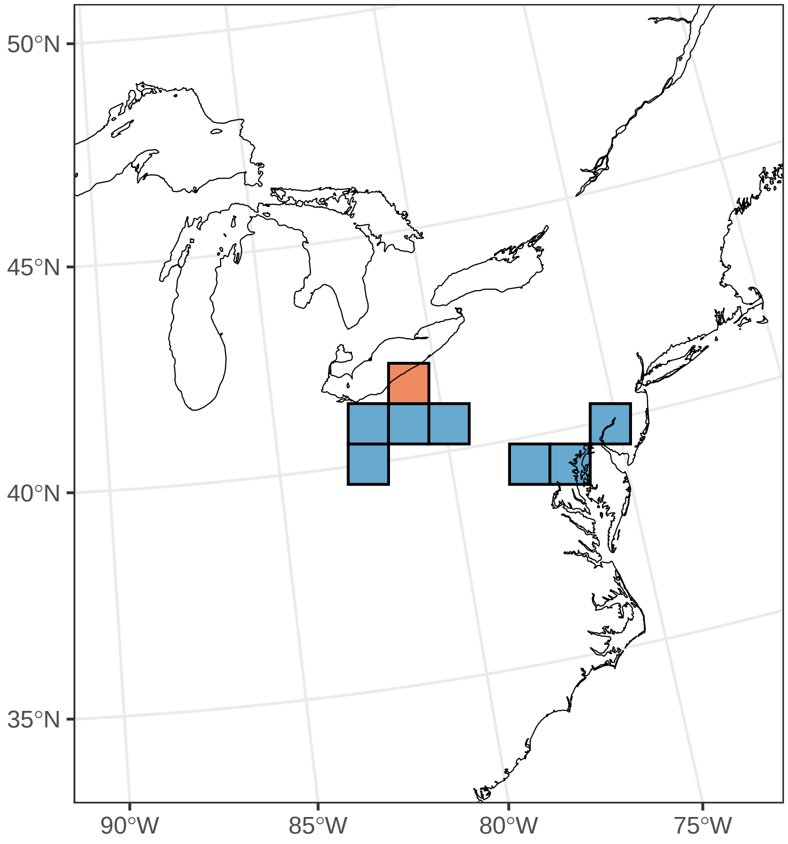

#### ***Galearis spectabilis***

Ephemerality index: 0.065

n = 31

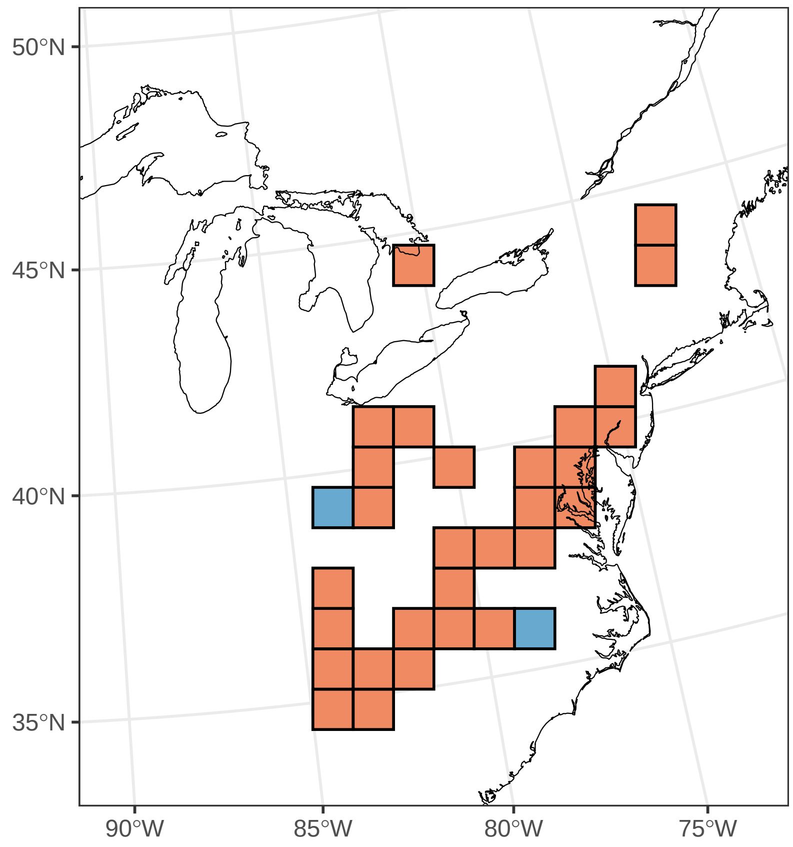

#### ***Hybanthus concolor***

Ephemerality index: 0.4

n = 5

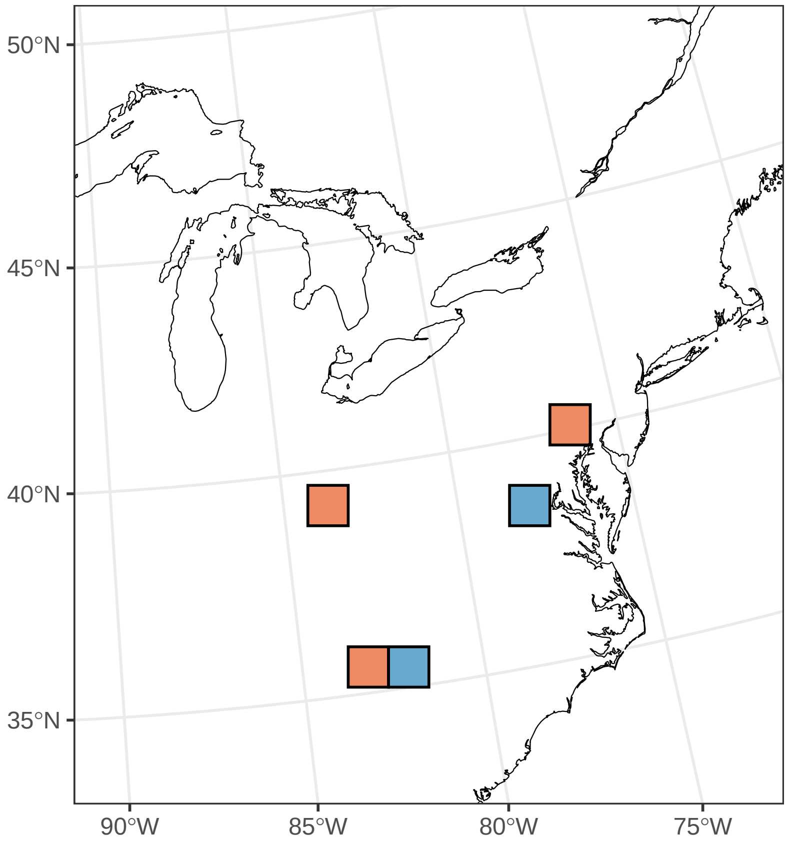

#### ***Hydrophyllum appendiculatum***

Ephemerality index: 0.143

n = 7

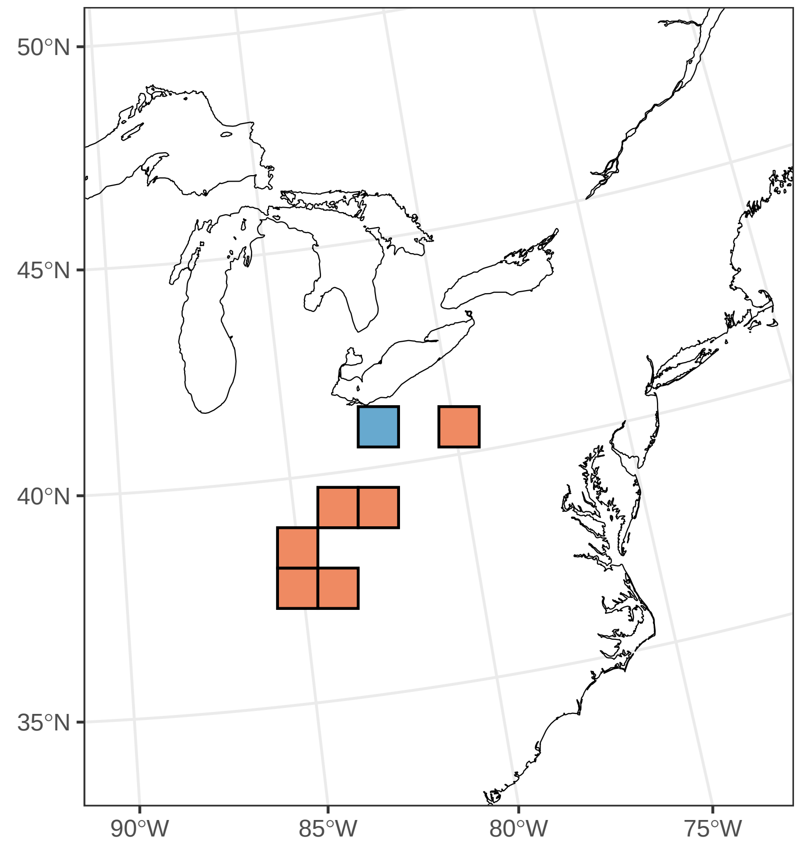

#### ***Hydrophyllum virginianum***

Ephemerality index: 0.024

n = 41

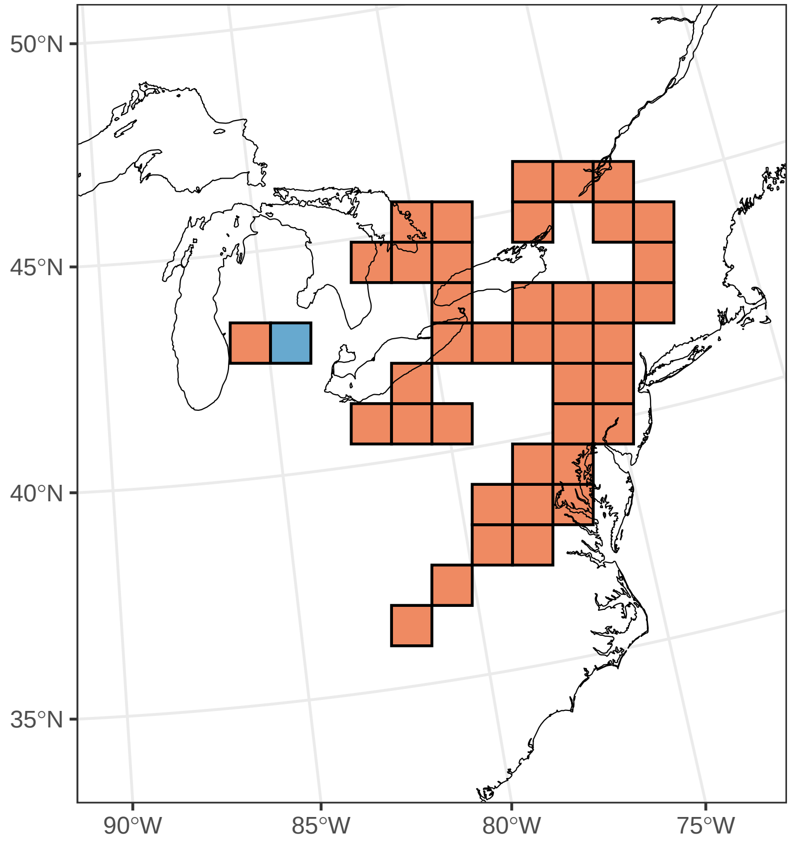

#### ***Hypoxis hirsuta***

Ephemerality index: 0.029

n = 35

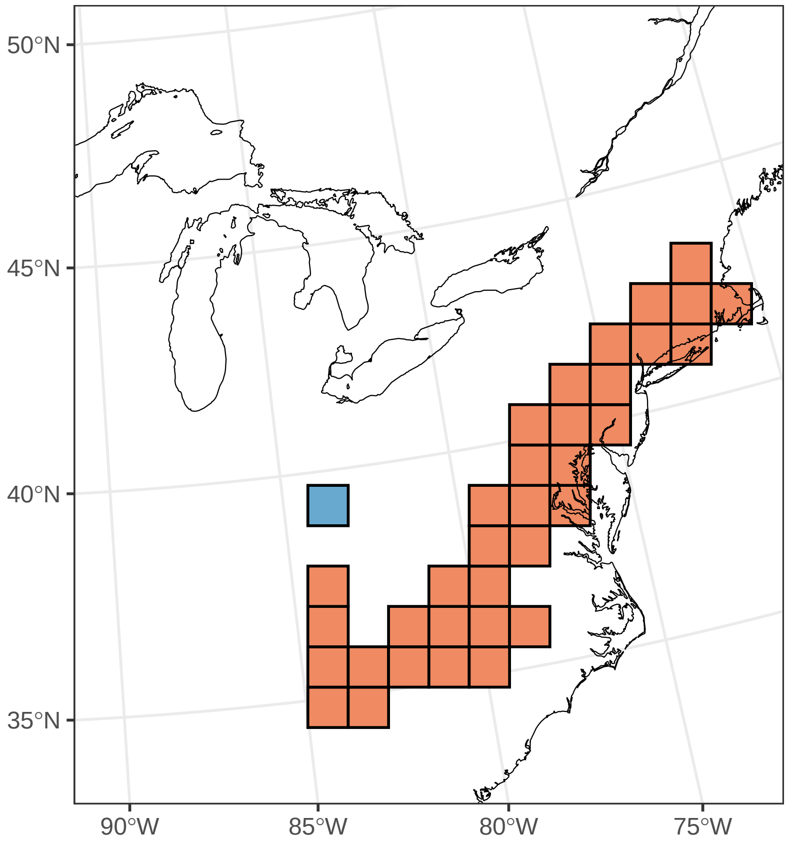

#### ***Iris cristata***

Ephemerality index: 0.174

n = 23

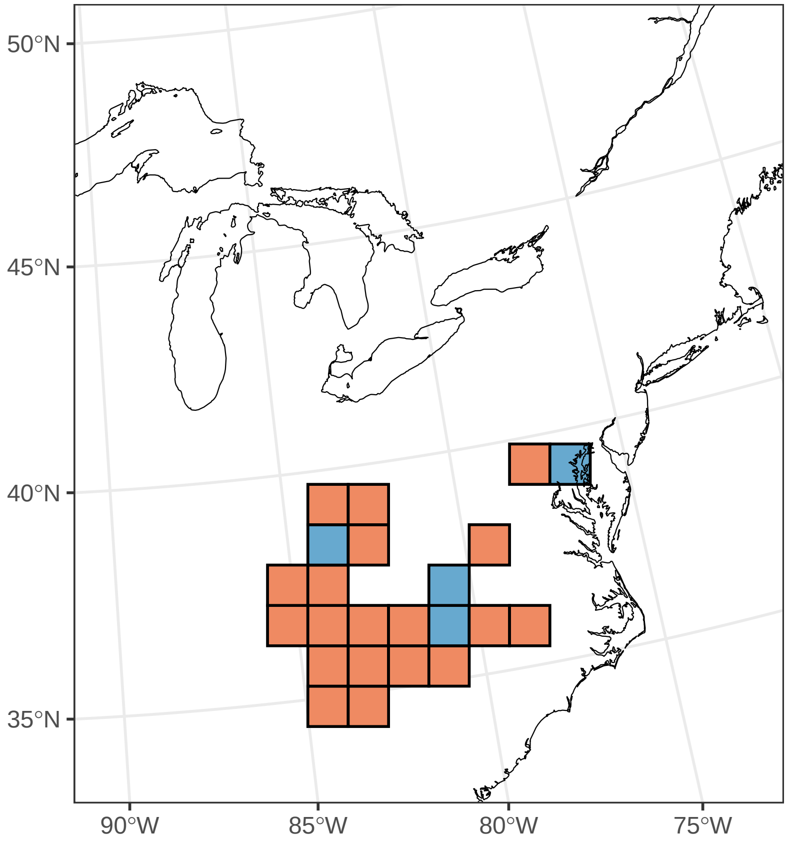

#### ***Iris verna***

Ephemerality index: 0.562

n = 16

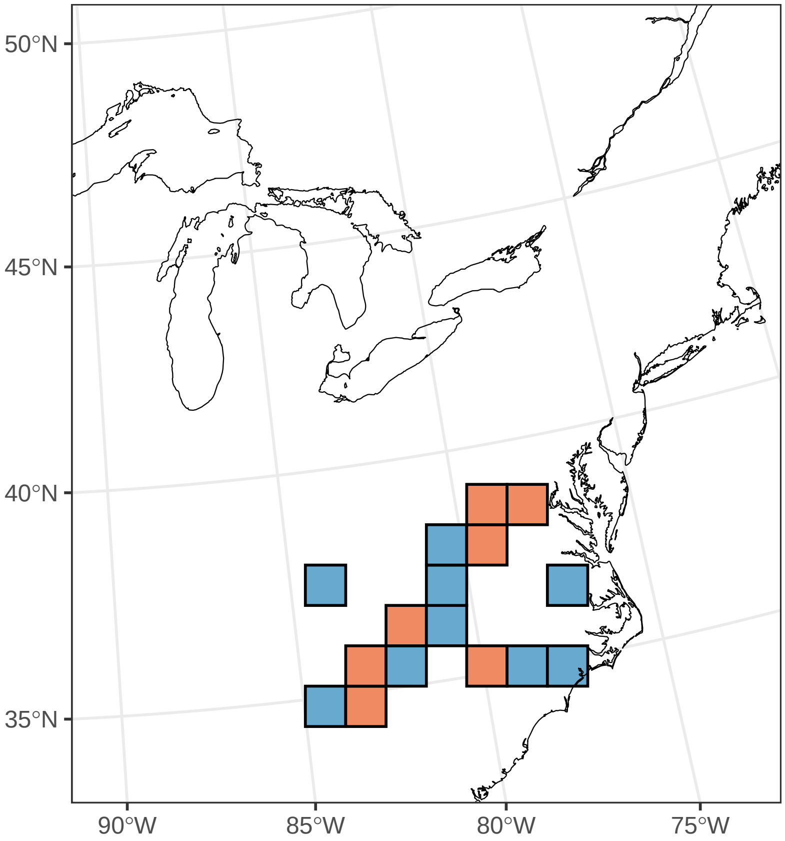

#### ***Jeffersonia diphylla***

Ephemerality index: 0.2

n = 20

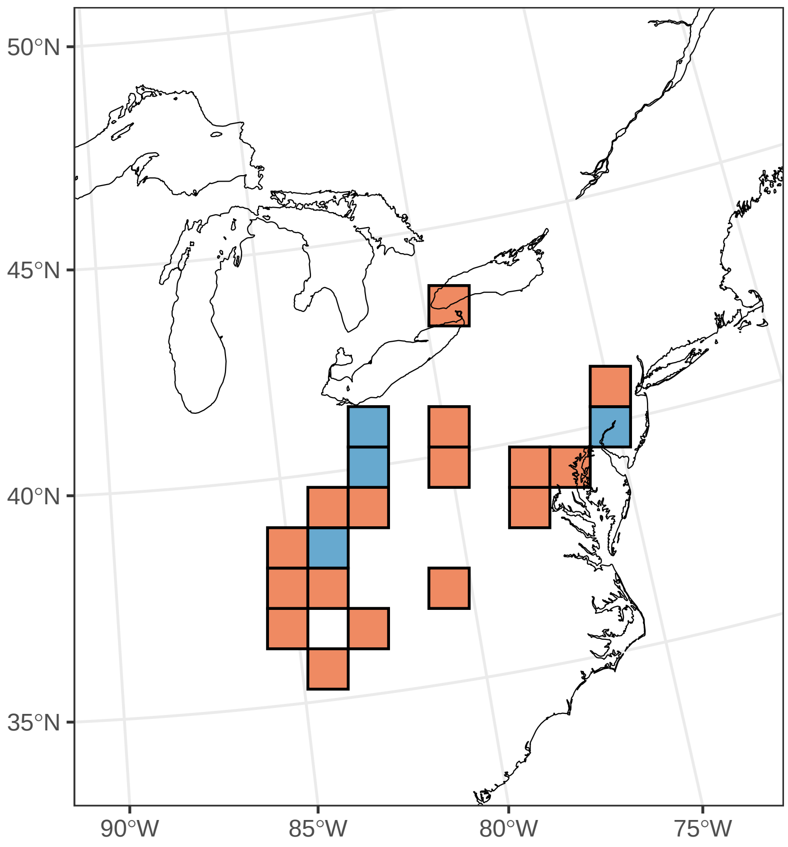

#### ***Krigia biflora***

Ephemerality index: 0.143

n = 7

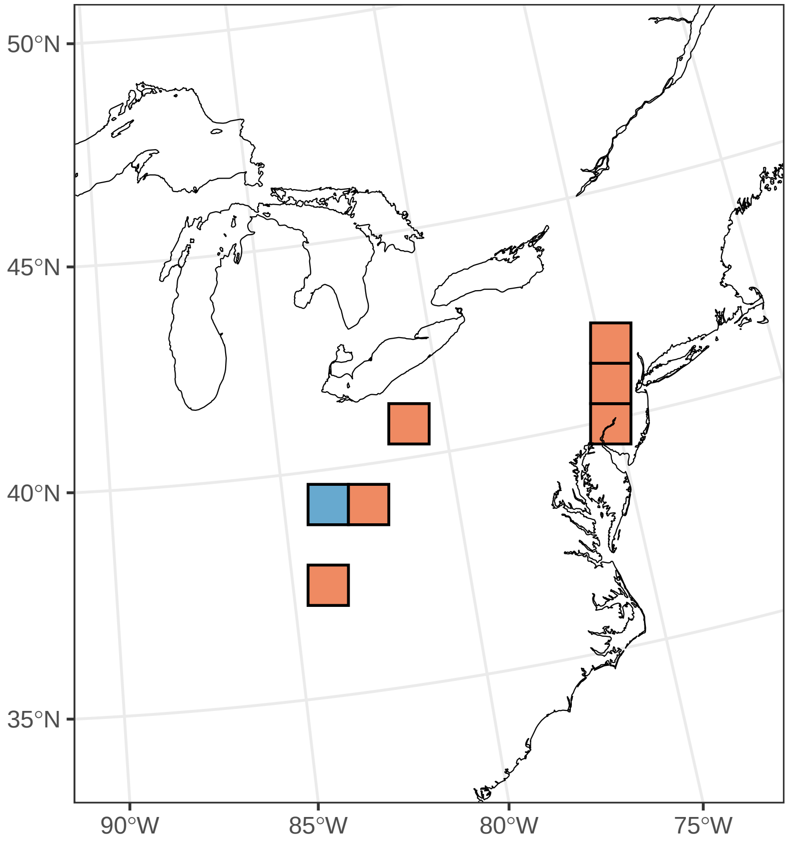

#### ***Krigia dandelion***

Ephemerality index: 1

n = 3

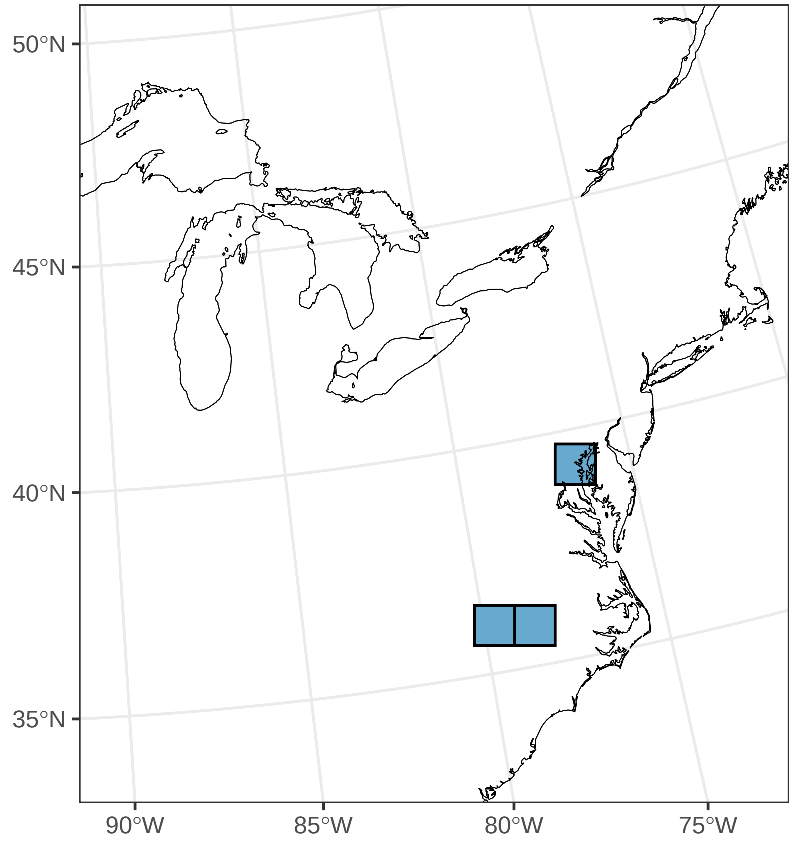

#### ***Krigia virginica***

Ephemerality index: 0.25

n = 8

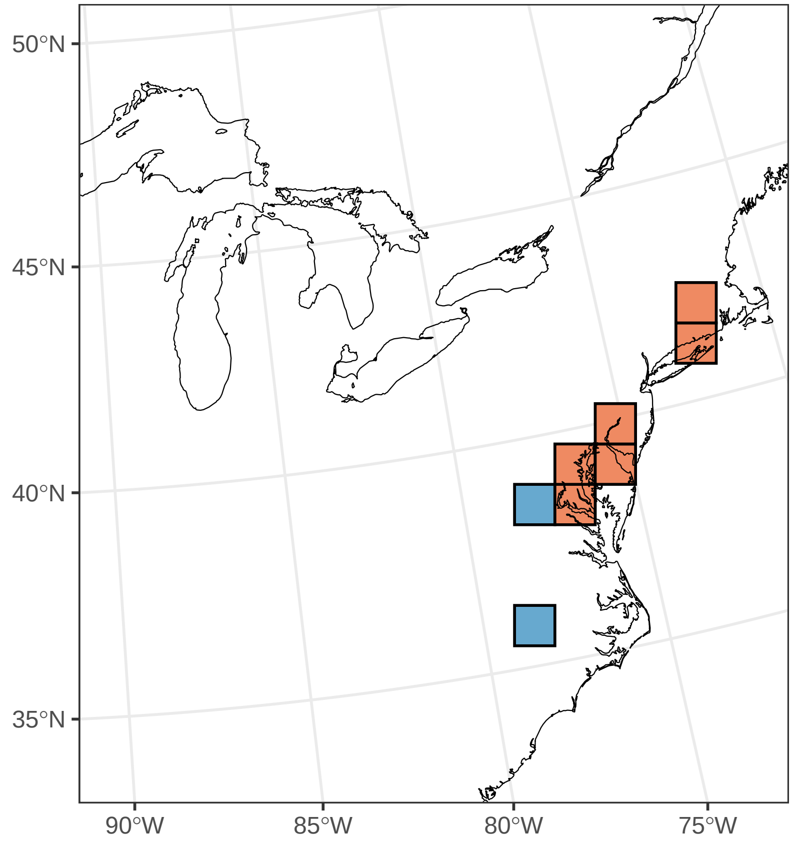

#### ***Lepidium campestre***

Ephemerality index: 0.154

n = 13

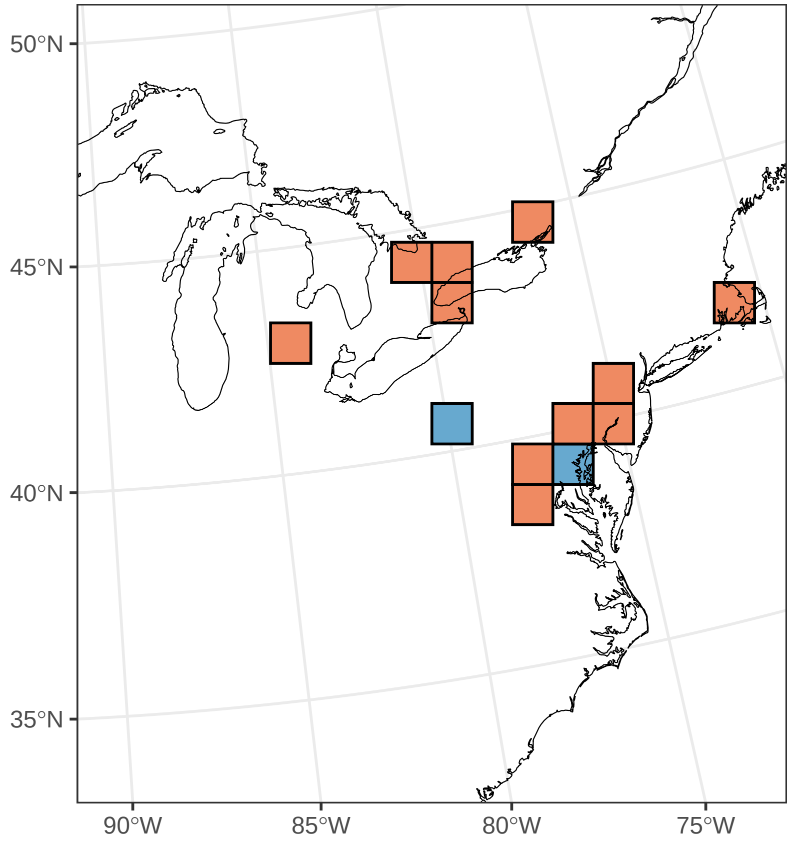

#### ***Lithospermum canescens***

Ephemerality index: 0.333

n = 3

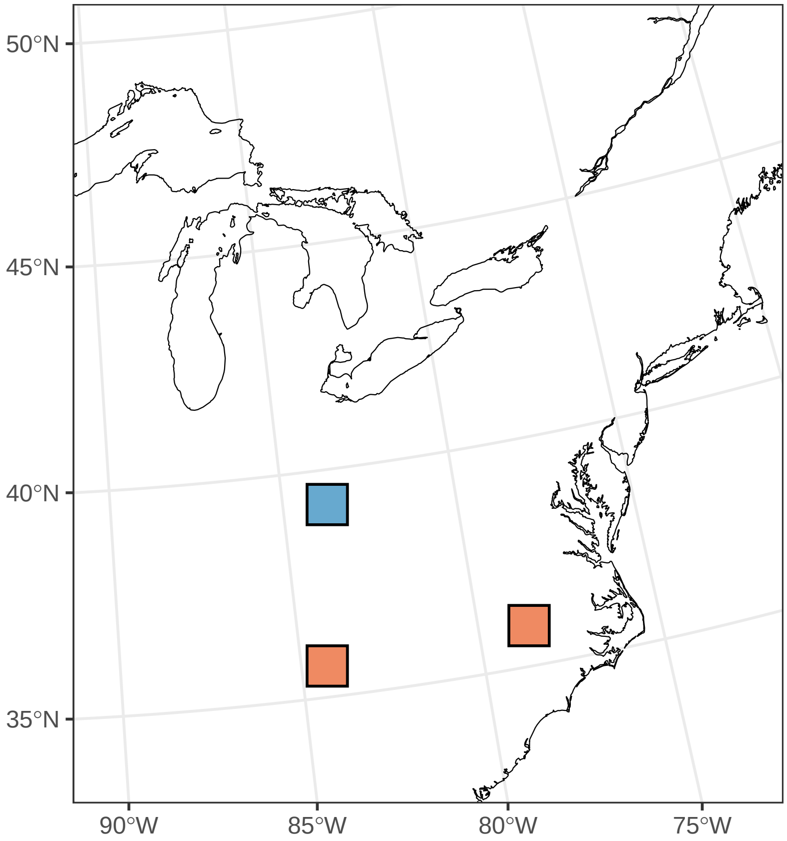

#### ***Lunaria annua***

Ephemerality index: 0.029

n = 35

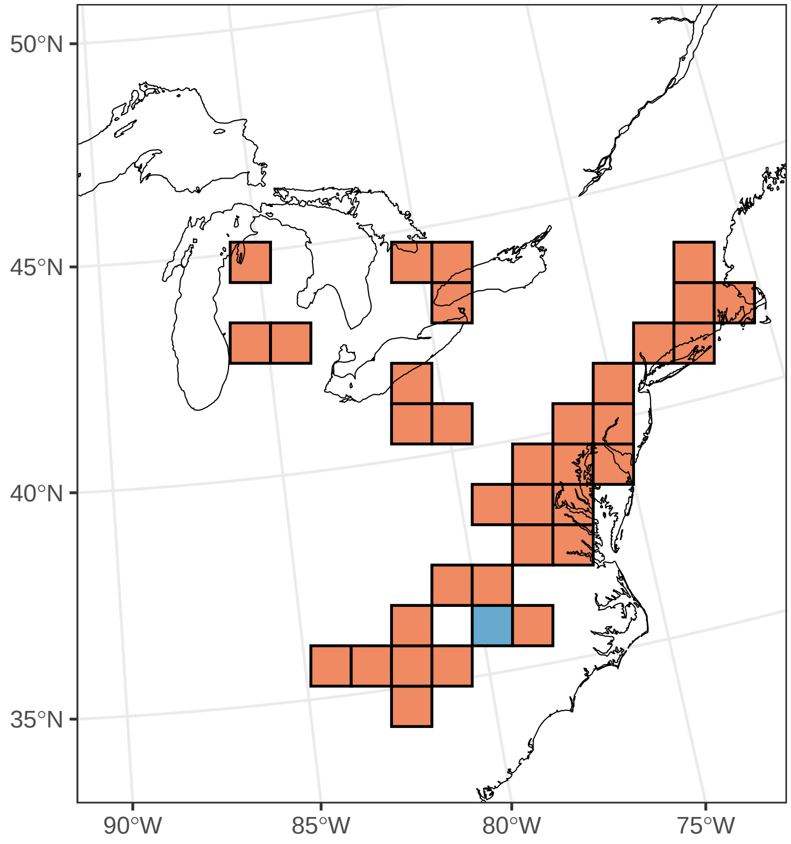

#### *Mertensia virginica*

Ephemerality index: 0.904

n = 52

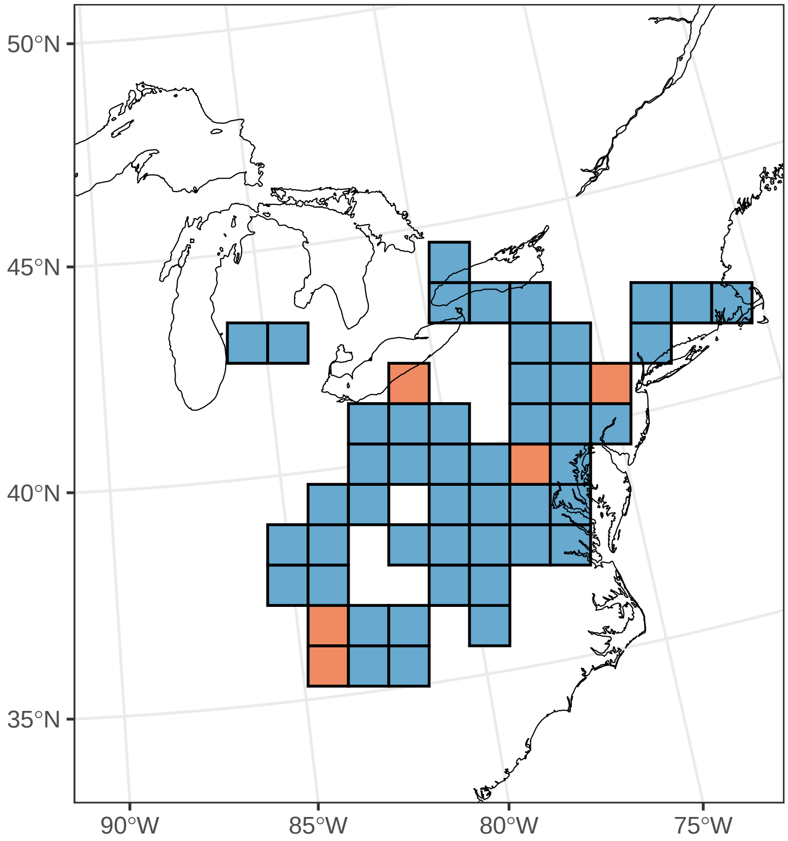

#### ***Muscari botryoides***

Ephemerality index: 1

n = 5

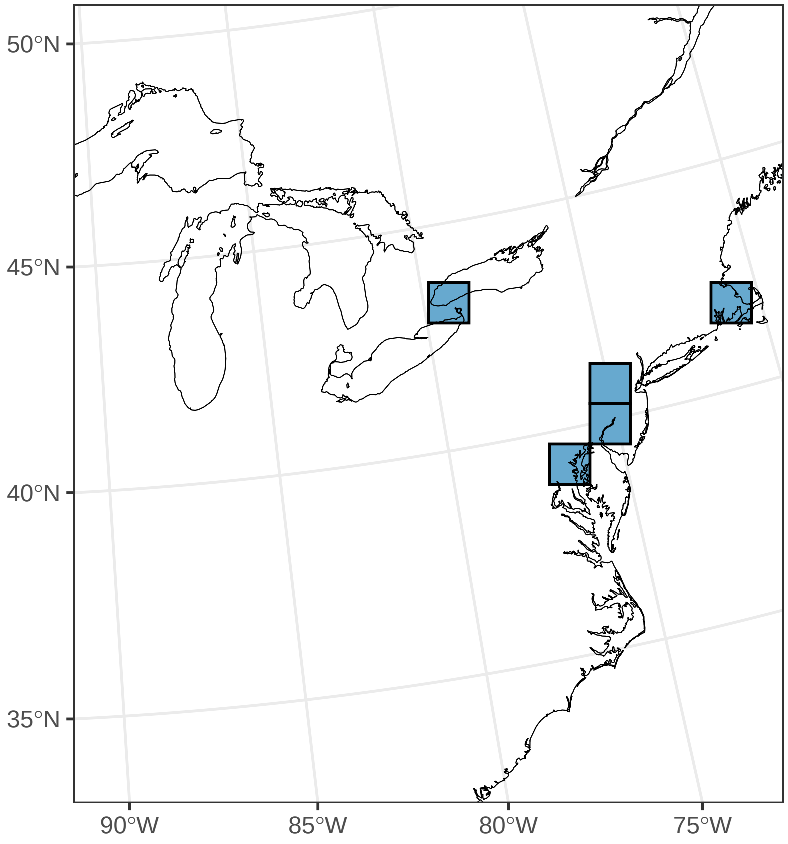

#### *Muscari neglectum*

Ephemerality index: 0.667

n = 3

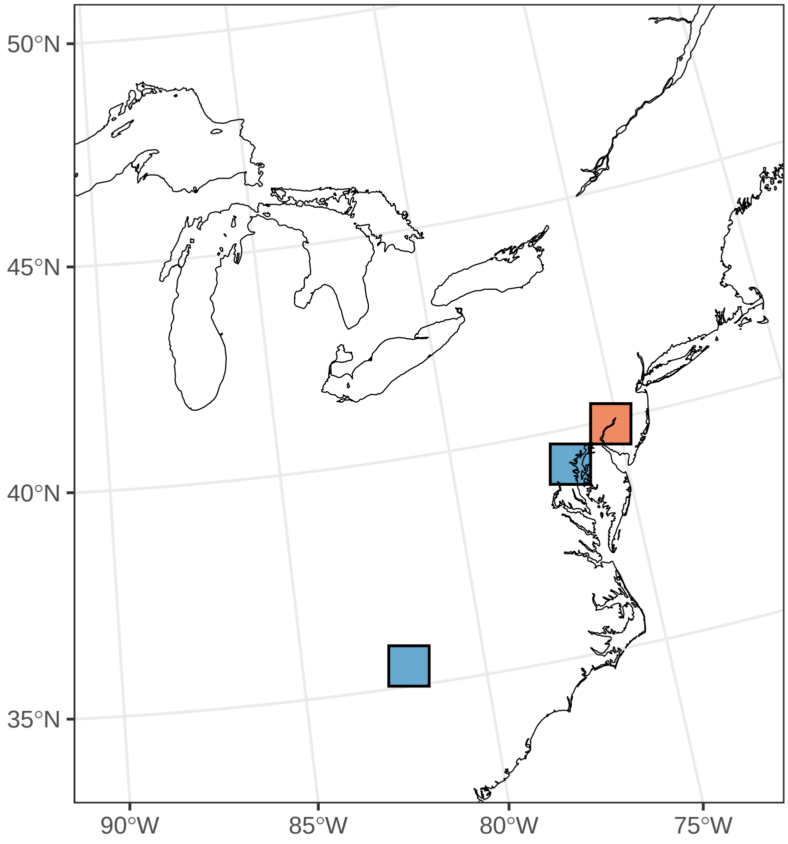

#### ***Myosotis macrosperma***

Ephemerality index: 1

n = 1

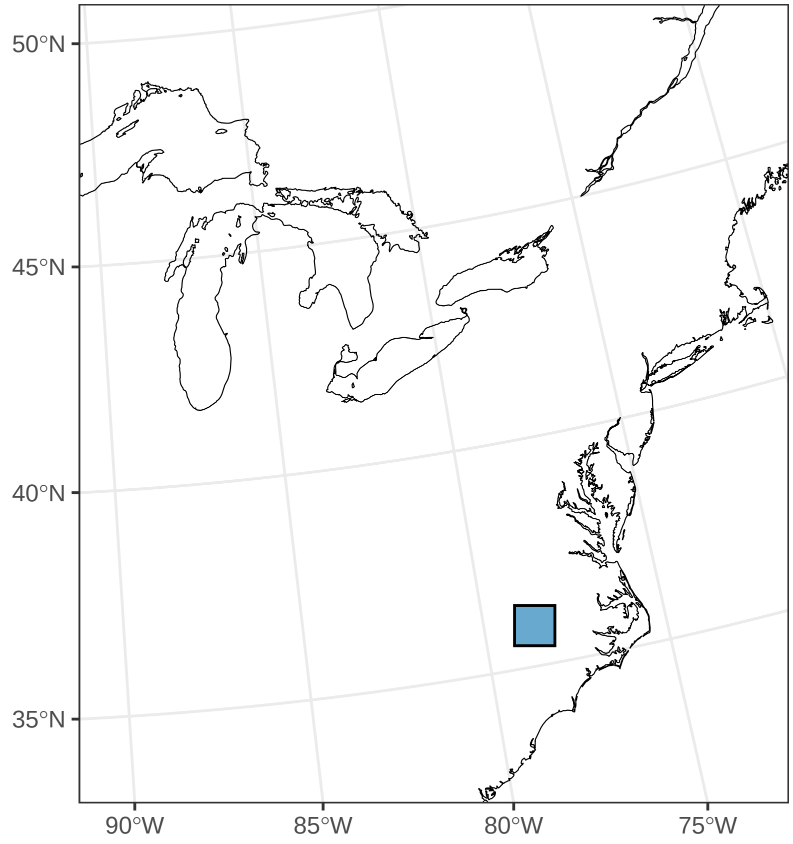

##### ***Narcissus pseudonarcissus***

Ephemerality index: 1

n = 3

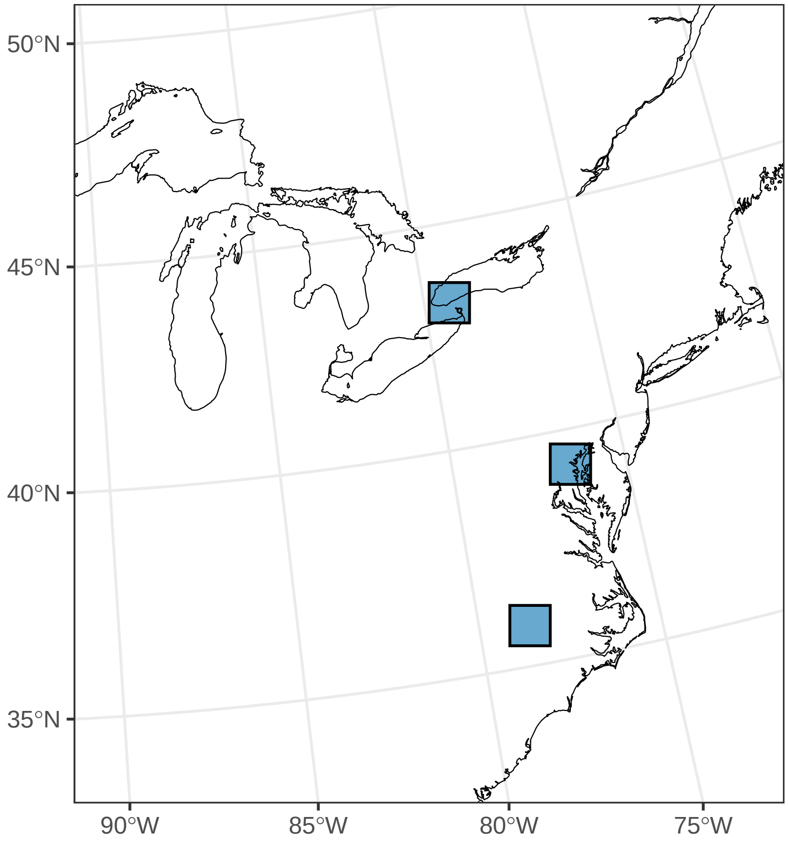

#### ***Obolaria virginica***

Ephemerality index: 0.4

n = 20

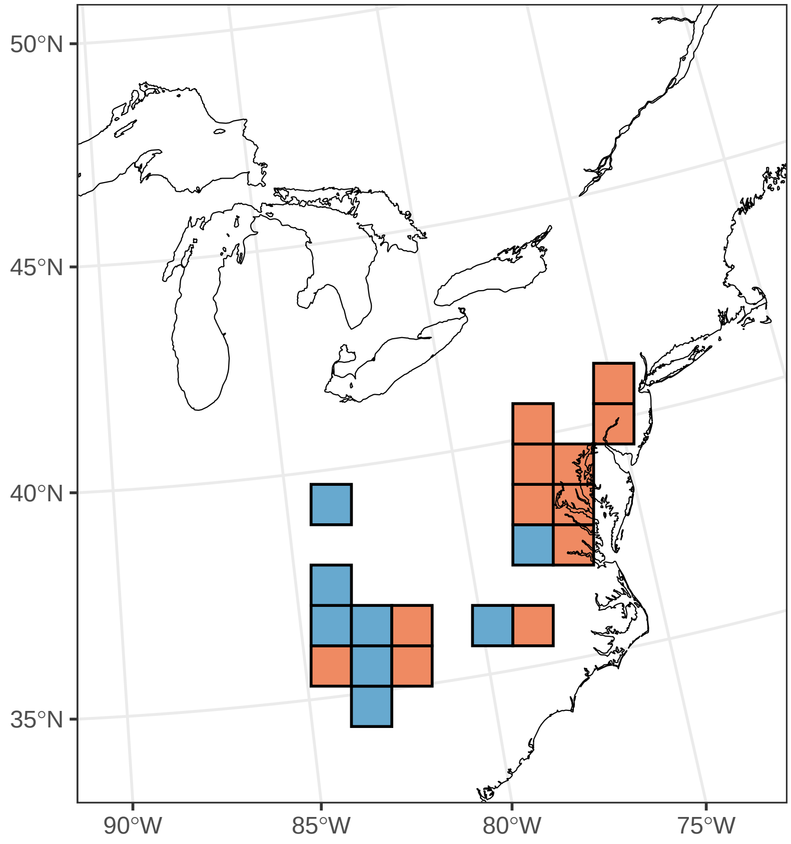

#### ***Ornithogalum umbellatum***

Ephemerality index: 0.902

n = 51

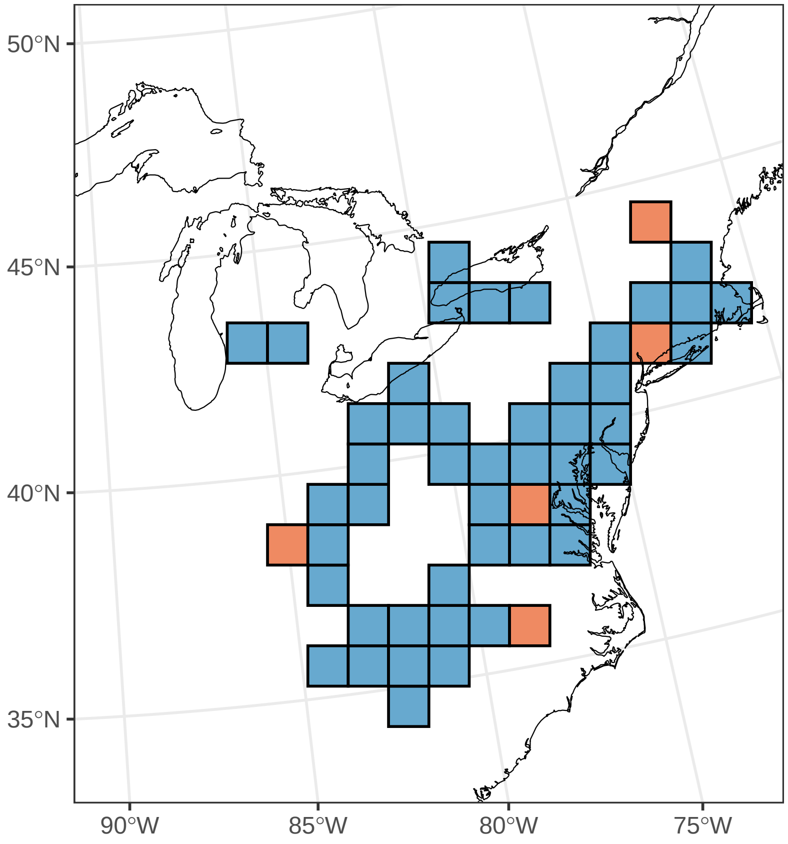

#### *Osmorhiza longistylis*

Ephemerality index: 0.333

n = 15

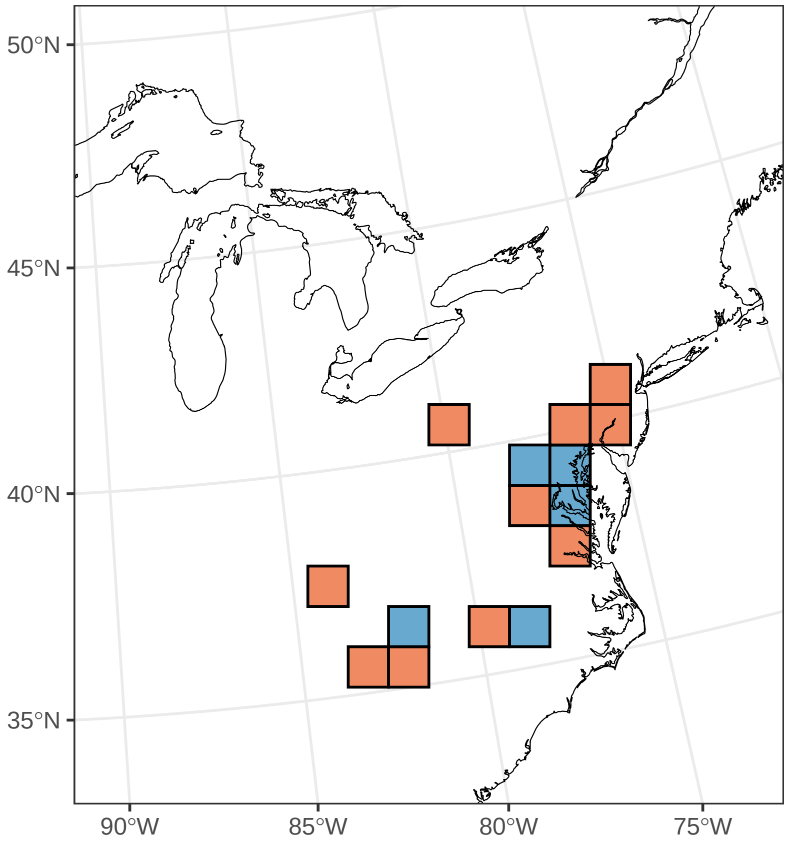

#### *Oxalis violacea*

Ephemerality index: 0.303

n = 33

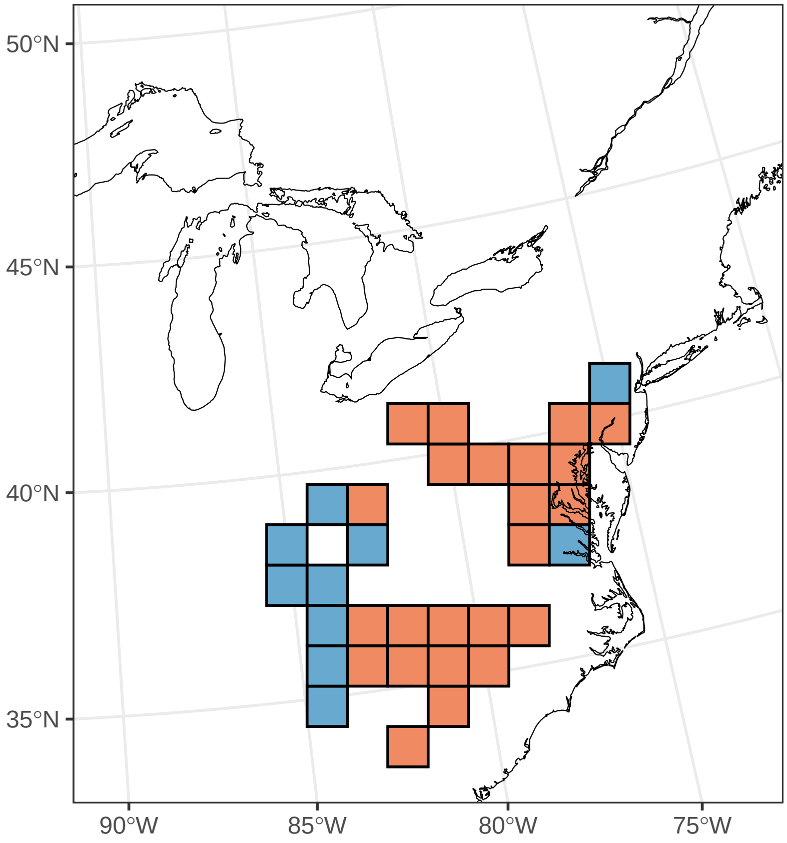

##### *Phacelia bipinnatifida*

Ephemerality index: 0.444

n = 9

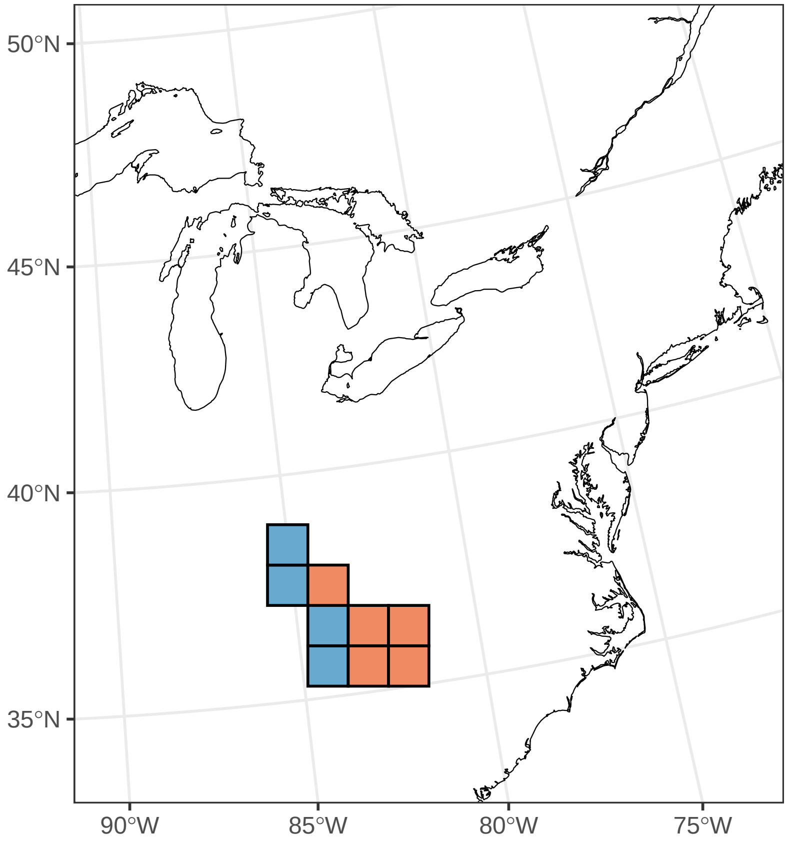

#### ***Phacelia purshii***

Ephemerality index: 0.75

n = 12

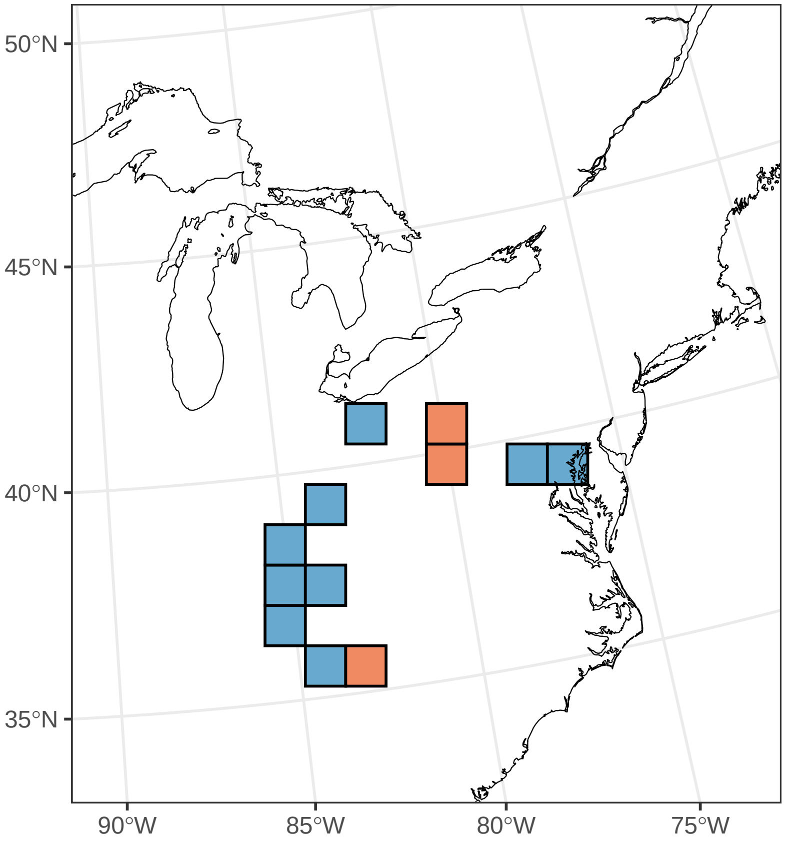

#### ***Phlox amoena***

Ephemerality index: 1

n = 1

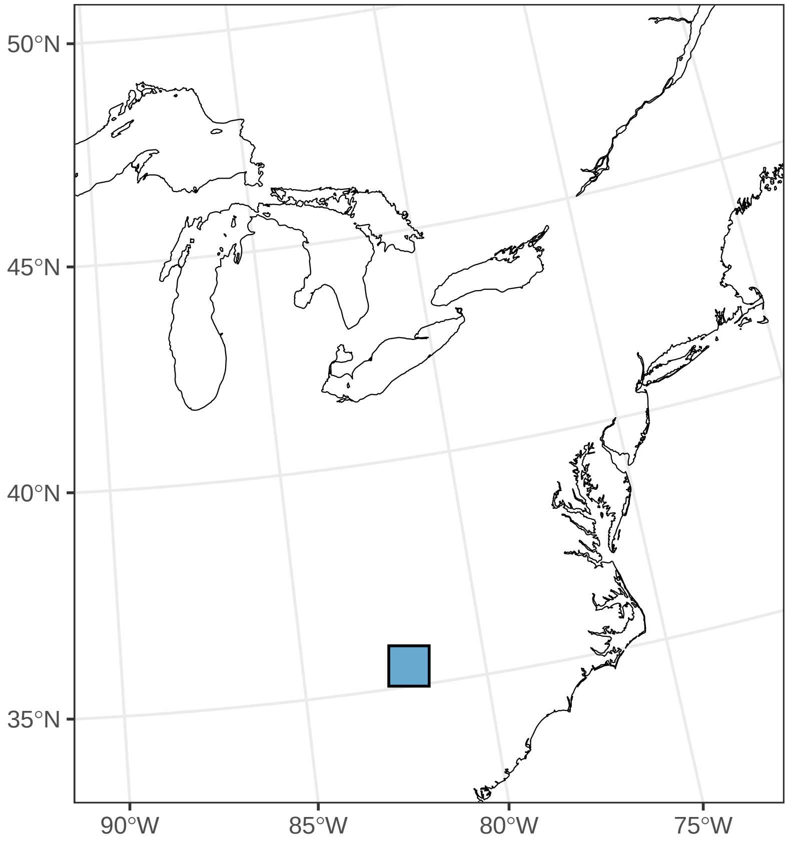

#### *Podophyllum peltatum*

Ephemerality index: 0.055

n = 91

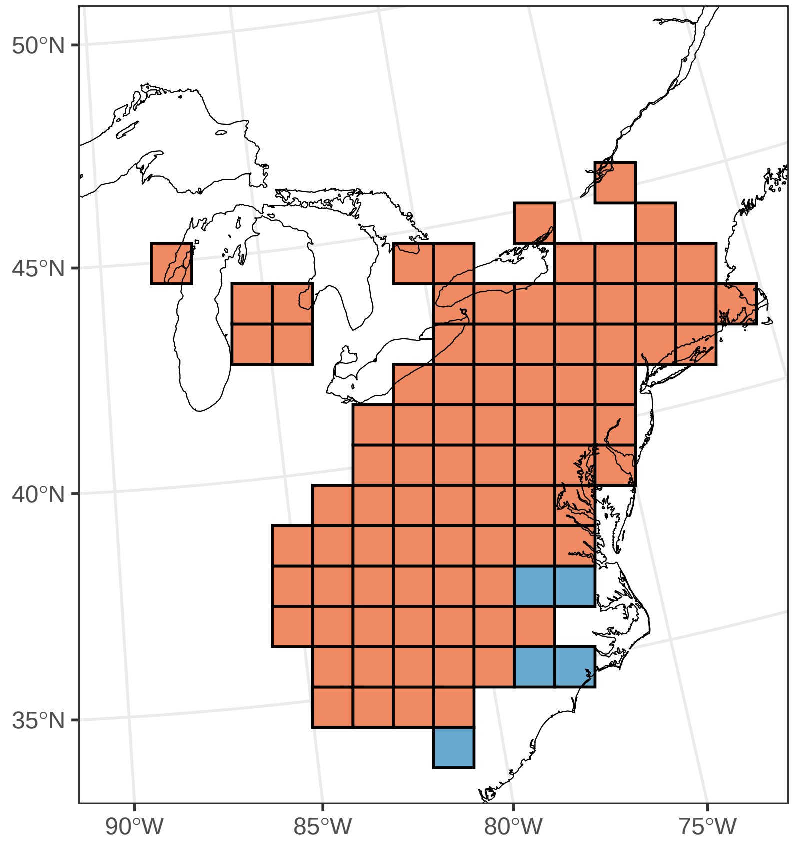

#### *Polygonatum pubescens*

Ephemerality index: 0.045

n = 44

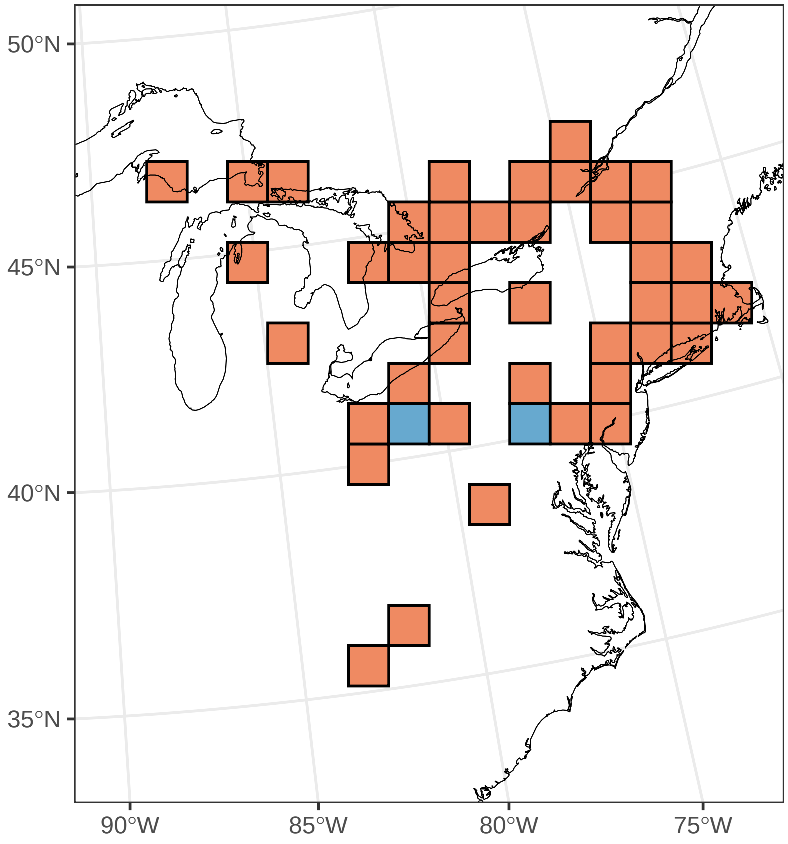

### ***Potentilla simplex***

Ephemerality index: 0.032

n = 31

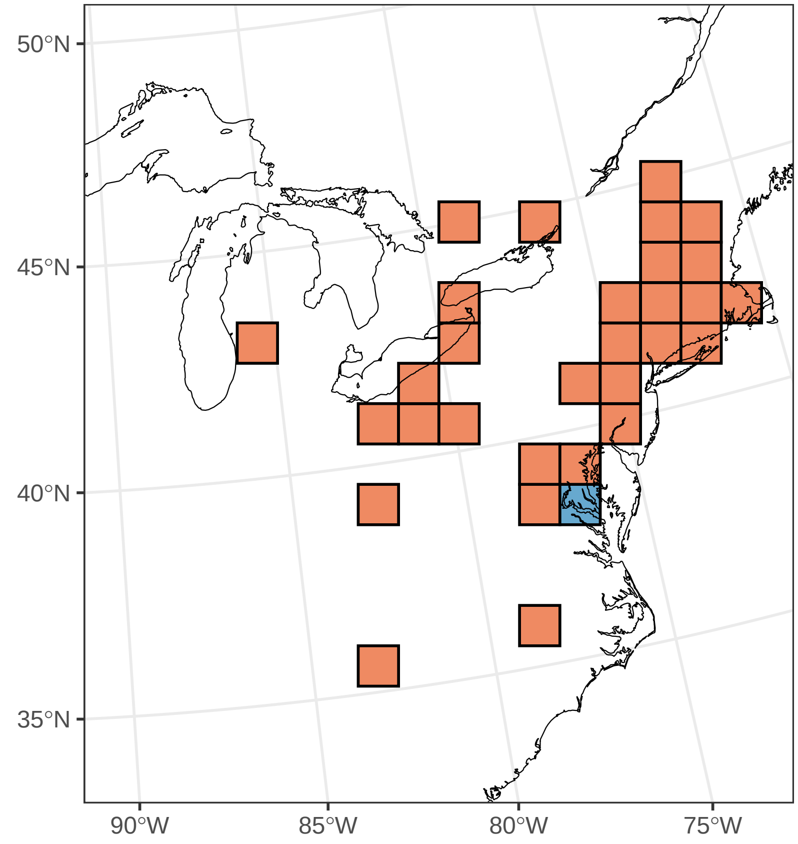

#### ***Primula mistassinica***

Ephemerality index: 1

n = 1

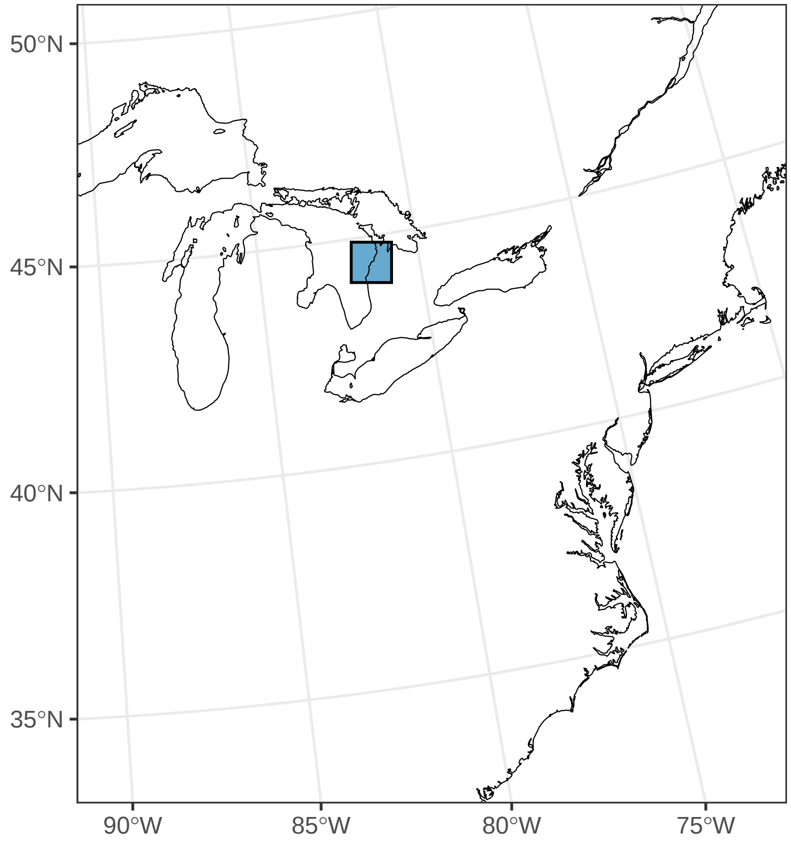

#### ***Prosartes maculata***

Ephemerality index: 1

n = 1

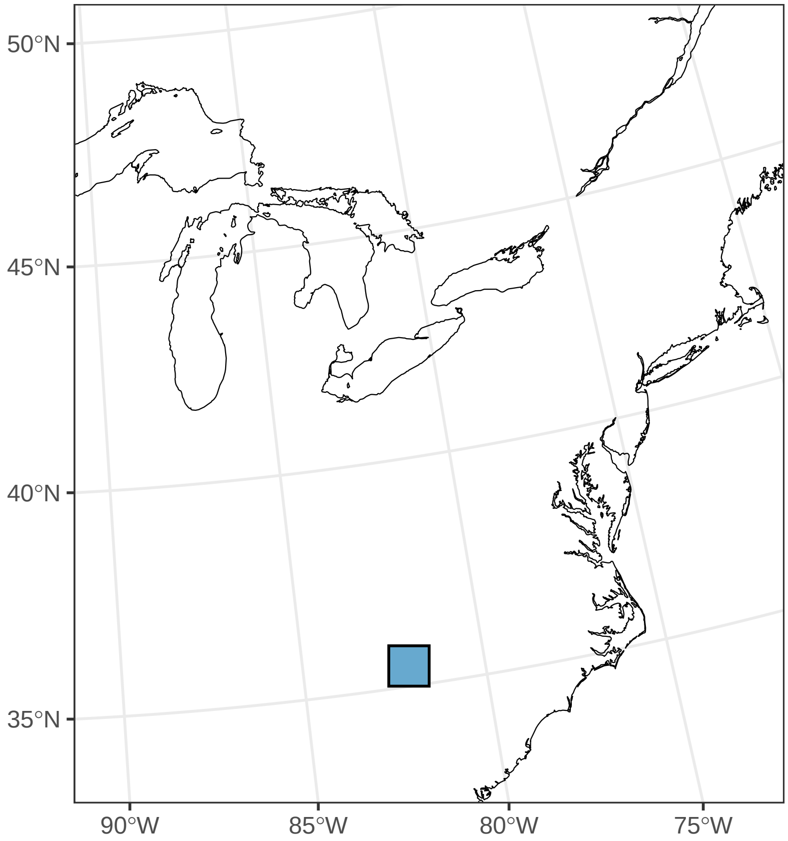

#### ***Ranunculus abortivus***

Ephemerality index: 0.548

n = 42

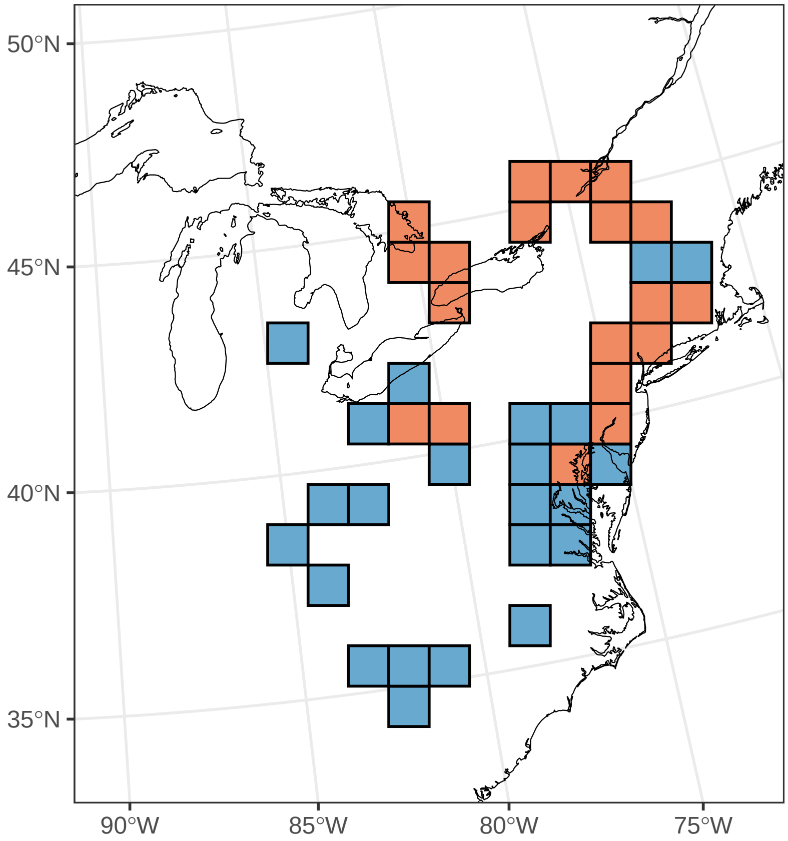

#### ***Ranunculus hispidus***

Ephemerality index: 0.6

n = 5

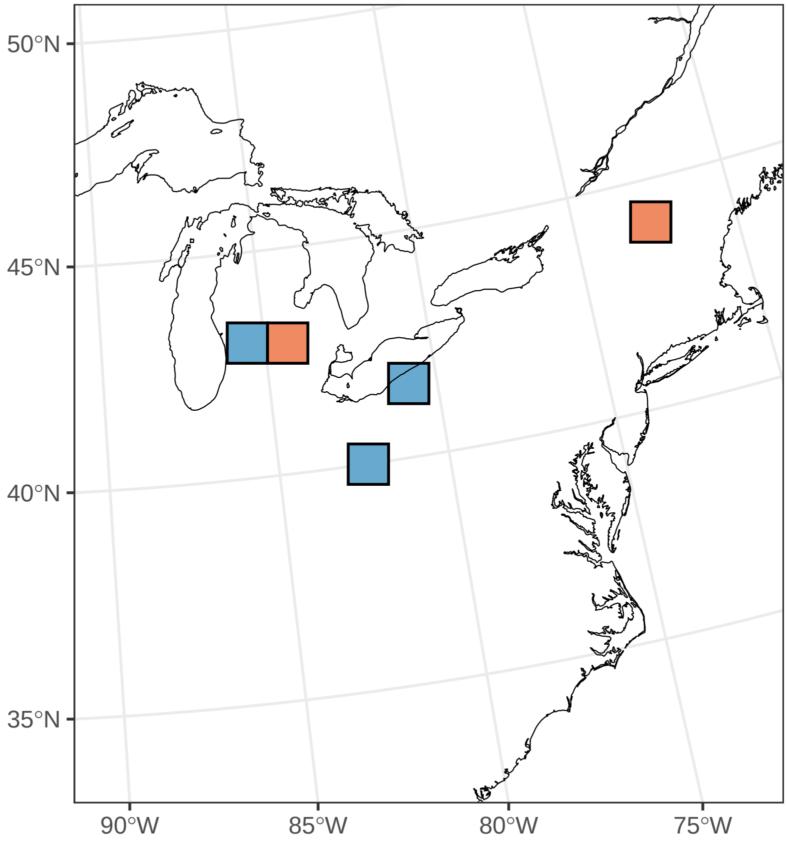

#### ***Ranunculus parviflorus***

Ephemerality index: 1

n = 1

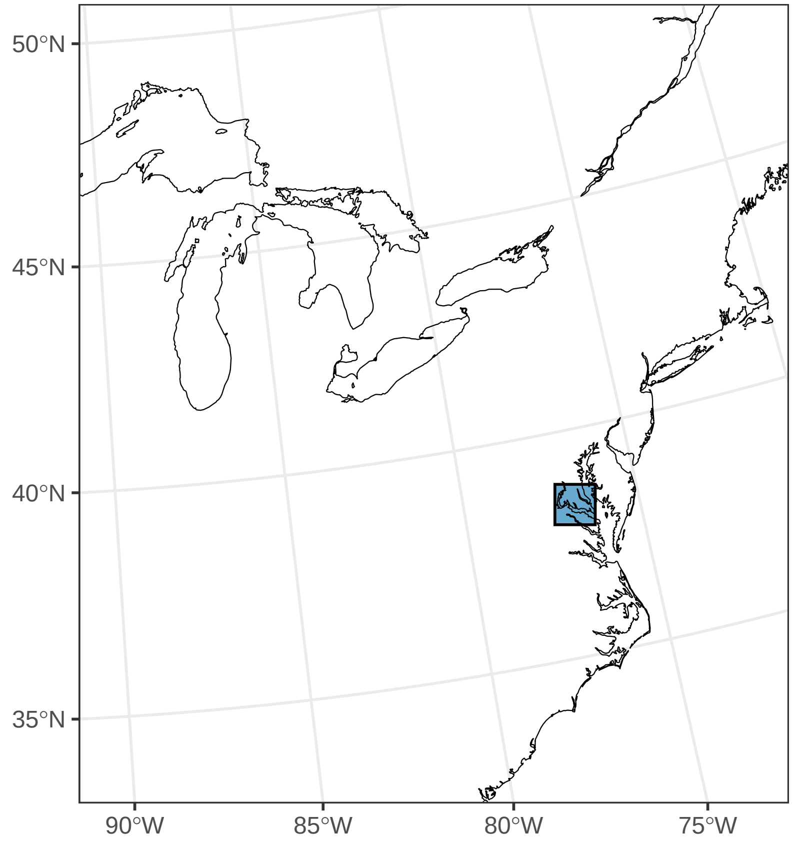

#### ***Ranunculus recurvatus***

Ephemerality index: 0.194

n = 31

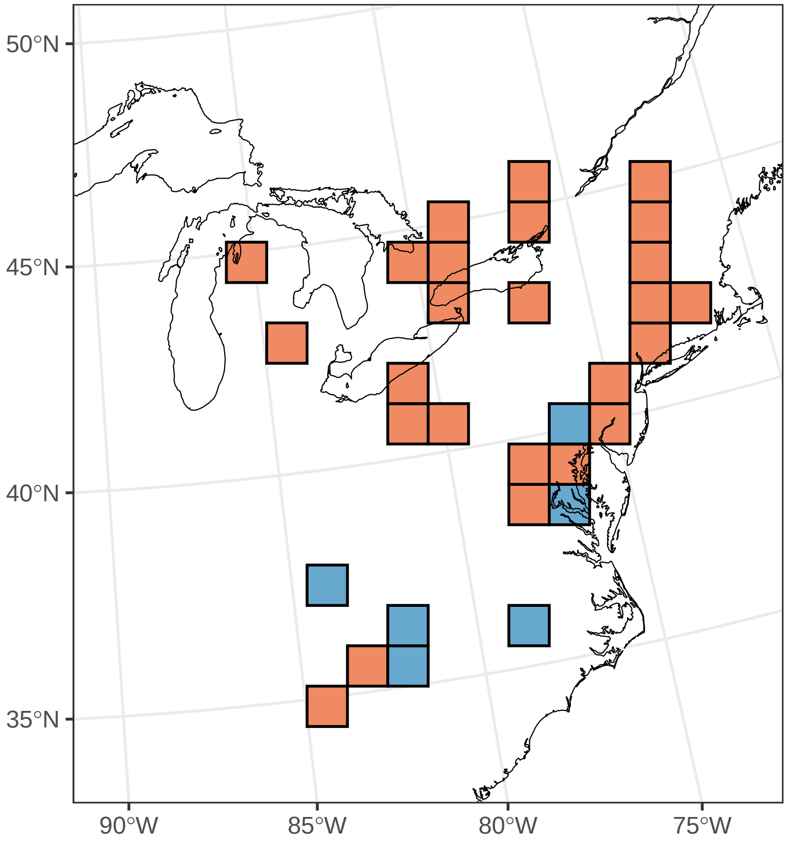

#### ***Salvia urticifolia***

Ephemerality index: 1

n = 1

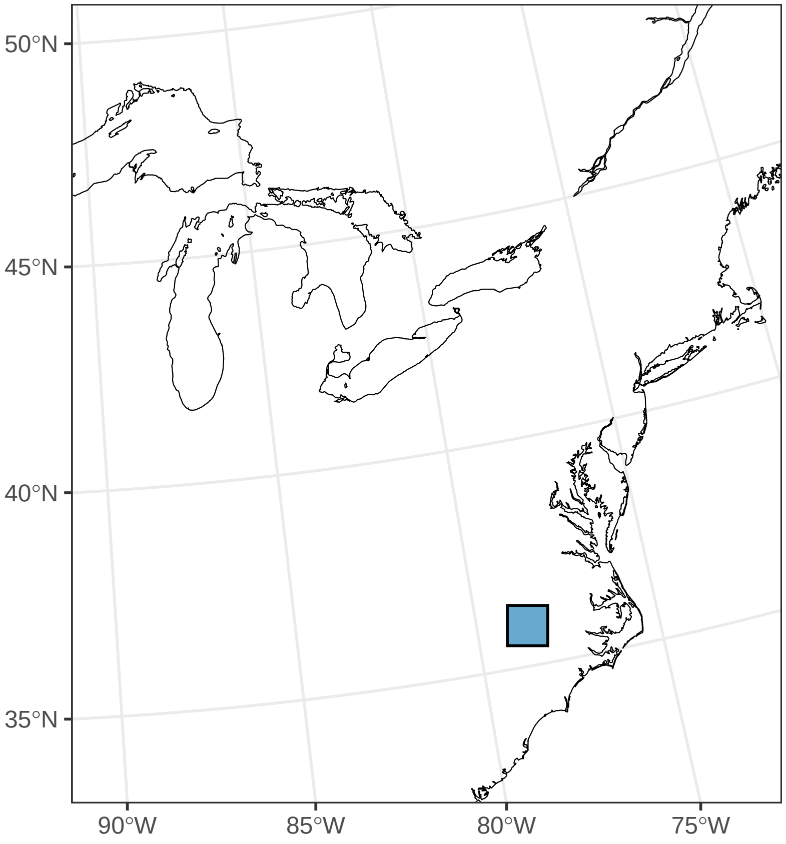

##### *Sanguinaria canadensis*

Ephemerality index: 0.059

n = 101

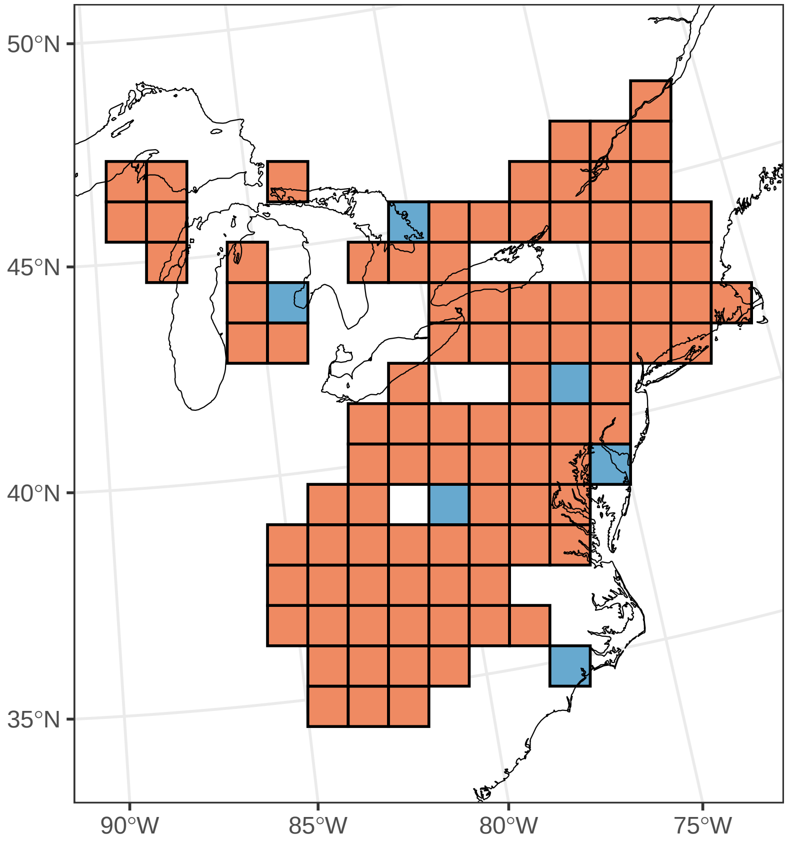

#### ***Scilla siberica***

Ephemerality index: 1

n = 20

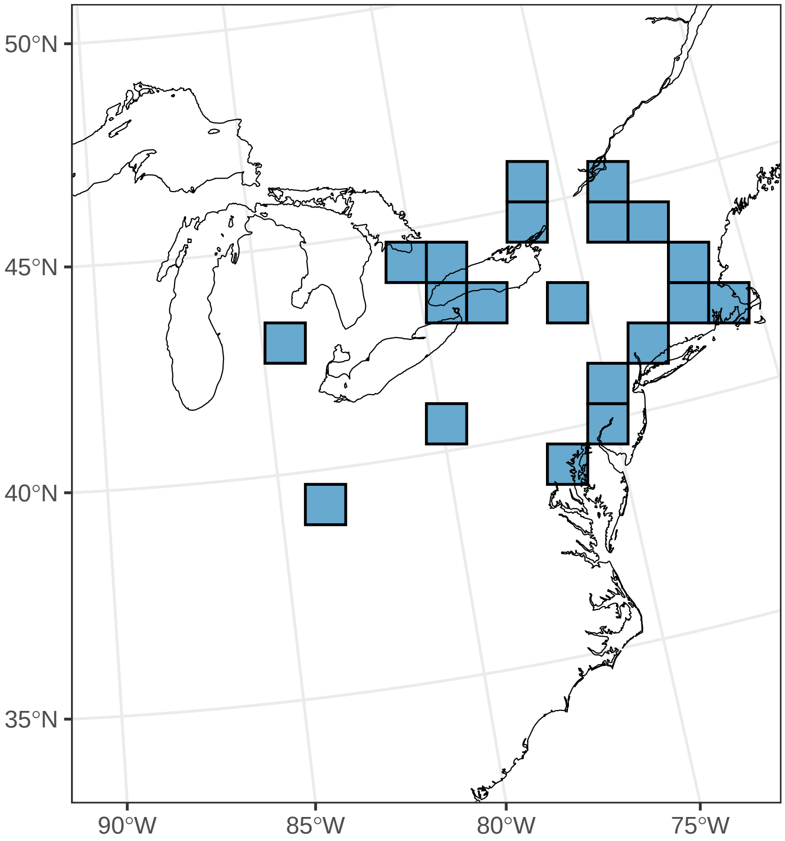

#### ***Sisyrinchium rosulatum***

Ephemerality index: 0.143

n = 7

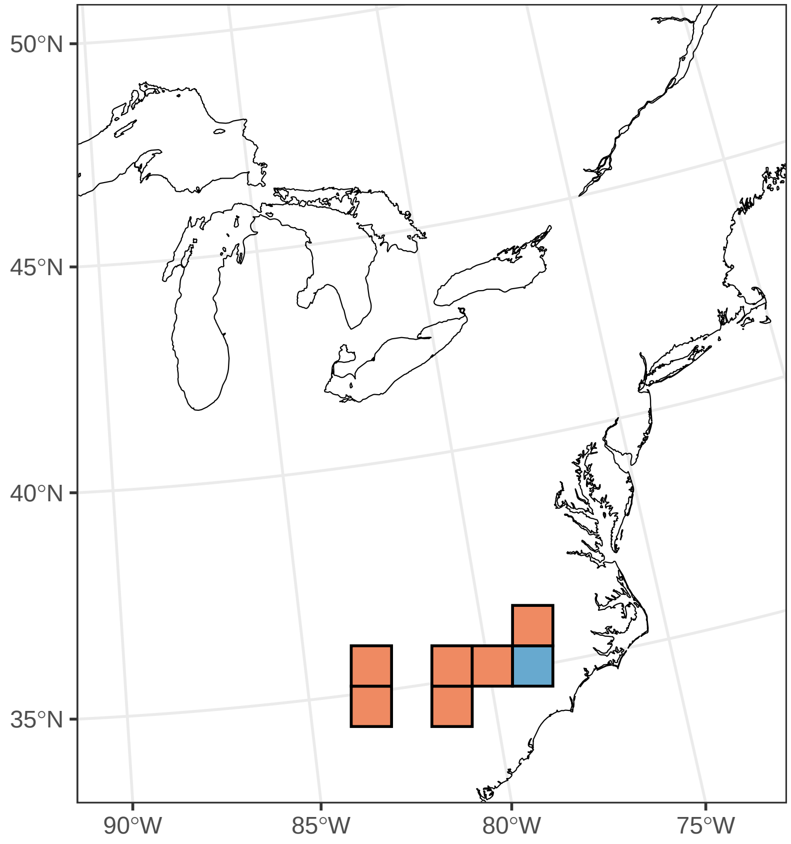

#### ***Stellaria media***

Ephemerality index: 0.031

n = 32

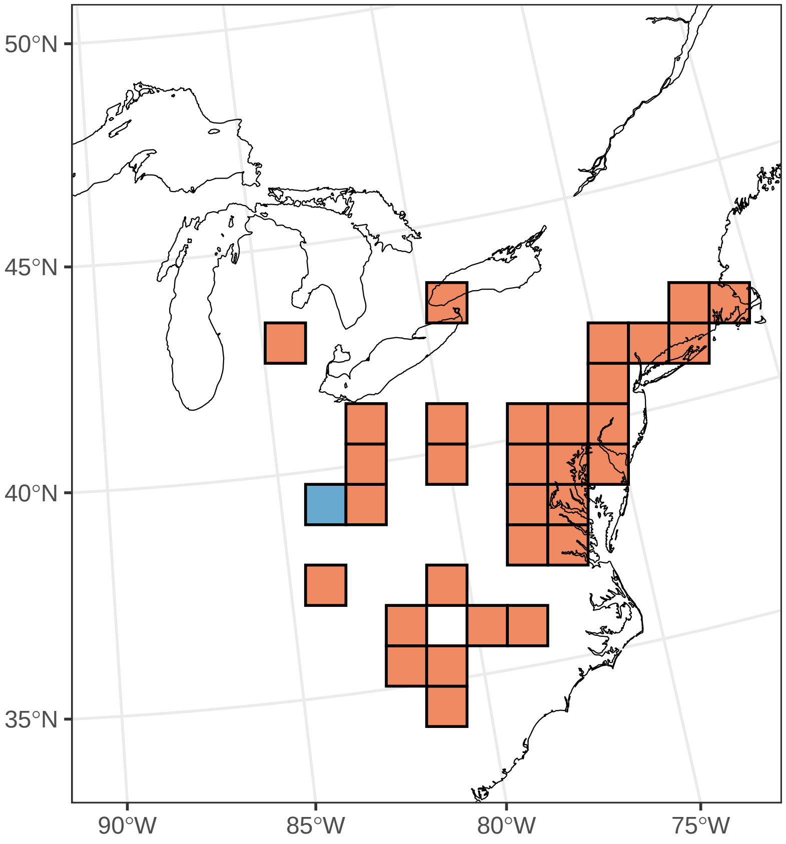

#### ***Stellaria pubera***

Ephemerality index: 0.417

n = 36

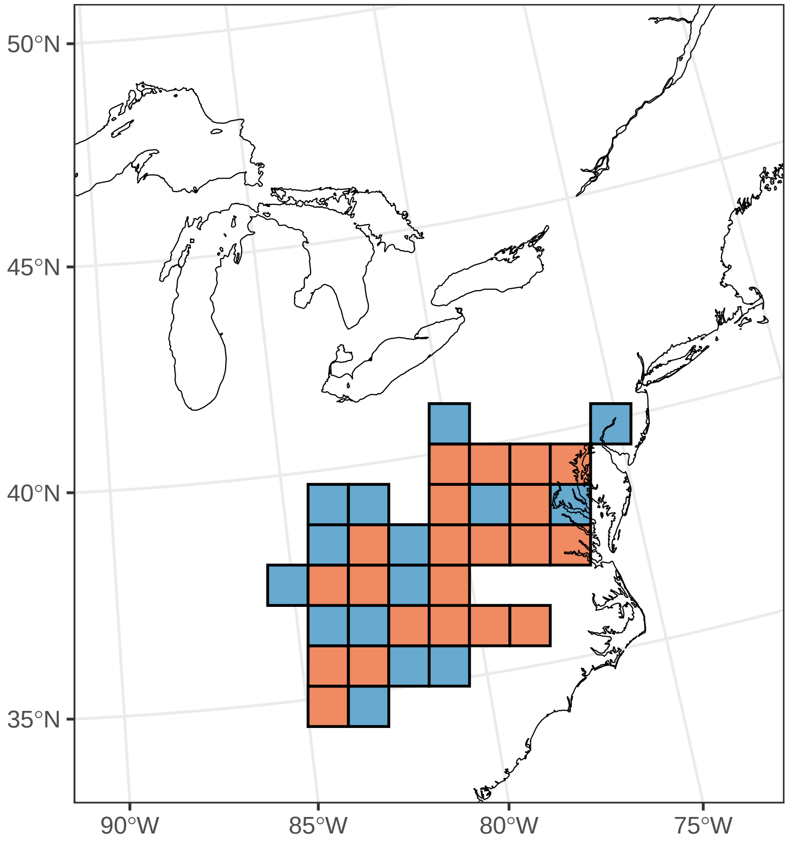

#### *Stylophorum diphyllum*

Ephemerality index: 0.133

n = 30

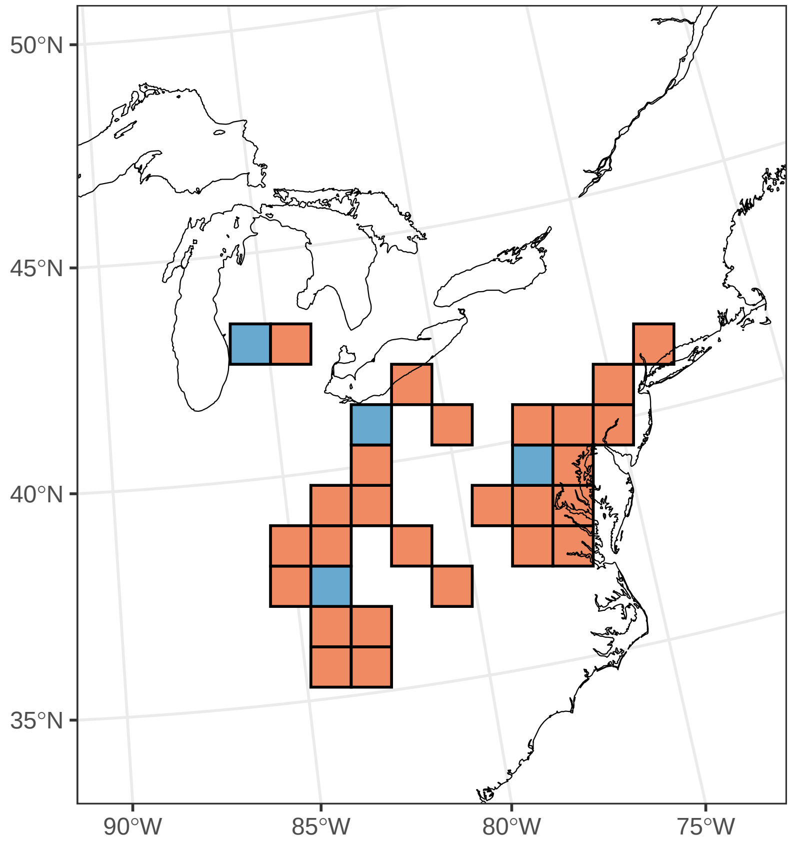

***Symplocarpus foetidus***

Ephemerality index: 0.06

n = 67

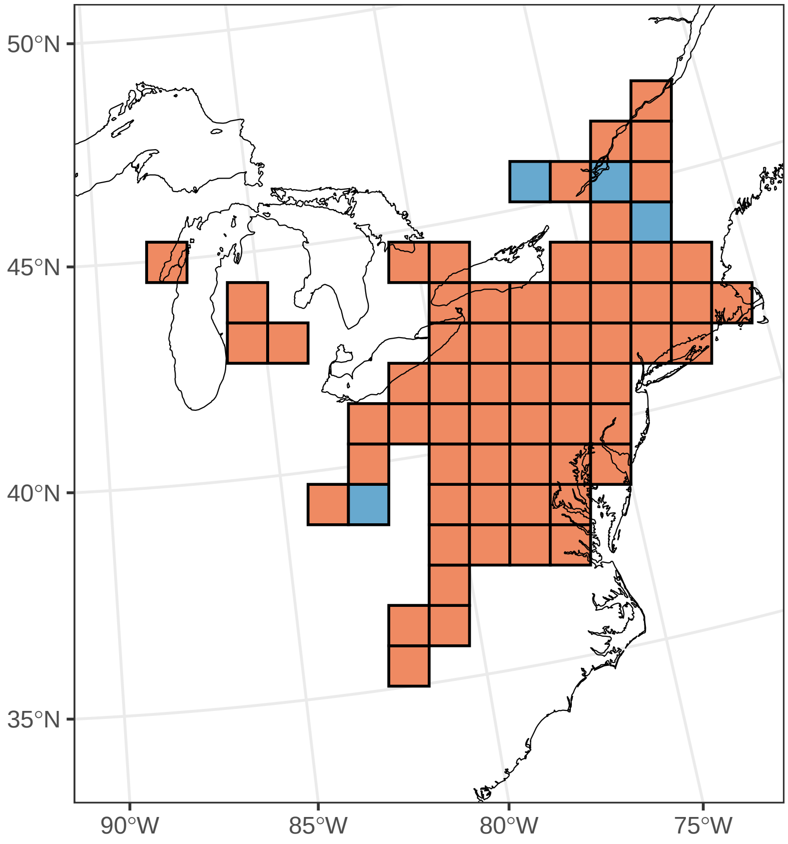

#### ***Thalictrum dioicum***

Ephemerality index: 0.111

n = 27

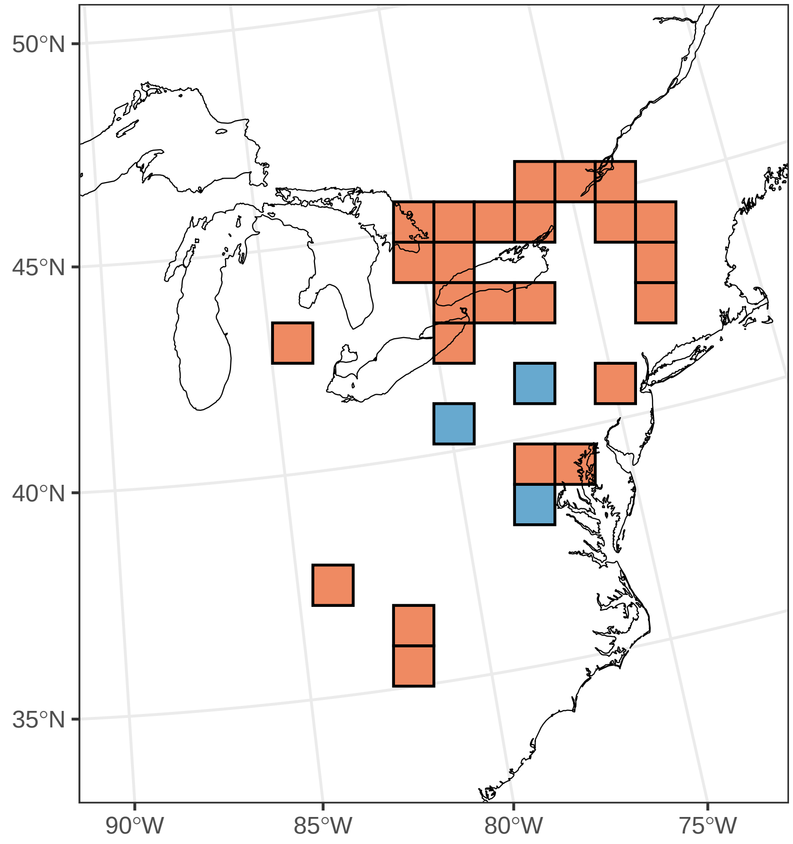

#### ***Thaspium barbinode***

Ephemerality index: 1

n = 1

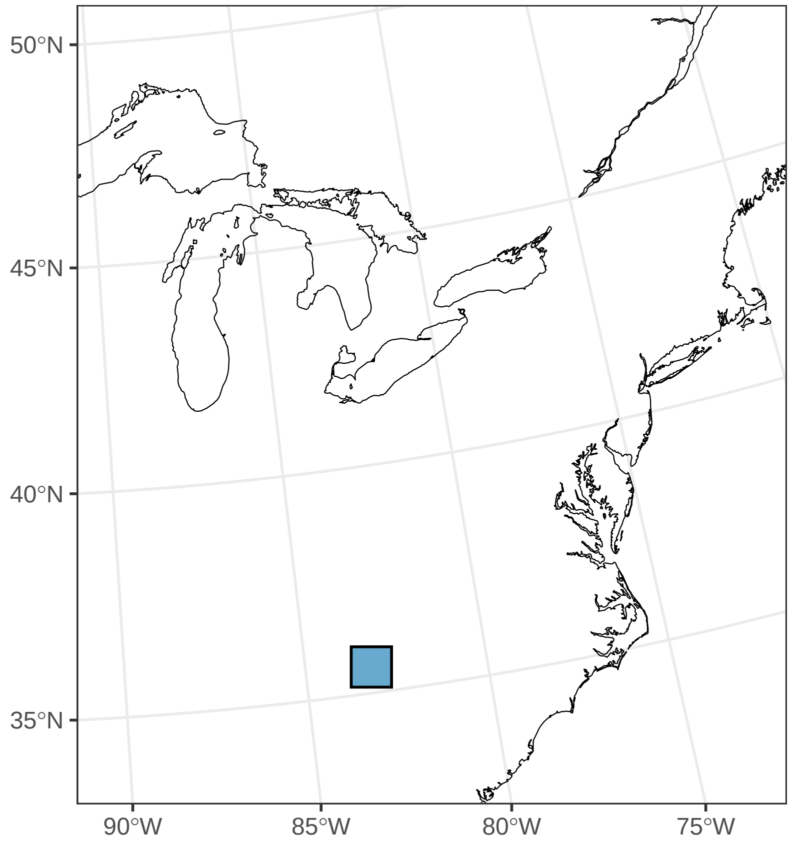

#### ***Trillium catesbaei***

Ephemerality index: 0.111

n = 9

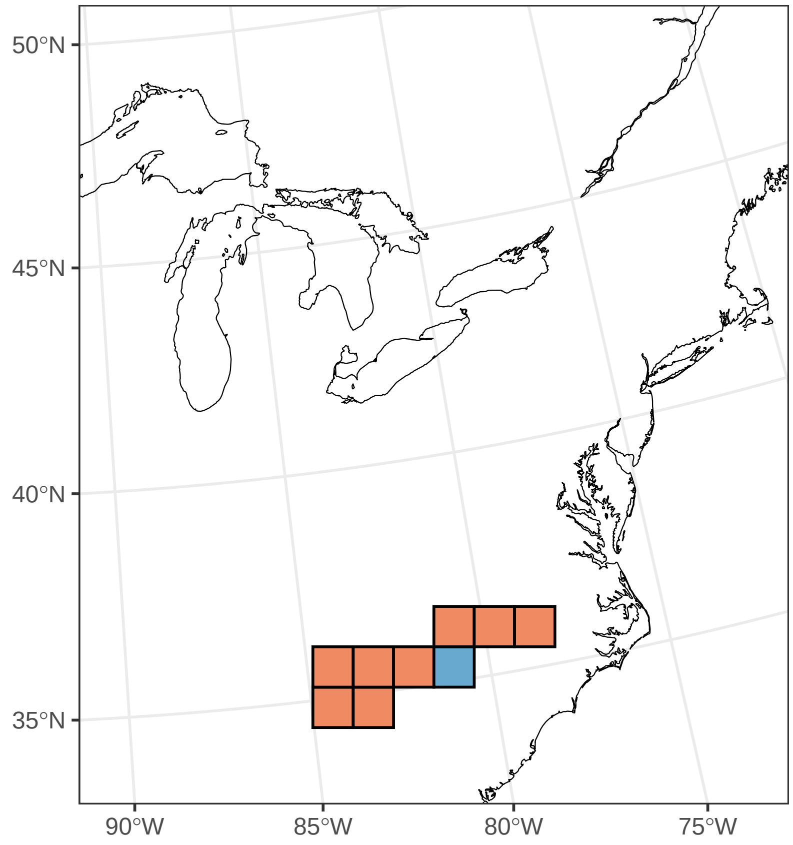

#### ***Trillium cernuum***

Ephemerality index: 0.154

n = 13

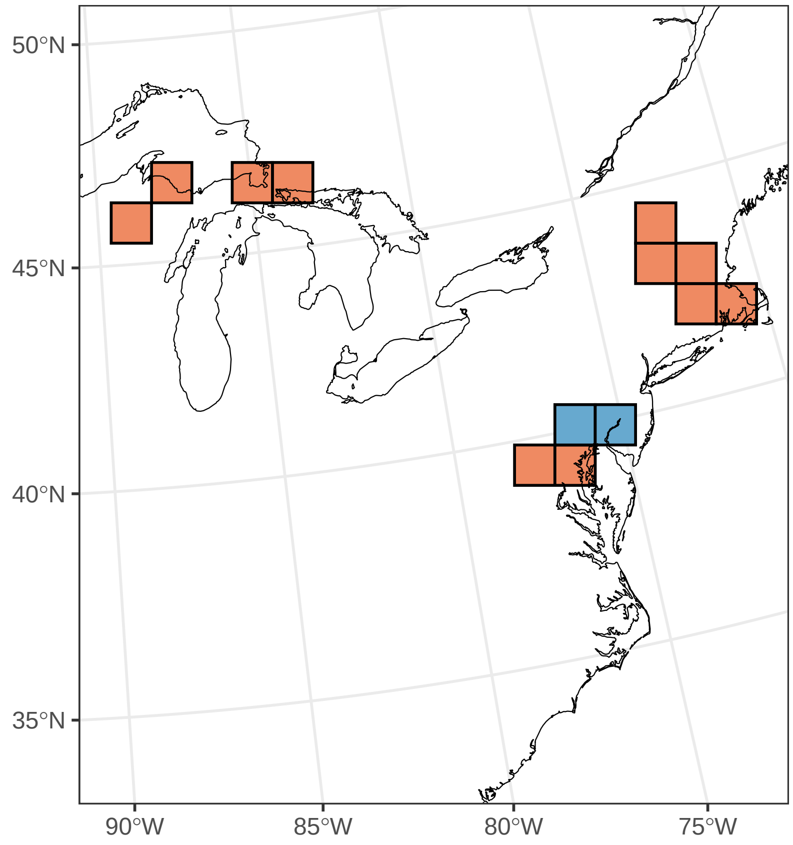

#### ***Trillium cuneatum***

Ephemerality index: 0.556

n = 18

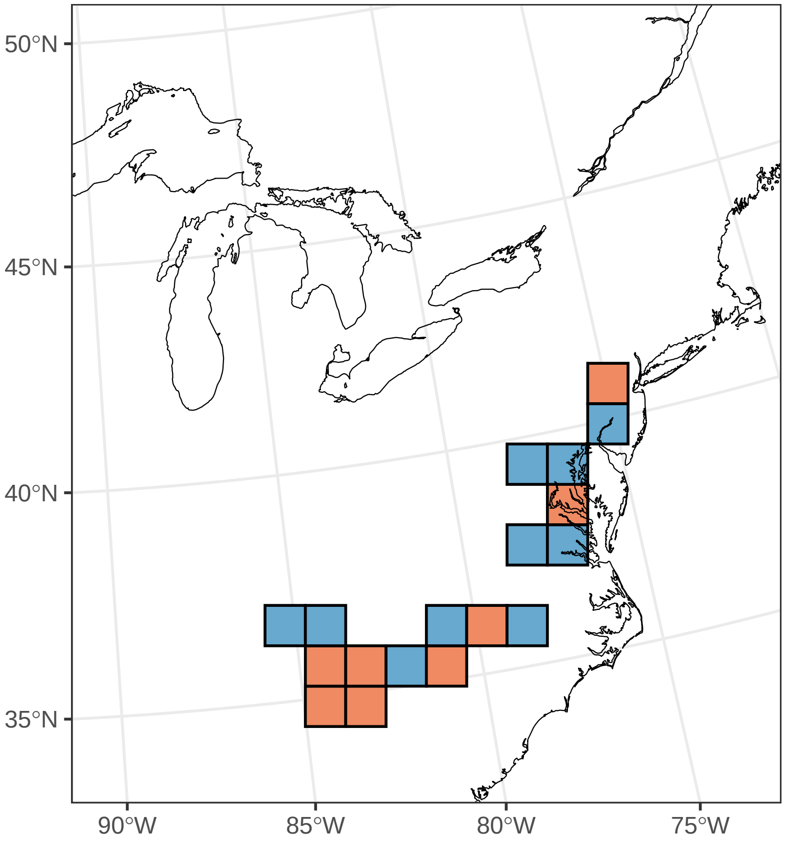

#### ***Trillium erectum***

Ephemerality index: 0.089

n = 79

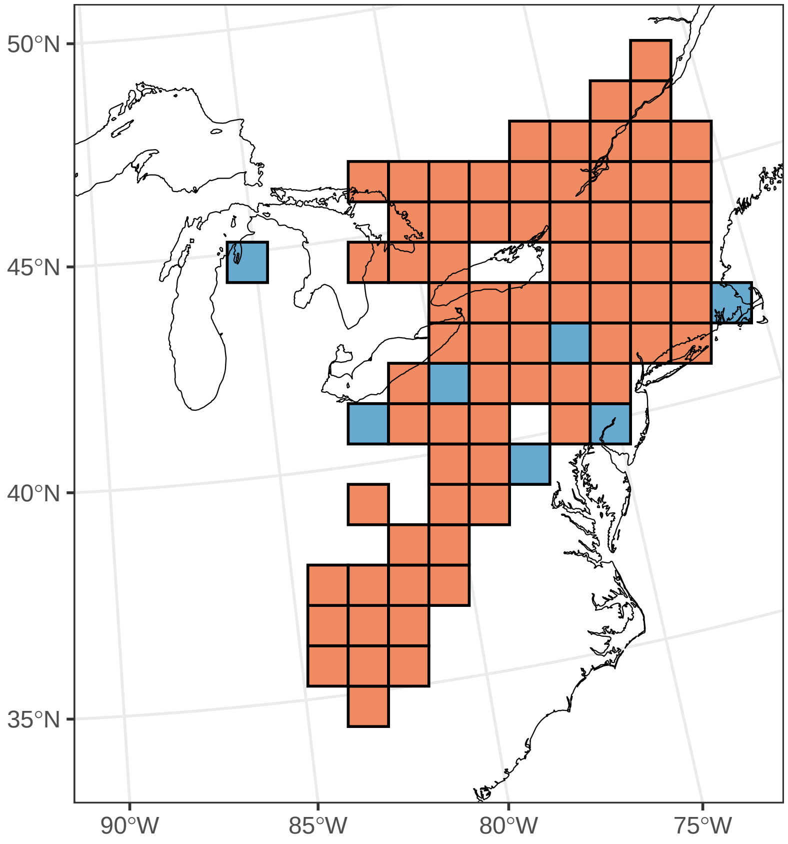

#### ***Trillium flexipes***

Ephemerality index: 0.875

n = 8

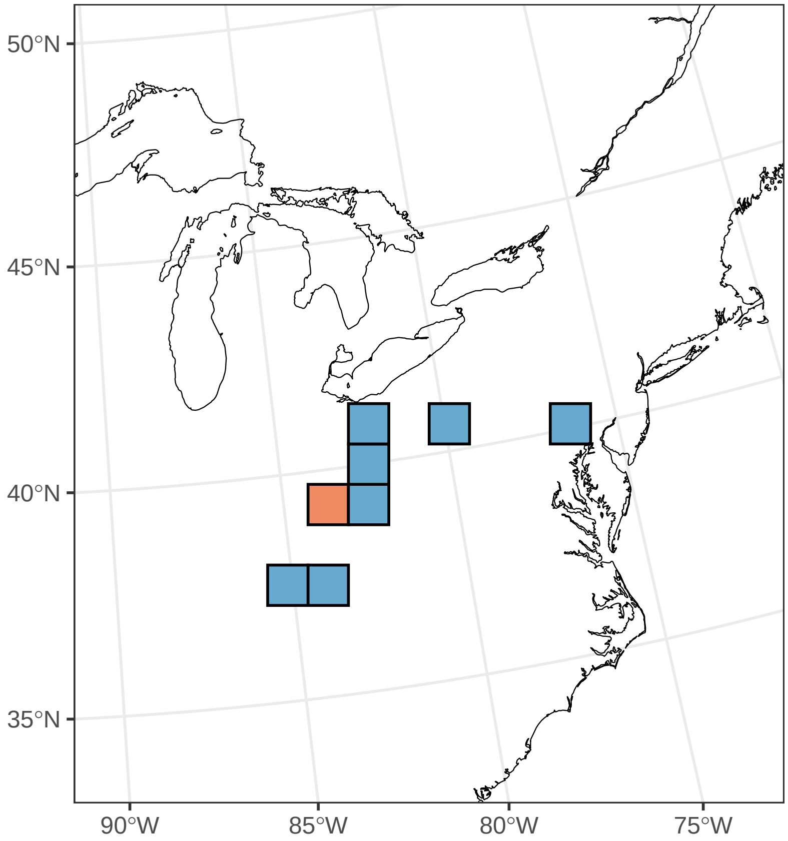

#### ***Trillium grandiflorum***

Ephemerality index: 0.35

n = 80

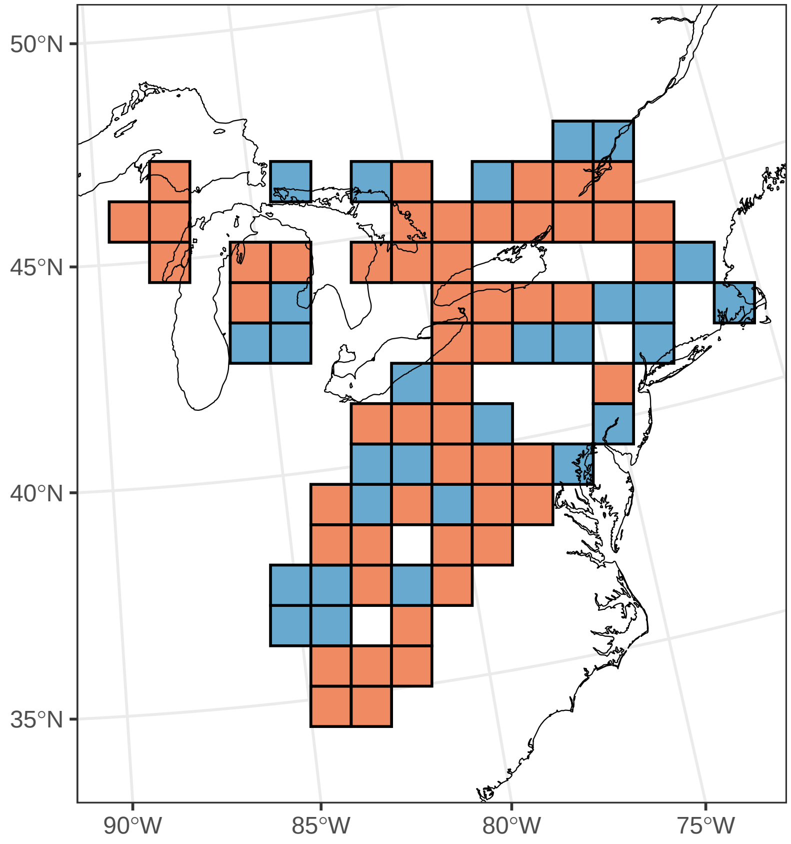

#### ***Trillium luteum***

Ephemerality index: 0.571

n = 7

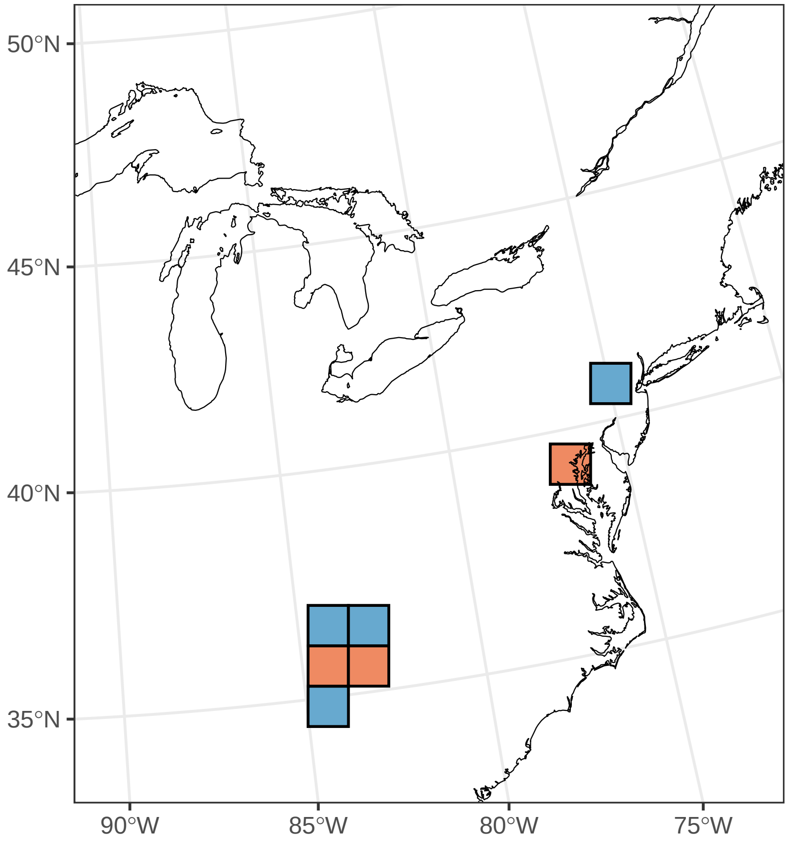

#### ***Trillium sessile***

Ephemerality index: 0.867

n = 15

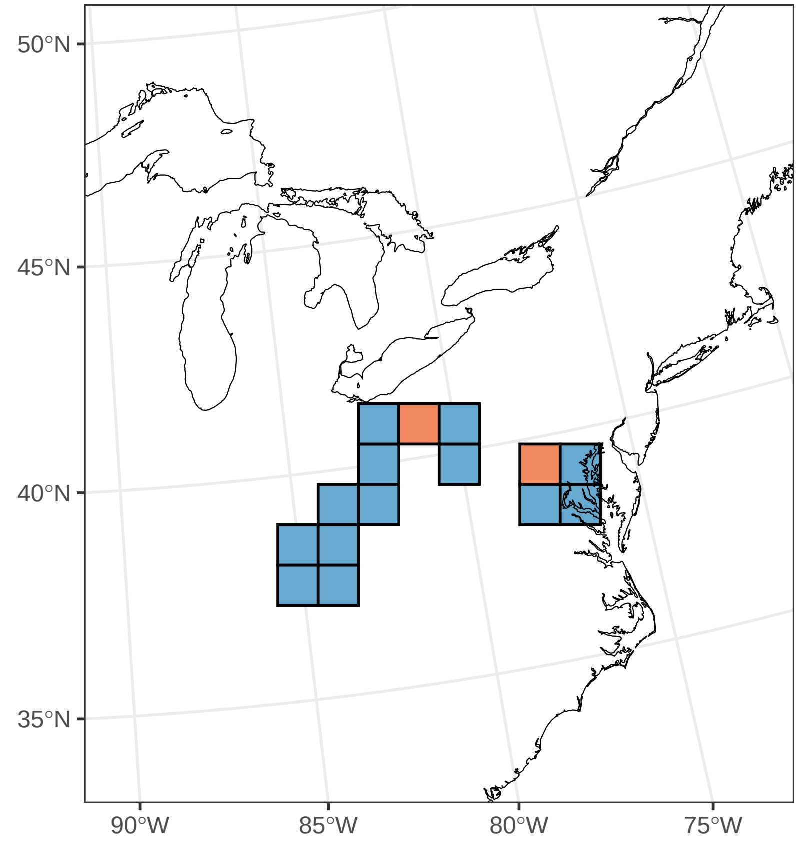

#### ***Trillium simile***

Ephemerality index: 1

n = 2

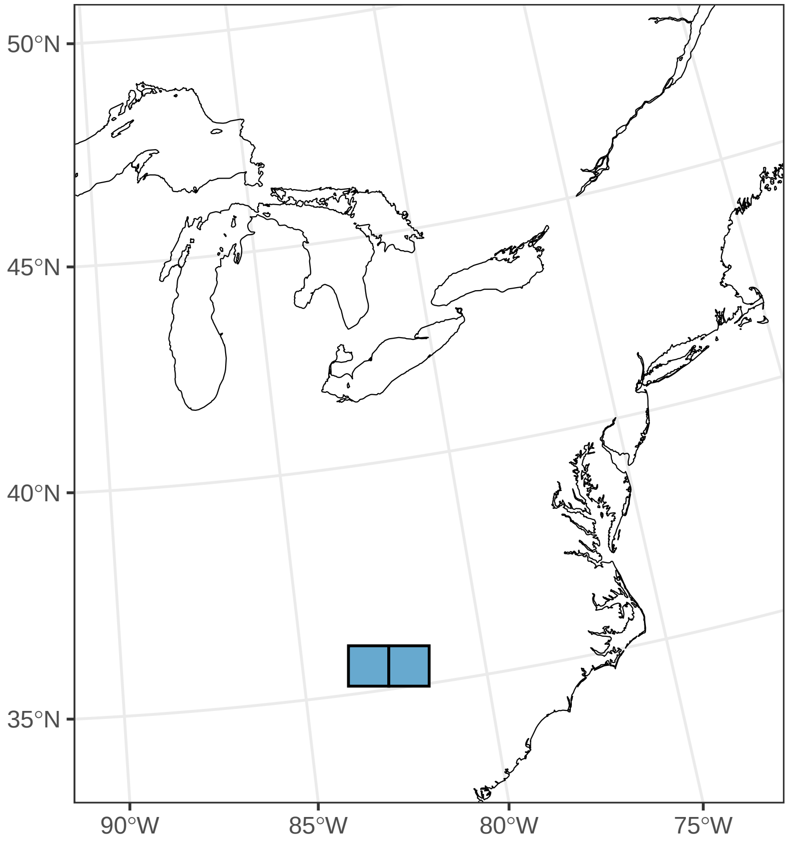

#### ***Trillium vaseyi***

Ephemerality index: 0.2

n = 5

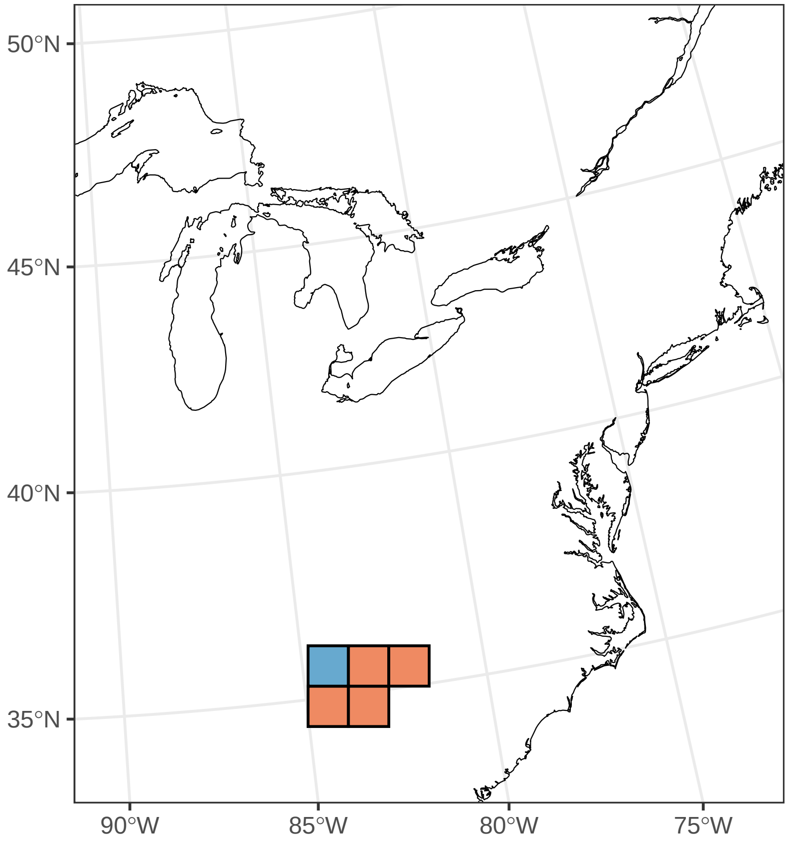

#### ***Uvularia grandiflora***

Ephemerality index: 0.162

n = 37

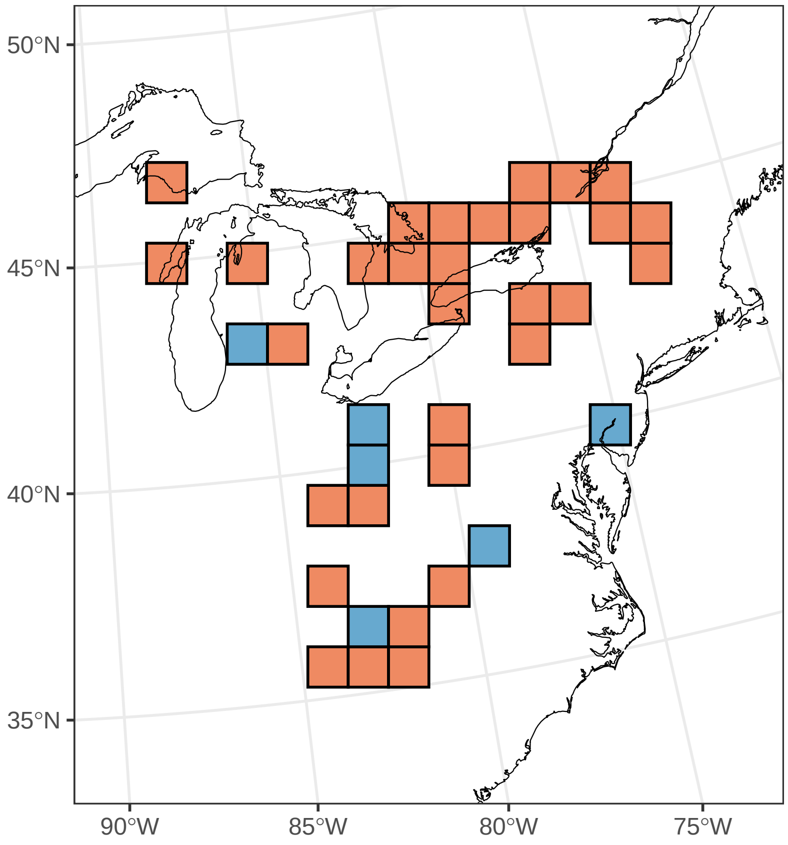

#### ***Uvularia perfoliata***

Ephemerality index: 0.103

n = 29

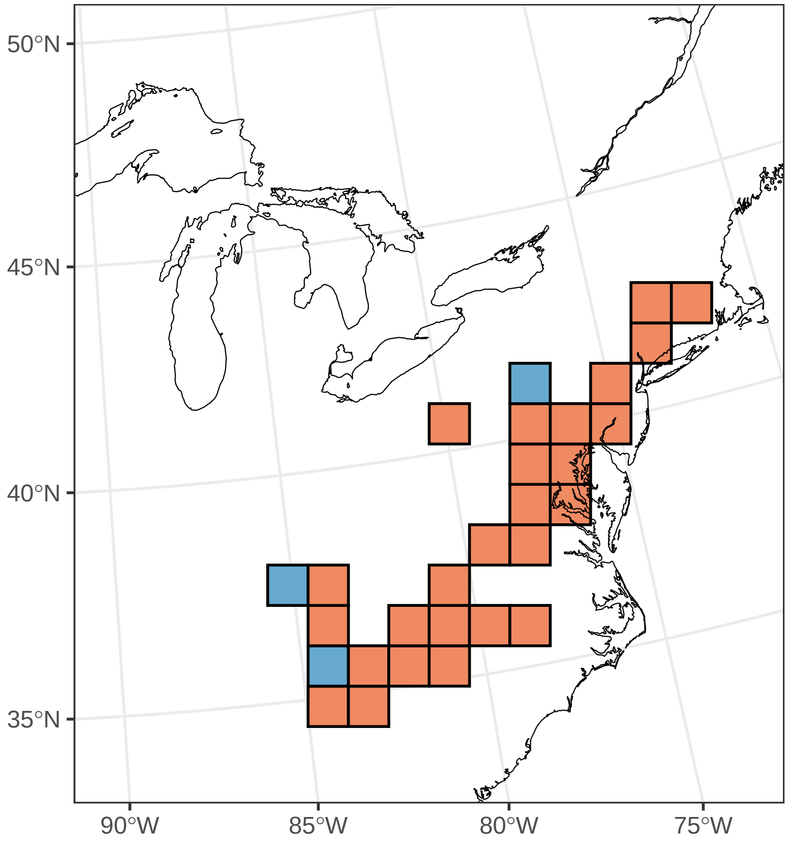

#### ***Uvularia sessilifolia***

Ephemerality index: 0.079

n = 38

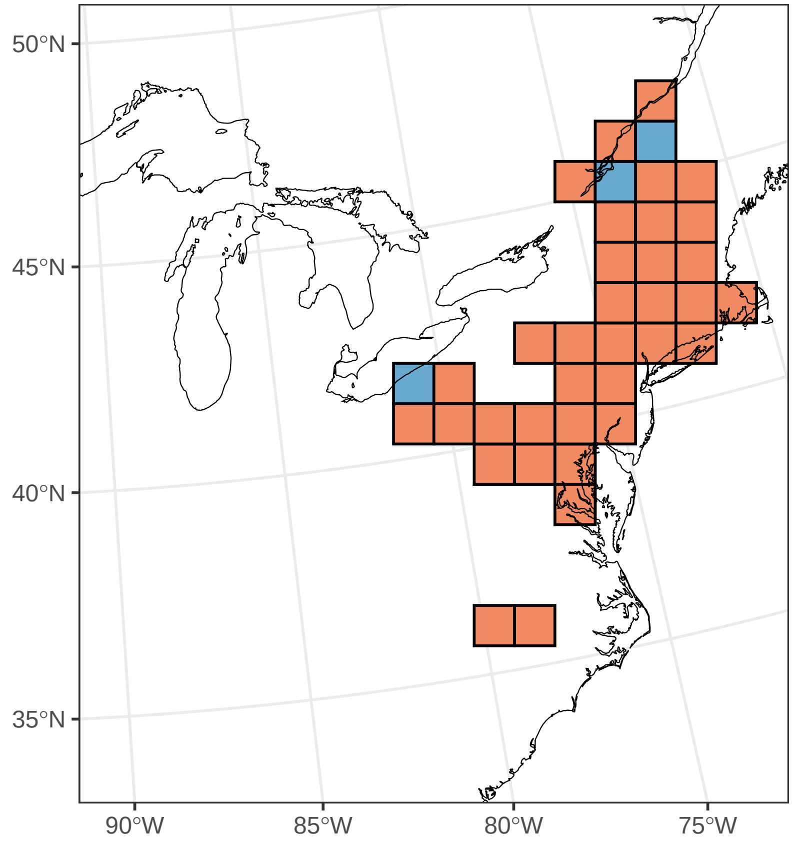

#### ***Valerianella locusta***

Ephemerality index: 0.9

n = 10

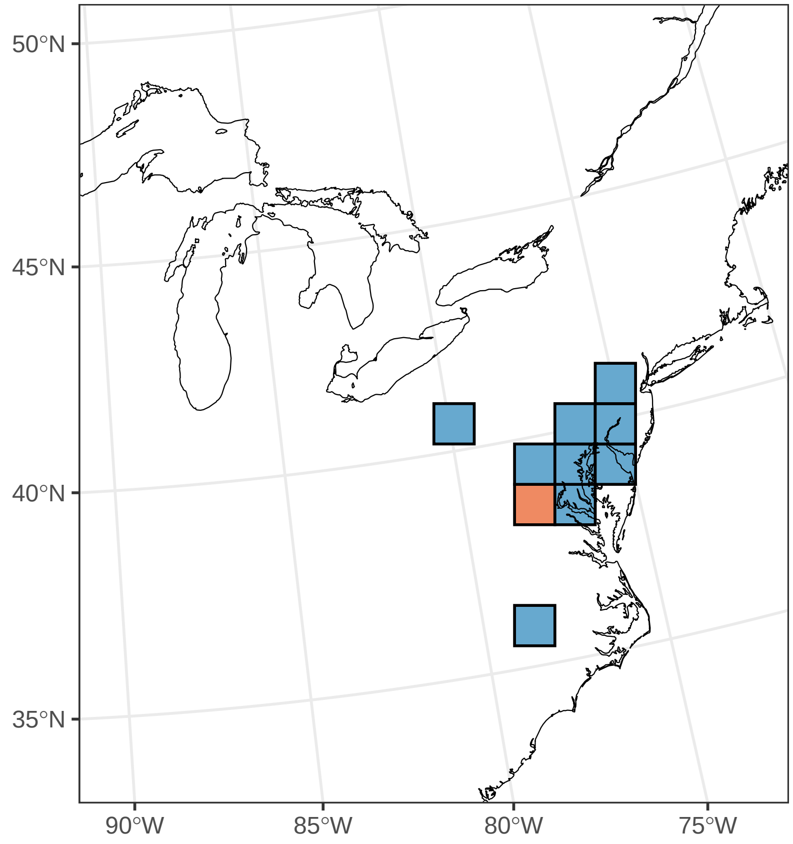

#### ***Valerianella radiata***

Ephemerality index: 1

n = 1

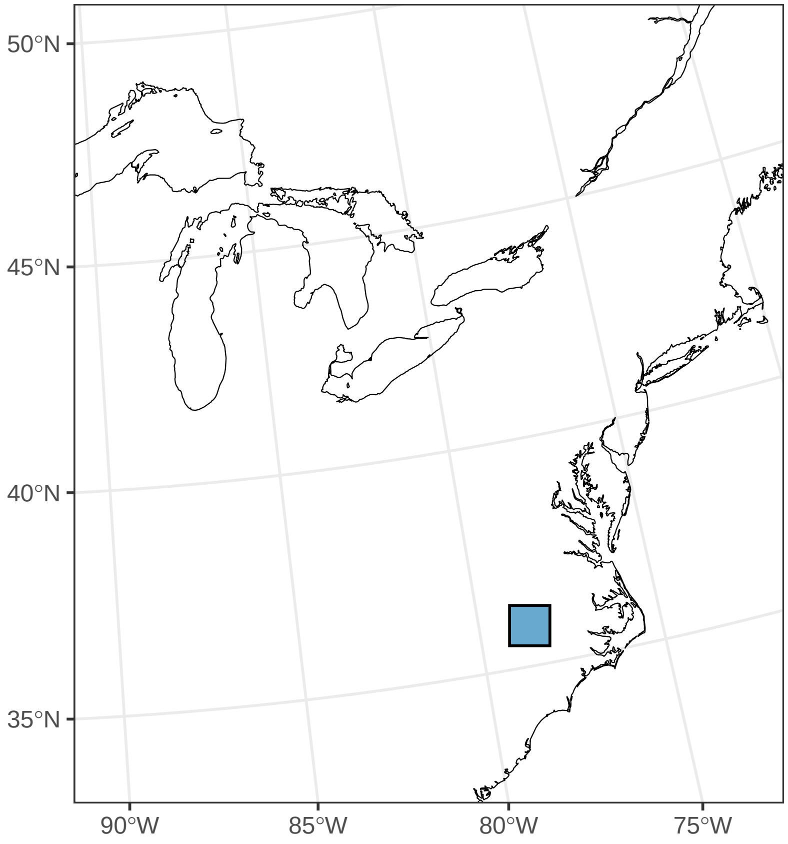

#### ***Viola bicolor***

Ephemerality index: 1

n = 18

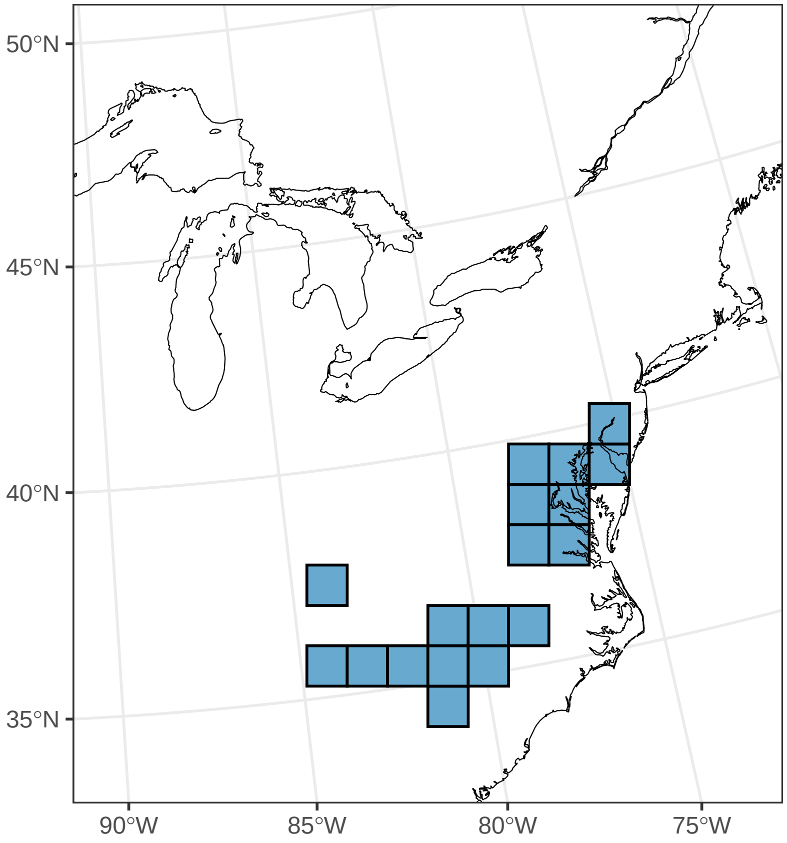

#### ***Viola blanda***

Ephemerality index: 0.143

n = 7

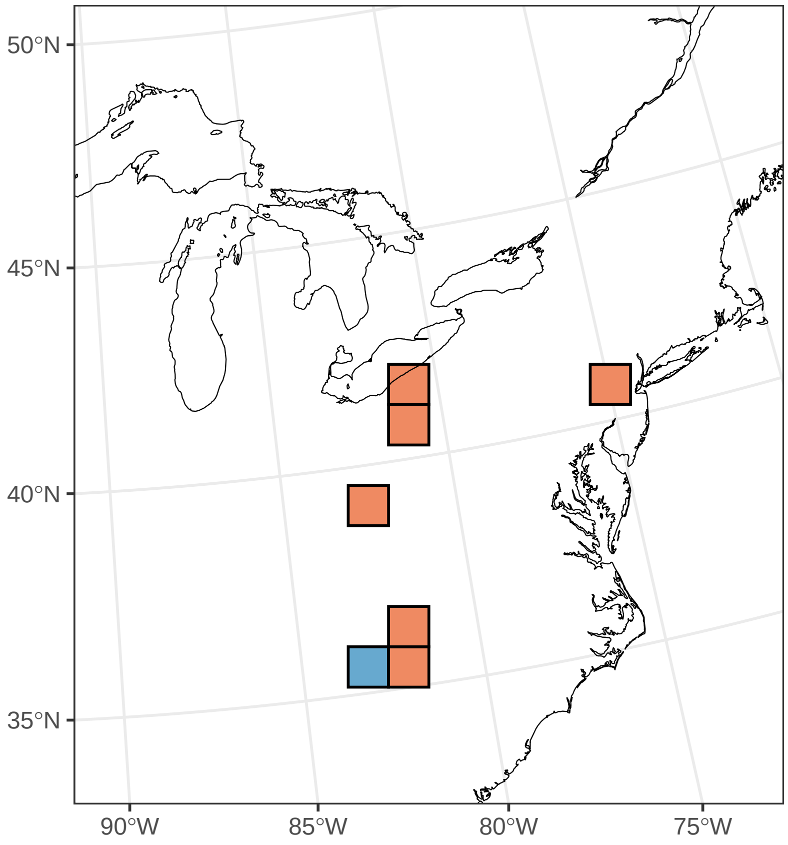

#### ***Viola canadensis***

Ephemerality index: 0.118

n = 34

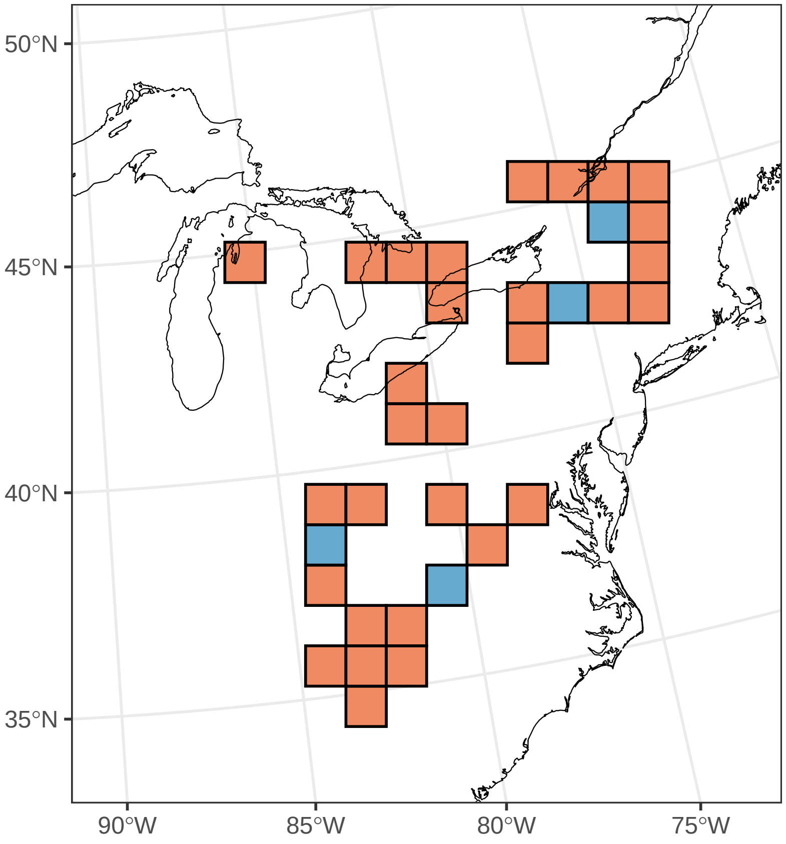

#### ***Viola hastata***

Ephemerality index: 0.368

n = 19

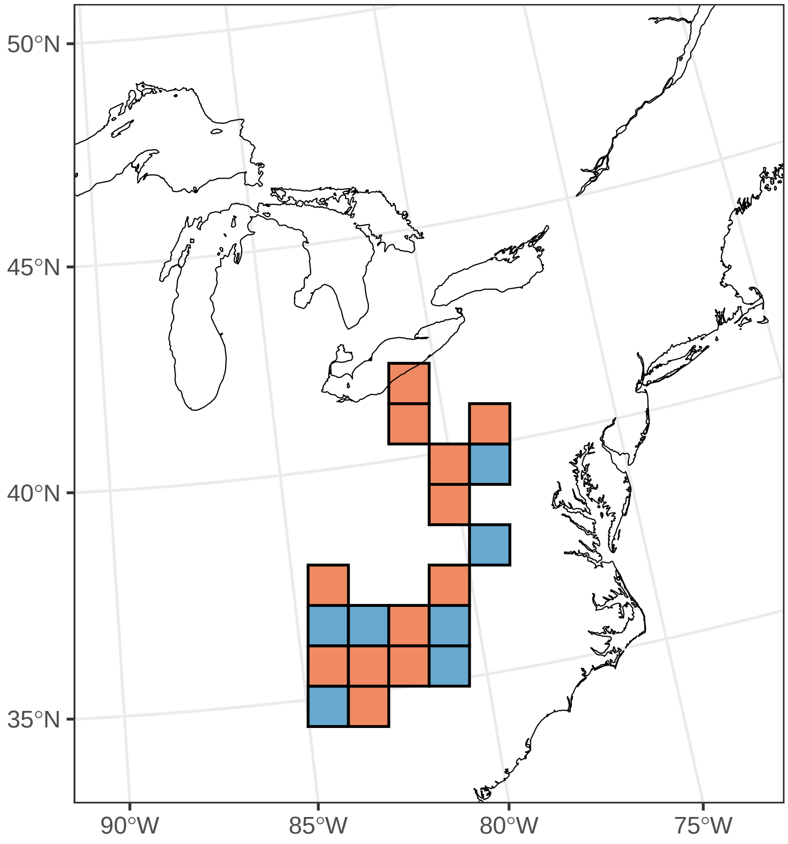

#### *Viola lanceolata*

Ephemerality index: 0.167

n = 6

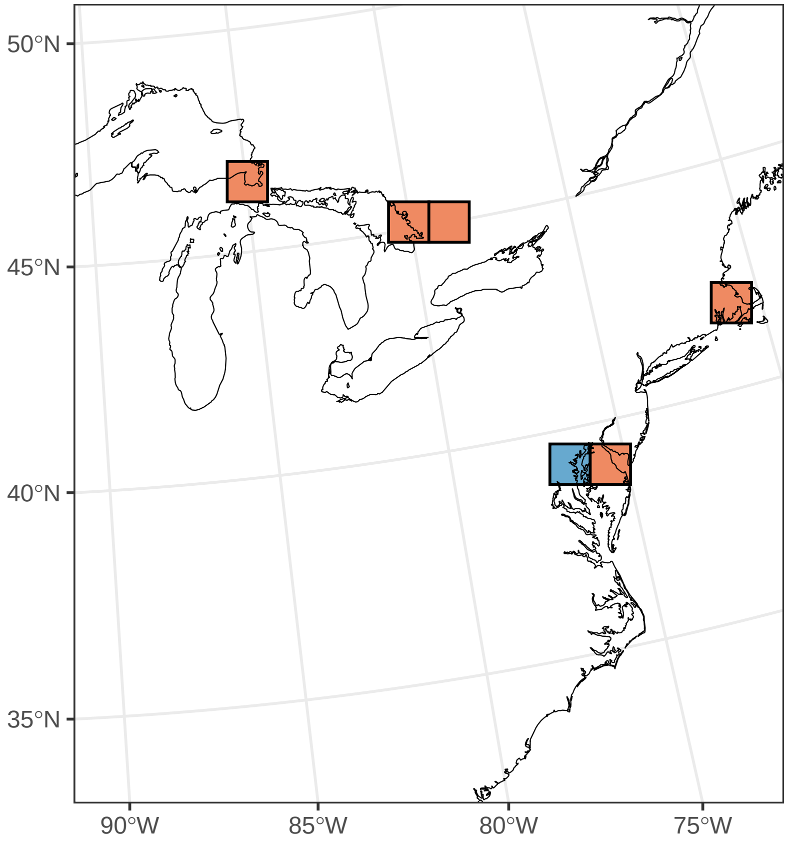

#### ***Viola palmata***

Ephemerality index: 0.167

n = 6

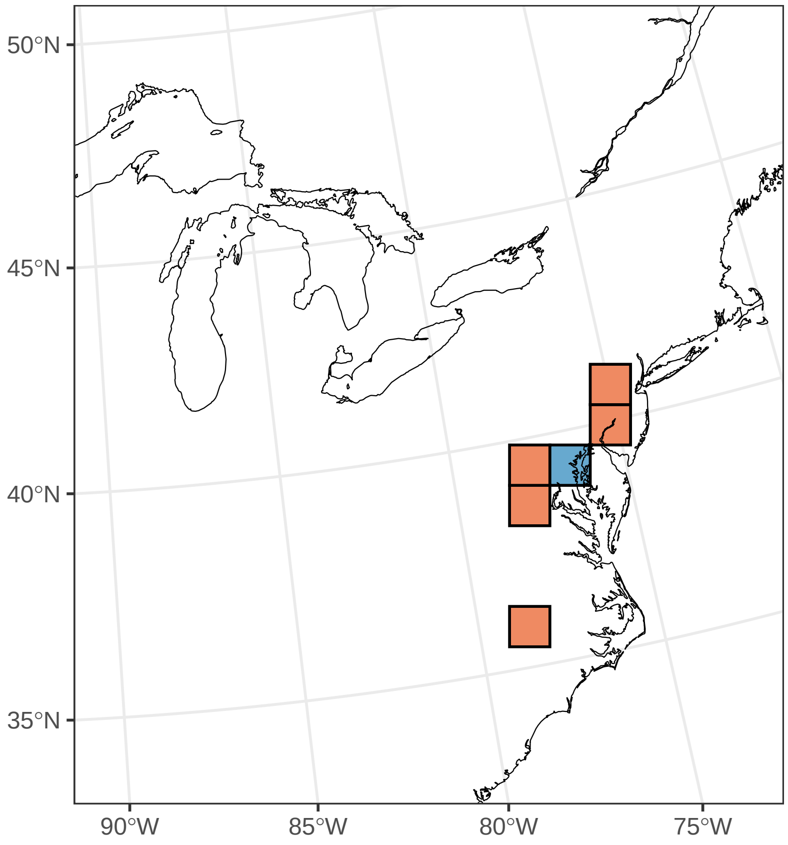

##### ***Viola pubescens***

Ephemerality index: 0.386

n = 57

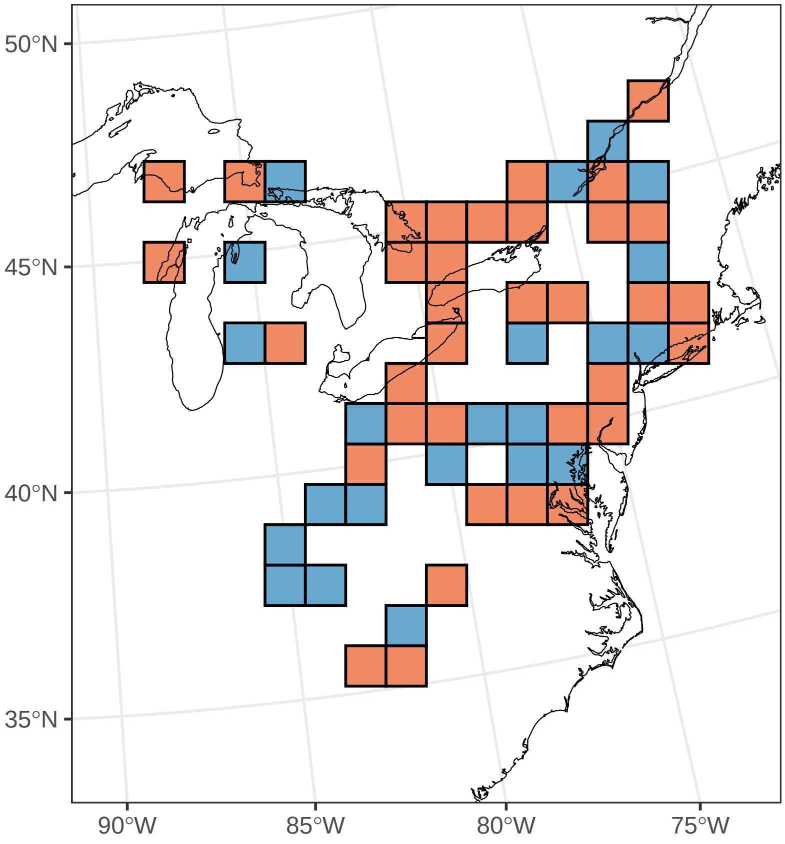

#### ***Viola rostrata***

Ephemerality index: 0.667

n = 24

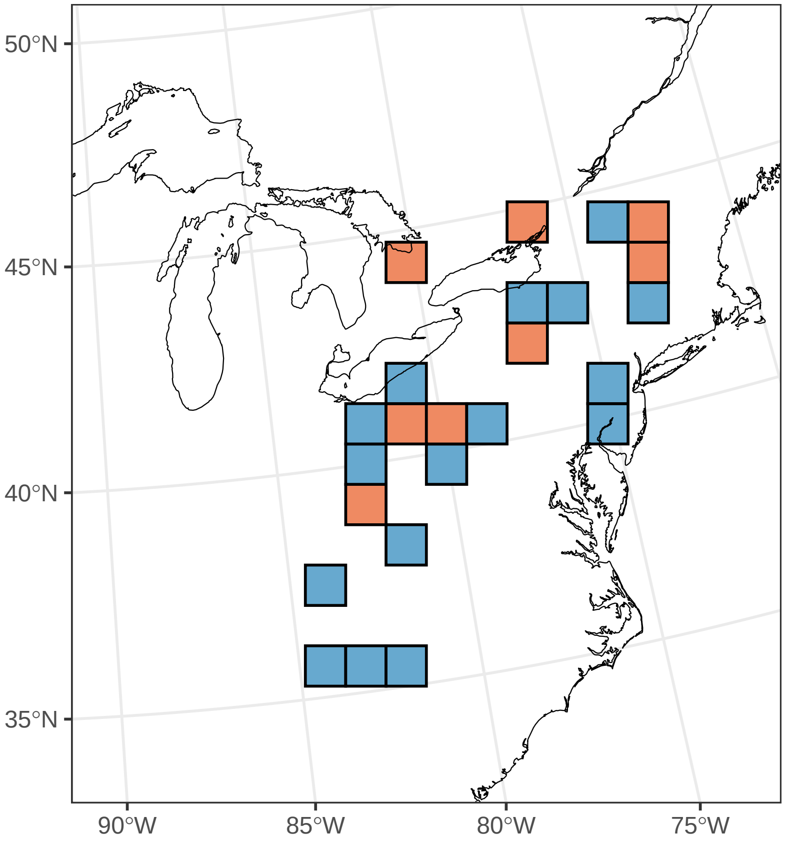

#### ***Viola rotundifolia***

Ephemerality index: 0.05

n = 20

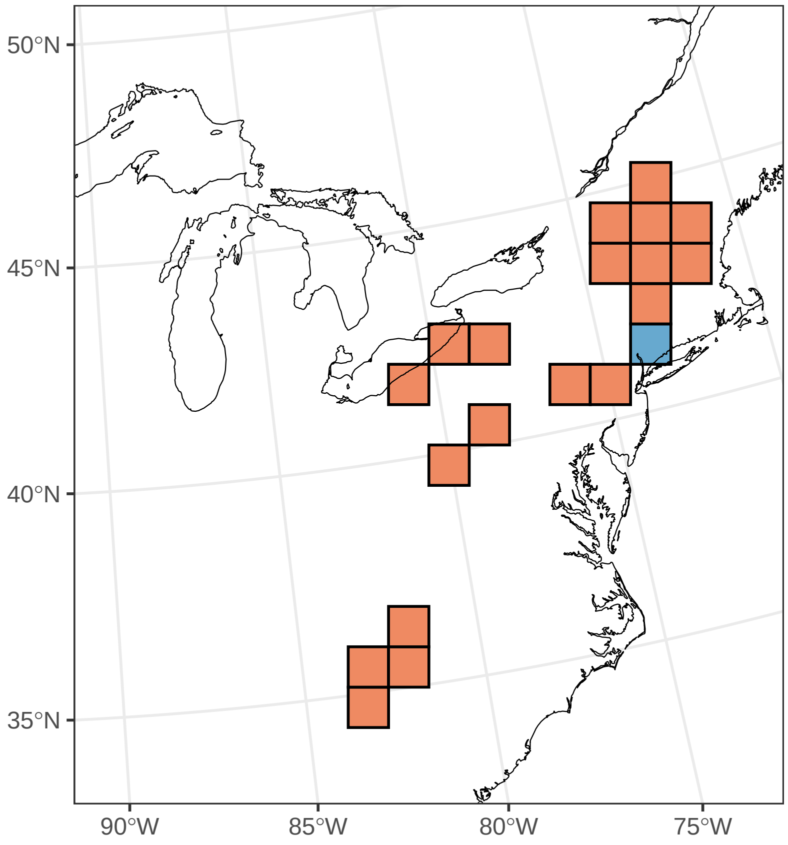

#### ***Viola sororia***

Ephemerality index: 0.273

n = 66

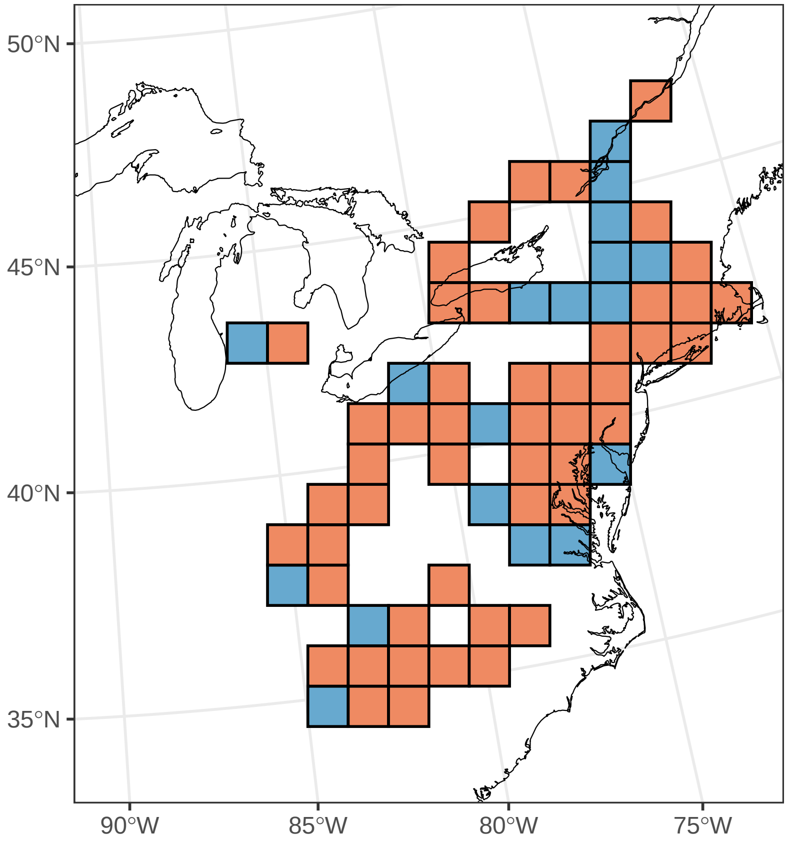

#### ***Viola striata***

Ephemerality index: 0.4

n = 20

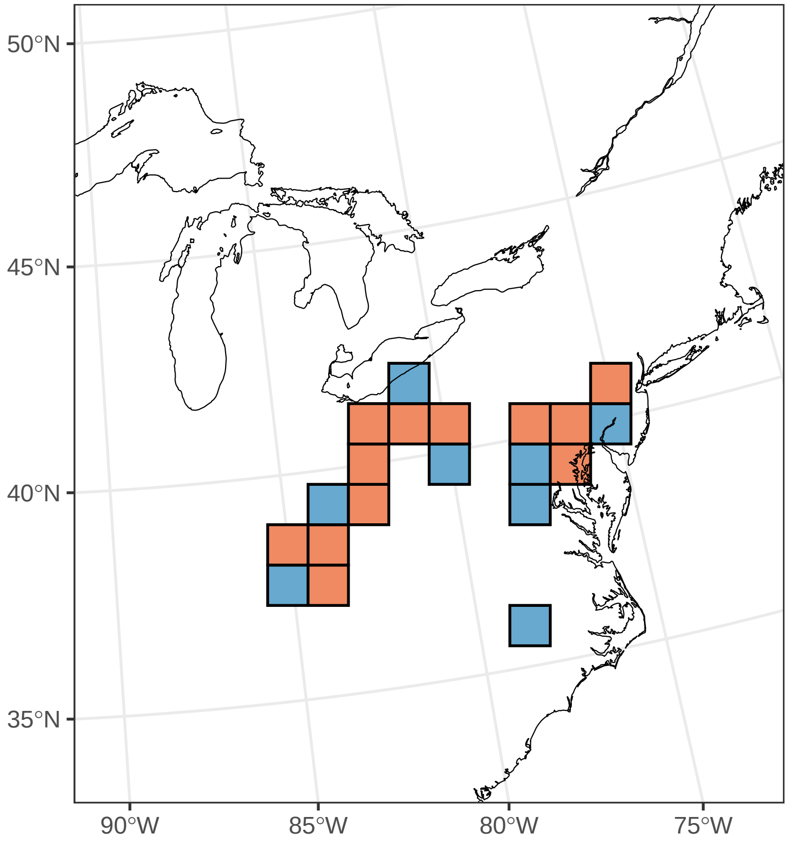

#### ***Youngia japonica***

Ephemerality index: 0.118

n = 17

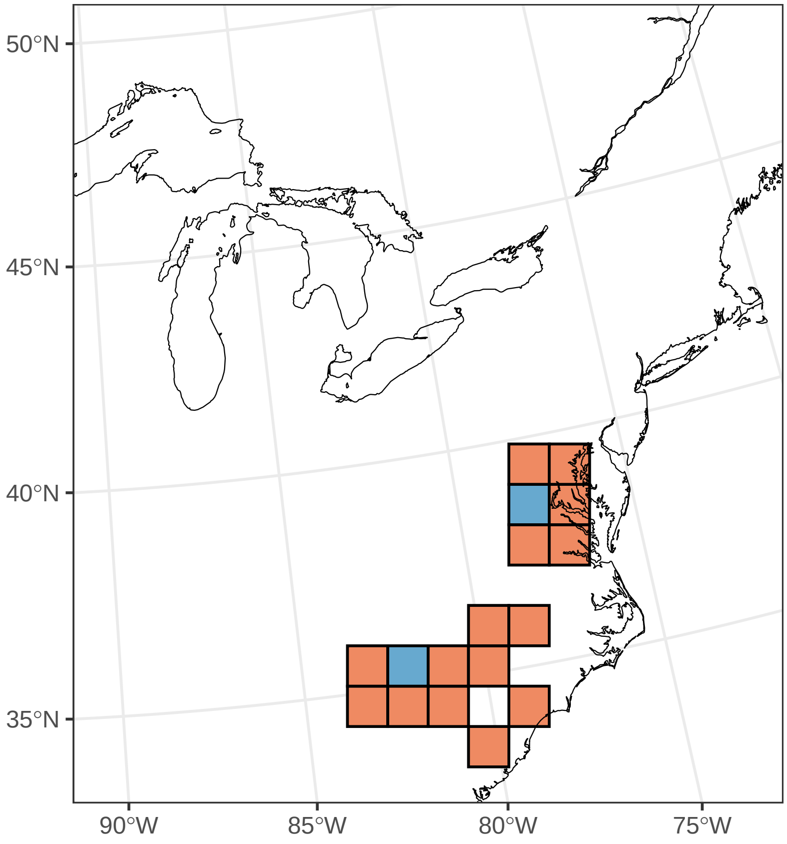

***Zizia aurea***

Ephemerality index: 0.031

n = 32

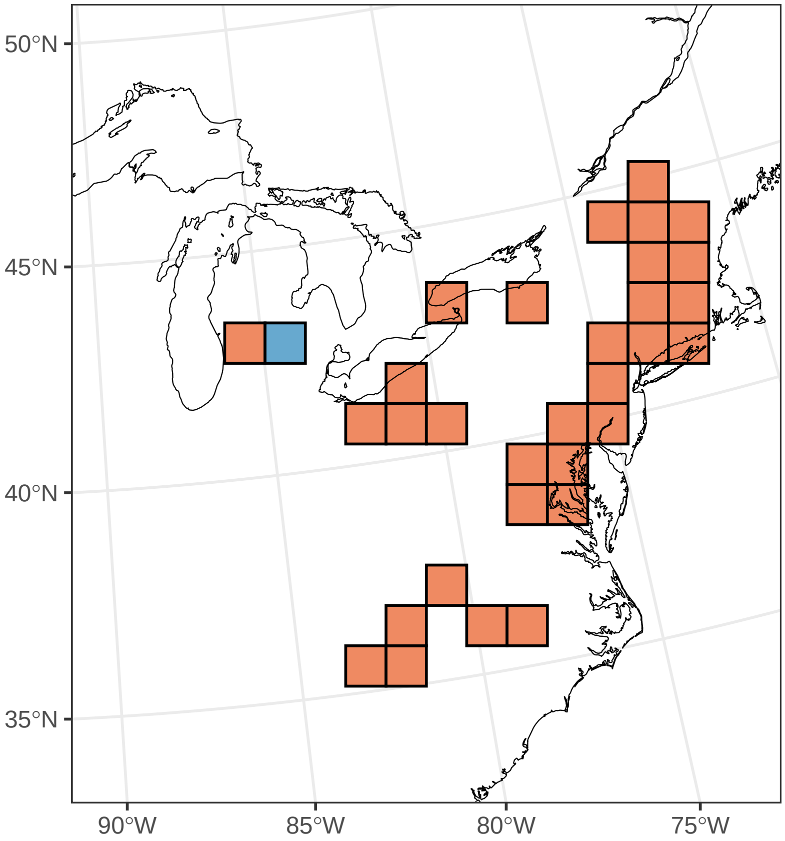
